# Inactivation of p53 drives breast cancer brain metastasis by altering fatty acid metabolism

**DOI:** 10.1101/2023.12.20.572490

**Authors:** Kathrin Laue, Sabina Pozzi, Yael Cohen-Sharir, Tom Winkler, Yonatan Eliezer, Sahar Israeli Dangoor, Alicia I. Leikin-Frenkel, Katharina Lange, Johanna Zerbib, Alessia A. Ricci, Andrea Sacconi, Jean Berthelet, Alexander Schäffer, Wei Shi, Yang Liao, Iris Barshack, Hind Medyouf, Delphine Merino, Giovanni Blandino, Luca Bertero, Ronit Satchi-Fainaro, Uri Ben-David

**Affiliations:** Department of Human Molecular Genetics and Biochemistry, Faculty of Medicine, Tel Aviv University, Tel Aviv 6997801, Israel; Department of Physiology and Pharmacology, Faculty of Medicine, Tel Aviv University, Tel Aviv 6997801, Israel; Department of Molecular Cell Biology, Weizmann Institute of Science, Rehovot 7610001, Israel; Faculty of Medicine, Tel Aviv University, Tel Aviv 6997801, Israel; The Strassburger Lipid Center, Sheba Medical Center, Tel Hashomer, Ramat-Gan, Israel; Pathology Unit, Department of Medical Sciences, University of Turin, Turin, Italy; Translational Oncology Research Unit, Regina Elena National Cancer Institute, Rome, Italy; Olivia Newton-John Cancer Research Institute, Melbourne, Australia; School of Cancer Medicine, La Trobe University, Melbourne, Australia; Georg-Speyer-Haus, Institute for Tumor Biology and Experimental Therapy, Frankfurt am Main, Germany; Institute of Pathology, Sheba Medical Center, Tel HaShomer, Ramat Gan, Israel; Department of Medical Biology, Faculty of Medicine, Dentistry, and Health Science, The University of Melbourne, Parkville, Australia; Immunology Division, The Walter and Eliza Hall Institute of Medical Research, Parkville, Australia; Sagol School of Neurosciences, Tel Aviv University, Tel Aviv 6997801, Israel

## Abstract

Brain metastasis (BM) is a dire prognosis across cancer types. It is largely unknown why some tumors metastasize to the brain whereas others do not. We analyzed genomic and transcriptional data from clinical samples of breast cancer BM (BCBM) and found that nearly all of them carried p53-inactivating genetic alterations through mutations, copy-number loss, or both. Importantly, p53 pathway activity was already perturbed in primary tumors giving rise to BCBM, often by loss of the entire 17p chromosome-arm. This association was recapitulated across other carcinomas. Experimentally, p53 knockout was sufficient to drastically increase BCBM formation and growth *in vivo*, providing a causal link between p53 inactivation and brain tropism. Mechanistically, p53-deficient BC cells exhibited altered lipid metabolism, particularly increased fatty acid (FA) synthesis and uptake, which are characteristic of brain-metastasizing cancer cells. FA metabolism was further enhanced by astrocytes in a p53-dependent manner, as astrocyte-conditioned medium increased FASN, SCD1, and CD36 expression and activity, and enhanced the survival, proliferation and migration of p53-deficient cancer cells. Consequently, these cells were more sensitive than p53-competent cells to FA synthesis inhibitors, in isogenic cell cultures, in BCBM-derived spheroids, and across dozens of BC cell lines. Lastly, a significant association was observed between p53 inactivation, astrocyte infiltration, and SCD1 expression in clinical human BCBM samples. In summary, our study identifies p53 inactivation as a driver of BCBM and potentially of BM in general; suggests a p53-dependent effect of astrocytes on BC cell behavior; and reveals FA metabolism as an underlying, therapeutically-targetable molecular mechanism.

## Introduction

Breast cancer (BC) is the most common malignancy in women worldwide. BC is a molecularly heterogeneous disease, and treatment strategies differ according to molecular subtype. However, advanced BC with distant organ metastases is extremely difficult to treat. BC can metastasize to various organs, a process known as organotropism. The most frequent metastatic sites are the bone (67%), lungs (37%), liver (41%) and brain (13%) (*1*, *2*). Brain metastases (BM) are associated with a particularly bleak prognosis, with increased morbidity and very high mortality rates (*3*). While different tumor subtypes exhibit distinct organotropism patterns, the molecular features that determine metastasis organotropism are poorly understood. Specifically, it remains unclear why some tumors metastasize to the brain, whereas others do not.

*TP53* (coding to the well-known tumor suppressor p53) is the most commonly mutated tumor-suppressor gene in BC, found in 41% of tumors overall, and in 84% of the basal-like tumors (*1*). p53 regulates the cellular response to various types of stress, and is critical to ensure genomic integrity by promoting DNA repair, cell cycle arrest, and apoptosis, among other functions (*4*). In addition to being commonly mutated, *TP53* is commonly perturbed through a copy-number loss of the short arm of chromosome 17 (chr17p), which occurs in 30-45% of tumors (*5–7*). Despite its well-established roles in virtually every step of tumor initiation and progression, the contribution of p53 to BC organotropism has yet to be determined.

The brain microenvironment (BME) is restrictive for cancer cell growth and proliferation. Reactive astrocytes are key components of the BME, and have been shown to promote BM by secreting growth factors, cytokines, and metabolites that enhance the survival and proliferation of metastasizing cells or modulate their interactions with the immune system (*8–12*). The amount of lipids available for cancer cells in the BME is limited in comparison to other tissues (*13*), and a couple of recent studies found that fatty acid synthesis (FAS) was elevated in BC brain metastases (BCBM), rendering BCBM cells sensitive to FAS inhibition (*13*, *14*). Similar results were obtained with acute lymphoblastic leukemia (ALL) cells metastasizing to the central nervous system (CNS)(*15*). Importantly, the BME has been suggested to contribute to increased *de novo* lipid synthesis in BCBM (*13*), but the role of astrocytes in this phenomenon has yet to be studied. Furthermore, until now, it has been unknown whether common genetic alterations, such as p53 inactivation, can modulate the response of brain-metastasizing cells to reactive astrocytes, and/or promote FAS in brain-metastasizing cells.

## Results

### p53 inactivation is ubiquitous in BCBM

*TP53* can be genetically perturbed through point mutations or through its copy-number loss. Biallelic inactivation of *TP53* often involves the loss of one allele through the deletion of one copy of the entire short arm of chromosome 17, on which *TP53* resides (del17p) (*7*). Across three datasets of primary human BC samples (*6*, *16*, *17*), we confirmed that *TP53* was commonly perturbed by point mutations (8%-17%), by copy-number loss (mostly del17p; 43%-55%), or by both (29%-46%) (**fig. S1A**). Monoallelic inactivation of *TP53* was sufficient to induce a significant reduction in its mRNA levels, suggesting its functional importance (**Fig. 1B**). We therefore analyzed the prevalence of *TP53* perturbation across various BC metastatic sites, using clinical data from 1,756 BC patients (mostly ER+ tumors) (*17*). We found >10-fold enrichment of del17p in BM in comparison to all other metastatic sites (**Fig. 1A** and **fig. S1C**; p=2*10^-5^). Extending the analysis to all modes of *TP53* perturbation, we identified a perturbation of at least one allele of *TP53* in 100% of the BM (n=26) – a significant enrichment over the prevalence of *TP53* perturbation in primary tumors (n=477; p=1*10^-4^) and in other metastatic sites (n=722; p=4*10^-4^) (**Fig. 1B**). A strong and significant enrichment was also observed when only biallelic *TP53* inactivation was considered (**Fig. 1B**).

**Figure 1.**
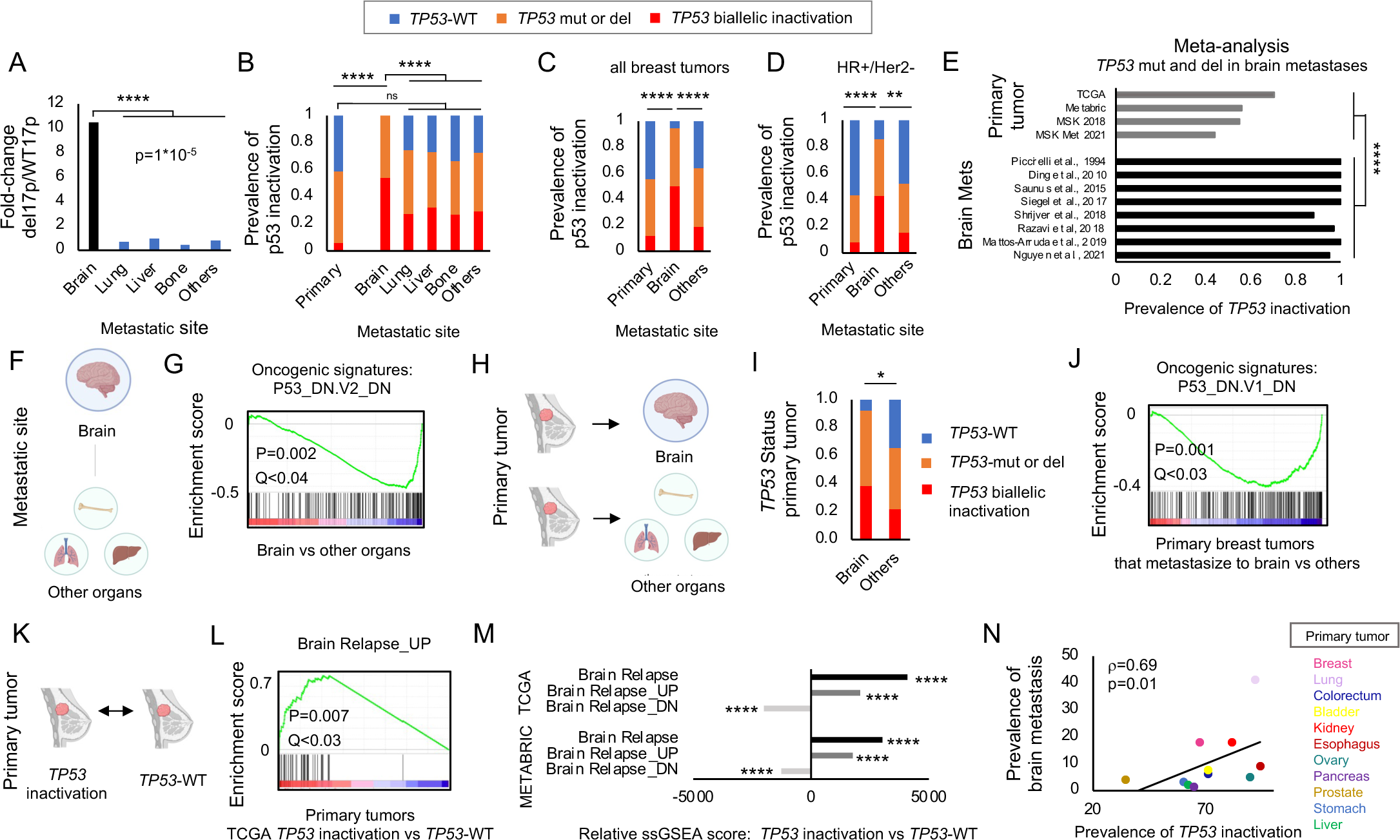
Genomic and transcriptomic analyses of clinical data reveal that *TP53* perturbation is highly enriched in BCBM. **A,** Prevalence of arm-level deletion of chromosome 17p (del17p) in metastases from common metastatic sites, presented as fold-change in del17p over WT17p tumors. Brain metastases showed a >10-fold enrichment of chr 17p deletion over other metastatic sites. Brain compared to other sites, two-tailed Chi-square test, p=2*10^-5^. Brain, n=23; Lung, n=24; Liver, n=135; Bone, n=98; Others, n=268. Data source: Razavi et al., *Cancer Cell* 2018 (*17*). See also fig. S1C. **B-D,** Comparison of the frequency of *TP53*-WT, *TP53* mutation or *TP53* deletion (focal or through del17p), and *TP53*-biallelic inactivation, in primary breast tumors and their metastatic sites. Nearly all brain metastases had genetic alterations of *TP53*, and the prevalence of p53 events significantly exceeded that of all other metastatic sites and primary tumors. No difference was observed between the frequency of p53 events in primary tumors and any other metastatic site except for the brain. Data source: Breast_MSK_2018 data (discovery dataset). Initial findings were validated with data from the larger MSK_Met_2021 cohort. **B**, MSK_2018: *TP53* inactivation vs *TP53*-WT: Brain mets vs primary tumors, Fisher’s exact test p=1*10^-4^; Brain mets vs other metastatic sites (Others), p=4*10^-5^; Primary tumors vs others, p=0.19. *TP53* biallelic inactivation vs *TP53*-WT: Brain vs primary tumors, Fisher’s exact test p=1*10^-4^; Brain vs others, p=1*10^-4^; Primary tumors vs others, p=0.06; Primary n=477, Brain n=26, Lung n=66, Liver n=218, Bone n=116, Others n=322. **C,D,** MSK_Met 2021. **c,** all breast tumors: *TP53* inactivation vs *TP53*-WT: Brain vs primary tumors, Fisher’s exact test p=5.6*10^-13^; Brain vs others, p=8.4*10^-7^, Primary tumors vs others, p= 1.2*10^-6^. Primary n=1351, brain n=38, others n=847. **D,** HR+/HR-tumors. *TP53* inactivation vs *TP53*-WT: Brain vs primary tumors, Fisher’s exact test p=3.4*10^-5^; Brain vs others, p= 0.007; Primary tumors vs others, p= 4.2*10^-6^. Primary n=941, brain n=14, others n=589. For stratification by BC subtype, see fig. S1D. **E,** Meta-analysis of the prevalence of *TP53* genetic perturbation in human BCBM. Analyses included *TP53* mutations, focal deletions and del17p. *TP53* perturbation is significantly more prevalent in brain metastases than in primary breast tumors. One-sided Student’s t-test, p=2*10^-6^. **F,G,** Gene set enrichment analysis (GSEA) comparing brain metastases to other metastatic sites. Transcriptional signatures reflecting p53 pathway activity were significantly lower in brain metastases as compared to metastases of other organs. Data source: GSE14017, Brain vs others, Oncogenic signature P53_DN.V2_DN p<0.002, q=0.04. Brain n=15, others n=14. For a second data set, see fig. S1F,G and Table S1. **H,I,** *TP53* status assessment in primary breast carcinomas with known metastatic sites. The prevalence of both overall and biallelic *TP53* alterations was significantly higher in primary tumors that metastasized to the brain than in those that metastasized to other organs. Data from the Metastatic Breast Cancer Project (MBP). Brain vs others, any *TP53* alteration, one-tailed Fisher’s exact test, p=0.03, TP53 mutation p=0.02 and *TP53* biallelic inactivation p=0.02. Brain n=13, others n=167. Tumors that metastasized to >4 sites were excluded from the analysis. **J,** Gene set enrichment analysis (GSEA) comparing primary tumors that eventually developed BM to those that spread to other organ sites. Transcriptional signatures reflecting p53 pathway activity were significantly lower in primary tumors that would spread to the brain. The Metastatic Breast Cancer Project (MBP), Brain vs Others, oncogenic signatures P53_DN.V1_DN p<0.001. Brain n=7 taken at diagnosis, others n=75. Also see Table S2. **K-M,** Gene set enrichment analysis (GSEA) comparing gene expression signatures of human primary breast carcinomas with *TP53* inactivation to *TP53*-WT primary tumors. **K**, schematic illustration of the comparison. **L,** Enrichment plot demonstrating upregulation of a gene expression signature associated with BM in primary tumors with *TP53* inactivation. *TP53* inactivation vs *TP53*-WT, SMID_BREAST_CANCER_RELAPSE_IN_BRAIN P<0.007. *TP53* inactivation=623, *TP53*-WT n=259. Also see Table S3. **M,** Single-sample gene set enrichment analysis (ssGSEA) of primary BC with or without *TP53* alterations, using data from TCGA and METABRIC datasets. Gene expression profiles associated with BM were increased in primary tumors with *TP53* perturbations. TCGA, SMID_BREAST_CANCER_RELAPSE_IN_BRAIN Student’s t-test p=2.3*10^-24^, _UP p=4.3*10^-23^, _DN p=2.06*10^-23^. METABRIC, SMID_BREAST_CANCER_RELAPSE_IN_BRAIN; Student’s t-test P=5.9*10^-59^, _UP p=3*10^-46^, _DN p=3.1*10^-62^. Table S3. **N,** *TP53* inactivation is correlated with the prevalence of BM across carcinomas. Each dot represents a primary carcinoma type (for lung and kidney, the highest-ranking subtypes were considered). Data source: TCGA and Riihimäki et al. *Cancer Med* 2018 (*36*). Spearman’s correlation rho = 0.69, p =0.01 (one-tailed). Number of tumors that metastasized to the brain from indicated primary tumors: Breast n=4583, lung=10422, colorectum n=1988, bladder n=339, kidney n=1332, esophagus n=206, ovary n=315, pancreas n=116, prostate n=1155, stomach n=235, liver n=123. See also fig. S6.

We validated the strong association between BCBM and *TP53* perturbation in an elaborated clinical dataset of BC metastases (MSK-MET) (*18*), and found that it held true both when all BC subtypes were considered together (**Fig. 1C**) and when each molecular subtype was considered separately (**Fig. 1D** and **fig. S1D**). We then extended the analysis to 6 additional published clinical studies (*18–24*), for which both *TP53* mutation status and *TP53* copy-number status were available, and found that nearly all (114/116) BCBMs had a *TP53* inactivating mutation, copy-number loss, or both (**Fig. 1E**); this high prevalence was significantly higher than the prevalence of these genetic alterations in primary mammary tumors (**Fig. 1E**). Importantly, BCBM prevalence was very similar across various modes of *TP53* perturbation (hotspot mutations, other point mutations, or deletions; **fig. S2A,B**), suggesting that it is indeed p53 inactivation (rather than its gain-of-function (*25*)) that drives the observed enrichment.

We next compared the genome-wide gene expression profiles of BC metastases from various metastatic sites, using gene set enrichment analysis (GSEA) (**Fig. 1F** and **Table S1**) (*26*, *27*). Consistent with the increased prevalence of *TP53* genetic inactivation in BCBM, the transcriptional signature of the p53 pathway was significantly downregulated in BCBMs in comparison to other metastases (p=0.002; **Fig. 1G** and **fig. S1E,F**). We validated these results using RNAseq data from a new clinical cohort of 6 BCBMs and 26 metastases from other organs (**fig. S1G** and **Table S1**). Lastly, we performed RNAseq on 7 metastases obtained from an autopsy of a single BC patient. The transcriptional signature of the p53 pathway was indeed lower in the BM than in the other metastases (**fig. S1H**). Together, these findings reveal an overwhelming enrichment for inactivating *TP53* genetic alterations, resulting in decreased activity of the p53 pathway, in BCBM.

### p53 inactivation in primary BC is associated with brain metastatic capacity

Genetic alterations that are enriched in metastases may emerge *de novo* during the metastatic process, or may pre-exist in the primary tumors that give rise to the metastases (*28–30*). Given that *TP53* is commonly altered already in primary breast tumors (**fig. S1A**), we hypothesized that p53-deficient primary tumors may be more likely to metastasize to the brain. We thus compared the *TP53* status of 180 primary metastatic breast tumors according to their known metastatic sites (using the Metastatic Breast Cancer (MBC) Project; **Fig. 1H**) (*31*, *32*). Primary tumors that would metastasize to the brain (n=13) were significantly more likely to harbor *TP53* genetic alterations in comparison to primary tumors that would metastasize to other organs (n=167; p=0.03 for all genetic alterations, p=0.02 for biallelic genetic alterations; **Fig. 1I**). This association was confirmed in the MSK-MET dataset as well (**fig. S3A**). Consistent with these results, the prevalence of BCBM correlated significantly with the prevalence of *TP53* perturbation in primary tumors across the main BC molecular subtypes (**fig. S3B,C**), and this association was unique for BM, as it was not observed for other metastatic organs, such as bone or lung (**fig. S3D-G**).

Next, we analyzed genome-wide gene expression profiles of tumors from the MBC project. The transcriptional signature of the p53 pathway was significantly downregulated in primary breast tumors that metastasized to the brain in comparison to those that metastasized elsewhere (**Fig. 1J**, **fig. S4A** and **Table S2**). BM-specific downregulation of the p53 pathway was also observed in an independent cohort of 204 primary tumors from BC patients (**fig. S4B, Table S2**) (*33*), as well as in a third cohort of 71 TNBC patients (**fig. S4C**) (*34*), with known metastatic sites. These results reveal reduced gene expression of the p53 pathway in primary breast tumors with capacity for BM.

We next compared the gene expression patterns of *TP53*-WT tumors vs. tumors with inactivation of *TP53* (*TP53*-deficient tumors) in 3 large clinical datasets (TCGA, METABRIC, MBC; **Fig. 1K**). GSEA revealed a strong and significant upregulation of transcriptional signatures associated with brain relapse, as well as a strong and significant downregulation of a transcriptional signature negatively associated with brain relapse, in TP53-deficient tumors (**Fig. 1L,M**, **fig. S4D,E** and **Table S3**); in all datasets, these BM-related signatures ranked highly among differentially-expressed signatures. We validated that these signatures were indeed increased in primary tumors that metastasized to the brain vs. those that metastasized to other organs (**fig. S4F, Table S4**), and that they were independent of the specific mode of p53 inactivation (**fig. S2C,D**). Importantly, *TP53* inactivation was negatively associated with bone-relapse transcriptional signatures (**fig. S4G,H**). Similar enrichments were observed when only tumors with biallelic TP53 inactivation were analyzed (**fig. S4I-N**), and also when each BC molecular subtype was considered separately (**fig. S5**). Together, these results further highlight the strong and specific association of p53 inactivation with BCBM, rather than with metastasis in general.

Overall, our data show that primary BC that metastasize to the brain are enriched for p53 inactivating genetic alterations and downregulate p53 pathway activity, in comparison to tumors that metastasize to other organs. Moreover, primary BC that have inactivated p53 exhibit elevated levels of gene expression signatures associated with BCBM. We thus conclude that p53 inactivation in primary breast tumors is strongly associated with BM potential and may predict BM risk.

### p53 inactivation is associated with BM capacity across carcinomas

With the exception of melanomas, which may commonly metastasize to the brain due to their common developmental origin, brain metastases mostly originate from carcinomas of the breast, lung, colon, and kidney^3^. We therefore set out to investigate whether p53 inactivation might be associated with BM from other primary tumor types. Remarkably, the prevalence of *TP53* genetic perturbation (*35*) was strongly correlated with the prevalence of BM across carcinomas (r=0.69, p=0.01; **Fig. 1N** and **fig. S6A**) (*36*). We next performed a pan-carcinoma analysis of the prevalence of *TP53* genetic alterations across primary tumors and metastatic sites in ∼11,000 patients from the MSK-IMPACT dataset (*37*). Both all and biallelic *TP53* genetic alterations were significantly (p<1*10^-4^) more common in BM than in primary tumors and in other metastatic sites (**fig. S6B**).

Next, we analyzed the association between p53 inactivation and BM capacity across hundreds of human cancer cell lines **(fig. S6C**)(*14*). A pan-carcinoma analysis revealed a significant enrichment for biallelic *TP53* inactivation in cell lines with high vs. low BM potential (p=0.03; **fig. S6D**). Furthermore, GSEA identified strong and significant downregulation of p53 pathway activity in cell lines with high vs. low BM potential (**fig. S6E, Table S2**).

We then focused on lung carcinomas due to their major contribution to BM (*3*). Both all and biallelic *TP53* alterations were significantly more common BM vs. other metastatic sites of non-small cell lung carcinomas (NSCLC) (**fig. S7A**), and the same trend was observed when the analysis was performed within specific molecular subtypes (adenocarcinomas, squamous cell carcinomas, and poorly-differentiated NSCLC; **fig. S7B-D**). We next compared the gene expression profiles of *TP53*-WT tumors vs. tumors with biallelic inactivation of *TP53* (*TP53*-deficient tumors), using the TCGA cohort of lung adenocarcinomas (*35*, *38*). GSEA revealed significant enrichment for brain-relapse (but not bone-relapse) transcriptional signatures in the *TP53*-deficient tumors (**fig. S7E-K**) These results suggest that the association between p53 pathway inactivation in primary tumors and BM capacity exists not only in BC but also in other carcinomas, including in lung cancer.

### Inactivation of p53 in BC cells increases their potential to metastasize to the brain

To examine whether p53 inactivation is causally linked to increased BCBM, we turned to BC cell lines. We first validated that human BC cell lines indeed recapitulate the association that we observed in tumors. Across 23 human BC cell lines (*14*), *TP53* genetic alteration was significantly associated with metastasis to the brain, but not to other organs (lung, liver, kidney, or bone), upon intra-cardiac injection (**Fig. 2A**). A comparison of the BC cell lines with high vs. low BM potential (*14*) (**Fig. 2B**) revealed that all (5/5) brain-metastasizing cell lines had a genetic alteration in *TP53* (p=0.02; **Fig. 2C**). GSEA of RNAseq data from these cell lines confirmed a significant (p=0.008) downregulation of the p53 pathway in the brain-metastasizing cell lines (**Fig. 2D, Table S2**). GSEA of *TP53*-WT vs. *TP53*-deficient cell lines revealed significant upregulation of transcriptional signatures associated with brain-specific relapse (**Fig. S8**). We thus conclude that BC cell lines recapitulate the clinical analysis and make for an appropriate system for functional and mechanistic studies.

**Figure 2.**
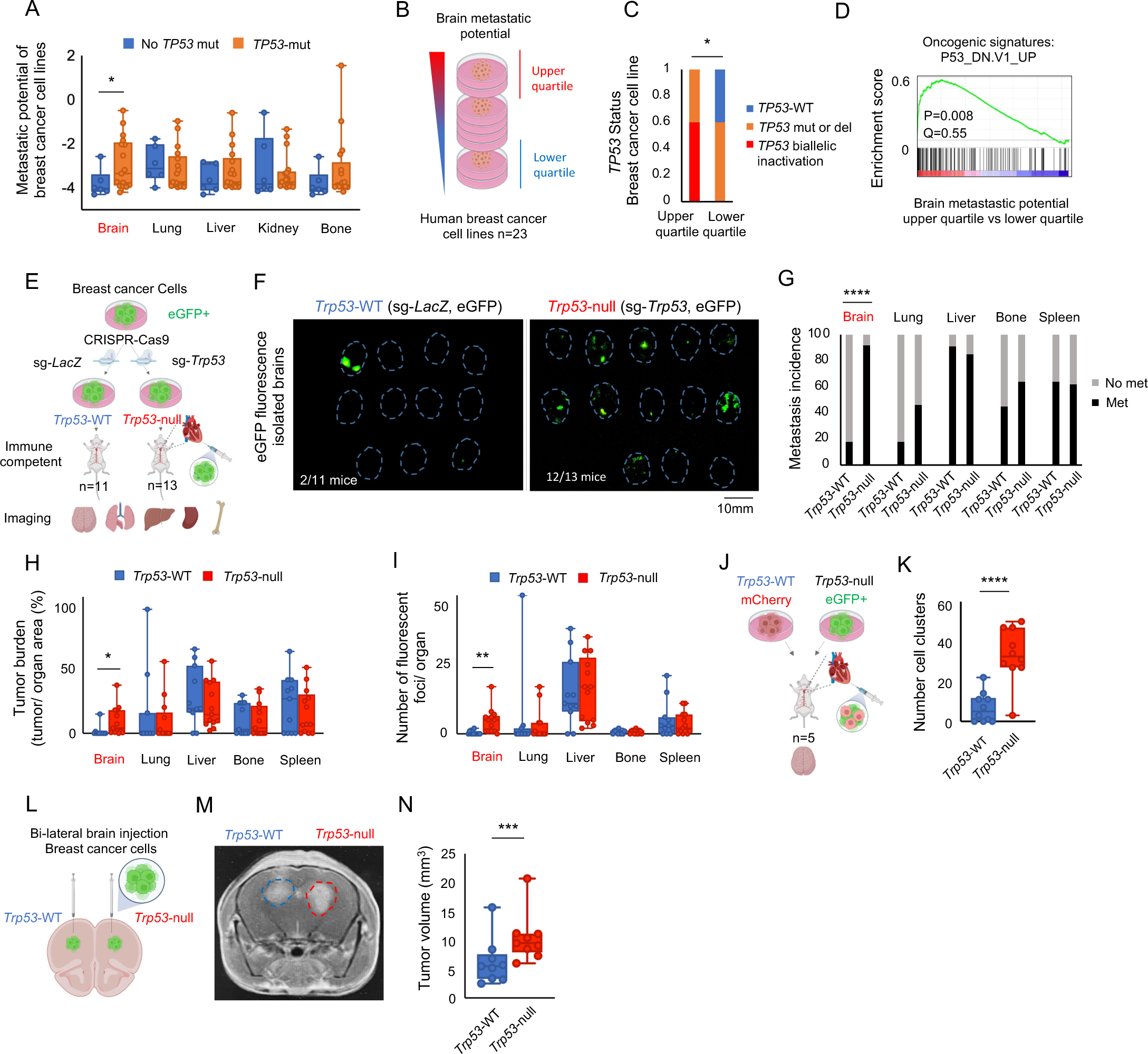
Inactivation of p53 in *TP53*-WT BC cells increases BM and tumor growth in the brain. **A,** Comparison of the effect of *TP53* alterations on the *in vivo* metastatic capacity of human BC cell lines following intra-cardiac transplantation in mice. Shown is the relative metastatic potential for five common metastatic organs in human BC patients: brain, lung, liver, kidney and bone. *TP53* perturbation is significantly associated with elevated BM in xenograft experiments. Data source: Xin et al., *Nature* 2020 (*14*); *TP53-*mut vs *TP53*-WT, metastatic potential to brain, one-tailed Student’s t-test, p=0.05. Lung, p=0.29; Liver, p=0.17; Kidney, p=0.30; Bone, p=0.16; Cell lines with *TP53*-mut n=17, *TP53*-WT n=6. **B,** Human BC cell lines were grouped by their *in vivo* brain metastatic potential, as determined by xenograft experiments of intra-cardiac transplantation. The top (upper 25%) vs. bottom (lower 25%) quartiles of brain metastatic cell lines were used for downstream analyses. **C,** Assessment of *TP53* Status. *TP53* mutations are enriched in breast carcinoma cell lines with high brain metastatic potential. *TP53-*mut vs no *TP53*-mut, one-tailed Fisher’s exact test, p=0.02, *TP53* biallelic inactivation vs no *TP53*-mut, one-tailed Fisher’s exact test, p=0.03. BC cell lines n=23, high brain metastatic potential n=5, low brain metastatic potential n=5. **D,** GSEA analysis of gene expression profiles from BC cell lines with high (top quartile; n=5) vs low (bottom quartile; n=5) metastatic potential reveals an enrichment of expression signatures consistent with downregulation of p53 signaling in highly brain metastatic BC cells. Oncogenic signatures: P53_DN.V1_UP, p=0.008. Table S2. **E,** Experimental work chart of eGFP-labeled EMT6 *Trp53*-WT/*Trp53*-null isogenic cell line generation and intra-cardiac injection of mouse isogenic cells into immune-competent mice. Five organs were collected, and cancer cells were imaged and quantified by fluorescence at endpoint, 11 days after injection. **F,** Green fluorescent images showing the detection of eGFP-labeled *Trp53*-WT and *Trp53*-null cancer cells in dissected brains following intra-cardiac injection. Brains outlined in blue based on brightfield images. See also fig. S10 for fluorescent and brightfield images of all organs. **G,** Fluorescence-based quantification of metastasis incidence by isogenic *Trp53*-WT and *Trp53*-null cells in the brain and in other organs following intra-cardiac injection. *Trp53*-null cells showed a significant increase in brain metastatic prevalence over *Trp53*-WT cells. *Trp53*-null vs *Trp53*-WT, Fisher’s exact test, brain p=5*10^-4^as compared to lung p=0.39, liver p=1, bone p=1, spleen p=1. Mice injected with *Trp53*-null cells: n=13, *Trp53*-WT cells n=11. **H,** Increased tumor burden in the brain, but not in other organs, in mice injected with *Trp53*-null cells in comparison to *Trp53*-WT controls. Tumor burden was defined as tumor area over organ area in %. *Trp53-null* vs *Trp53*-WT: Two-tailed Student’s t-test brain p=0.04, lung p=0.61, liver p=0.42, bone p=0.94, spleen p=0.47. *Trp53*-WT n=11, *Trp53*-null n=13. **I,** Increased number of fluorescent foci in brain, but not in other organs, in mice injected with *Trp53*-null cells when compared to those injected with isogenic *Trp53*-WT controls. Number of fluorescent foci per organ, *Trp*53-null vs *Trp*53-WT, two-tailed Student’s t-test, brain p=0.004, lung p=0.67, liver p=0.79, bone p=0.94, spleen p=0.53. *Trp53*-WT n=11, *Trp53*-null n=13. **J,** Intracardiac injection of isogenic *Trp*53-WT and *Trp*53-null cells now labeled in different colors, into the same mice in a competition setup (competitive intra-cardiac injection assay). **K,** The number of *Trp53*-null cell clusters was significantly higher. Two-tailed Student’s t-test p=5*10^-5^. Mice n=5. **L,** Bilateral intra-cranial injection of mouse isogenic *Trp53*-WT and *Trp53*-null cells into immune-competent mice. **M,** A representative MRI image of a mouse brain 8 days after bilateral intra-cranial injection of the isogenic cells. **N,** MRI-based quantification of tumor volume of isogenic *Trp53*-WT and *Trp53*-null EMT6 BC cells injected into both hemispheres of the same mice. Tumor volume of *Trp53*-null cells was significantly larger than that of *Trp53*-WT cells 7 days after bilateral intra-cranial injection. *Trp53*-null vs *Trp53*-WT, paired two-tailed Student’s t-test p=5.5*10^-4^. Trp53-WT n=9, *Trp53*-null n=9. n=4. See also fig. S12.

We next genetically engineered isogenic mouse and human BC cell lines with and without functional p53 activity by knocking-out *Trp53* / *TP53* in *Trp53*-WT / *TP53*-WT mouse and human BC cell lines, respectively. We chose the mouse BC cell line EMT6 (*39*) and the human BC cell line CAL51 (*40*), as they are known to have a functional activity of the p53 pathway and a limited ability to metastasize to the brain (*14*, *41*). Cells were fluorescently-labeled, and *Trp53*/*TP53* was knocked-out using CRISPR-Cas9 to establish *Trp53-*null EMT6 / *TP53*-null CAL51 cell lines (**fig. S9A**). Sanger sequencing of the targeted locus confirmed the on-target disruption of *Trp53* (**fig. S9B**), RT-qPCR confirmed that the mRNA levels of both *Trp53*/*TP53* and its transcriptional target *CDKN1A* (encoding for p21) were drastically reduced (**fig. S9C,D,F,G**), and western blots showed that p53 protein levels were reduced to non-detectable levels in the knocked-out populations (**fig. S9E,H**). We therefore confirmed the generation of isogenic cell lines that differ only in their p53 status.

We next performed intra-cardiac injection of the isogenic mouse *Trp53*-WT and *Trp53*-null EMT6 cells into immunocompetent mice. Eleven days post-injection, we euthanized the mice, harvested five tissues – brain, lung, liver, spleen, and bone – and assessed the metastatic organotropism by fluorescence microscopy (**Fig. 2E** and **fig. S10A,B**). Remarkably, at this time point, 12 of 13 mice transplanted with *Trp53*-null cells developed BM, in comparison to only 2 of 11 mice transplanted with *Trp53*-WT cells (p=5*10^-4^; **Fig. 2F,G**). This strong enrichment was specific to the brain, as *Trp53* status did not significantly change the metastatic prevalence in any other tissue (although a milder, non-significant increase was observed in lung and bone, consistent with previous reports of a general metastasis-promoting effect for p53 inactivation) (**Fig. 2G** and **fig. S10B**). In addition to increased prevalence, *Trp53*-null cells were also associated with a higher tumor burden and a higher number of cell foci in the brain, but not in other organs (**Fig. 2H,I** and **fig. S10C**). We then performed a competition assay in which we intra-cardially injected mCherry-labeled *Trp53*-WT cells and eGFP-labeled *Trp53*-null cells into the same mice and assessed their presence in the brain after 8 days (**Fig. 2J** and **fig. S11A**). In this experimental setup as well, *Trp53*-null gave rise to BM more frequently than *Trp53*-WT cells (**fig. S11B**), and these metastases were larger and contained more cell foci (**Fig. 2K** and **fig. S11C-G**). These results confirm that p53 inactivation selectively promotes the capacity of BC cells to disseminate into the brain, establishing for the first time a causal link between p53 inactivation and metastatic brain organotropism.

### Inactivation of p53 in BC cells increases their growth in the brain microenvironment

The intra-cardiac injection experiments indicated that BC cells can either colonize the brain more efficiently, or grow better in the brain once they arrive there, or both. To address whether the p53 status affects the growth of the cells in the BME, we transplanted the isogenic cells intra-cranially and followed their growth in the brain. We first performed a bilateral injection of *Trp53*-WT and *Trp53*-null EMT6 cells into the two hemispheres of the same immunocompetent mice (**Fig. 2L**). We assessed tumor growth by MRI, and found a 2-fold increase in the volume of tumors originating from *Trp53*-null vs. *Trp53*-WT cells on day 8 post-injection (p=3*10^-4^; **Fig. 2M,N**). Next, we performed unilateral injections (**fig. S12A**) in order to assess the effect of p53 status on tumor progression and animal survival. Consistent with our previous findings, *Trp53*-null cells gave rise to significantly larger tumors (**fig. S12B**), which progressed more quickly (**fig. S12C**), resulting in a trend towards worse survival. Intra-cranial transplantation of the human isogenic CAL51 cells into immune-deficient mice (**fig. S12D**) recapitulated these results: *TP53*-null cells gave rise to larger tumors (p=4*10^-4^; **fig. S12E,F)** and progressed faster than *TP53*-WT-derived tumors (**fig. S12G,H**). We conclude that p53 inactivation promotes the ability of BC cells to thrive in the BME, independently of the immune microenvironment.

### Astrocytes promote viability, proliferation, and migration of BC cells in a p53-dependent manner

Tumor growth involves complex interactions of cancer cells with cells in their microenvironment (*42*). We performed immunofluorescence staining of slides obtained from the cancer cells that grew in the mouse brain, in order to compare the abundance of 3 main brain cell types between *Trp53*-WT and *Trp53*-null tumors. Whereas no difference was observed in the abundance of activated microglia (Iba1+) cells and endothelial (CD31+) cells, activated astrocyte (GFAP+) cells were significantly more abundant in the *Trp53*-null tumors (p=0.001; **Fig. 3A,B** and **fig. S13A,B**). Astrocytes, the most abundant cell type of the brain, have been shown to enhance the survival, motility, and proliferation of metastatic cancer cells (*10*, *11*, *43–46*). The increased astrocyte infiltration into *Trp53*-null tumors suggests that they may affect tumor behavior in a p53-dependent manner.

**Figure 3.**
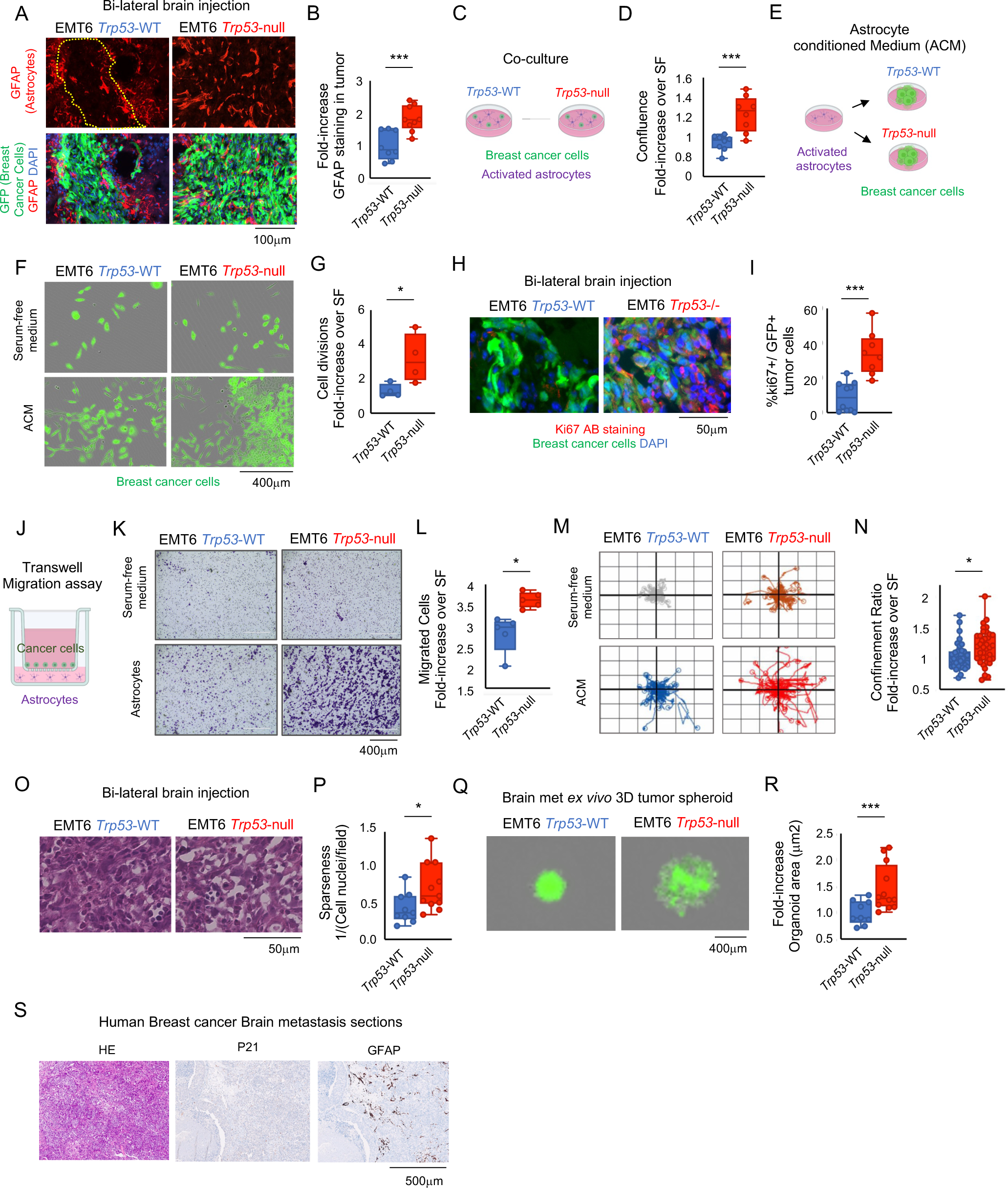
Astrocytes infiltrate p53-null brain metastases and promote the survival, proliferation and migration of p53-null BC cells in the brain. **A,** Mouse brain tumor sections stained with antibodies against astrocytes in red (GFAP) and against BC cells in green (GFP) following intra-cranial brain injection of isogenic *Trp53*-WT and *Trp53*-null EMT6 cells. **B,** Fluorescence-based quantification of astrocytes in the EMT6-derived tumors. GFAP staining intensity was significantly stronger in tumors growing from *Trp53*-null BC cells in comparison to those from *Trp53*-WT cells within the same brain. Data are shown as fold-increase in staining intensity over *Trp53*-WT, two-tailed Student’s t-test p=0.001. *Trp53*-WT n=8, *Trp53*-null n=9. **C,** Co-culture of activated astrocytes with either *Trp53*-WT or *Trp53*-null isogenic BC cells. **D,** Quantification of the effect of astrocytes on the growth of BC cells. *Trp53*-null cells grow better in the presence of astrocytes than their isogenic *Trp53*-WT controls. Confluence is quantified as growth increase fold-change in the presence of astrocytes over SFM after 3 days of co-culture, two-tailed Student’s t-test p=0.001. *Trp53*-WT n=8, *Trp53*-null n=8. n=3. **E,** Culture of *Trp53*-WT or *Trp53*-WT BC cells in the presence of astrocyte-conditioned medium (ACM). **F,** Representative images of *Trp53*-WT and *Trp53*-null EMT6 cells grown in SFM or in ACM. *Trp53-null* cells cultured in ACM had a significant growth advantage over their isogenic *Trp*53-WT and serum-free controls. **G,** Live-imaging-based quantification of the effect of ACM on the growth of BC cells. *Trp53*-null cells grow better in the presence of ACM than their isogenic *Trp53*-WT controls. Cell divisions quantified as growth increase fold-change in ACM over SFM. Two-tailed Student’s t-test p=0.04. *Trp53*-WT n=4, *Trp53*-null n=4. See also fig. S13 and S14. **H,** Mouse brain tumor sections stained with an antibody against proliferating cells (Red, Ki67) and against BC cells (Green, GFP). Shown are representative images 6 days after intra-cranial injection. **I,** Fluorescence-based quantification of ki67+ cells in the EMT6-derived tumors. *Trp53*-null cells were significantly more frequently positive for Ki67 in comparison to *Trp53*-WT cells. Quantification shown as % ki67-positive cells/GFP-positive tumor cells 6 days after injection, p=3*10^-4^, *Trp53*-WT n=3, *Trp53*-null n=3. n=2. **J,** Transwell migration assay to assess the migration of BC cells towards astrocytes. BC cells were placed on a porous filter, which was large enough for the cells to pass through. This chamber was then placed on a well that contained activated astrocytes. **K,** Representative images of *Trp53*-WT and *Trp53*-null BC cells, stained in purple, showing their migration through the filter towards the lower chamber, which contains either SFM or astrocytes. **L,** Quantification of cell migration in the transwell assay after 20h of co-culture of BC cells and astrocytes. The effect of astrocytes on cell migration was significantly stronger in *Trp53-*null cells. Quantification of migrated cells is shown as the fold-increase of migrating cells in astrocytes over serum-free conditions, two-tailed Student’s t-test p=0.03, n=4. See also fig.S14. **M,** Representative live cell imaging-based tracking of cell movement of individual BC cells. Displacement plots show that ACM stimulated cell movement both in *Trp53*-WT and in *Trp53*-null cells, with a larger effect on *Trp53*-null cells. **N,** Live-imaging-based quantification of the effect of ACM on the confinement ratio (degree of cell motility) of *Trp53*-WT and *Trp53*-null cells. Confinement ratio shown as fold-increase in ACM over SFM, Two-tailed student’s t-test, p=0.02. Each data point represents a confinement measurement. **O,** Haematoxylin-Eosin staining of mouse brain tumor sections after bilateral intra-cranial injection of *Trp53*-WT and *Trp53*-null BC cells. **P,** Quantification of cellular density (defined as the number of cell nuclei per field) based on the H&E staining. *Trp53*-null-derived lesions were les compact and more sparse than *Trp53*-WT-derived lesions. Two-sided t-test, p=0.04. **Q,** Representative images of *ex vivo* 3D tumor spheroids derived from *Trp53*-WT and *Trp53*-null EMT6 cells after bilateral intra-cranial injection into mouse brains. Spheroids derived from *Trp53*-null intracranial tumors grew better and had a less compact, more widespread appearance in comparison to their isogenic *Trp53*-WT controls. **R,** Quantification of spheroid size (µm^2^) from *Trp53*-WT- and *Trp53*-null-derived tumors. Data are shown as size increase fold-change in full medium relative to SFM, two-tailed Student’s t-test p=0.001, *Trp53*-WT n=9, *Trp53*-null n=12. **S,** Human BCBM sections stained with Haematoxylin-Eosin (H&E) or with antibodies labeling Astrocytes (GFAP) and p21 (a major downstream effector of p53 pathway activity). Tumor tissue was infiltrated by astrocytes in samples with low to absent p21 staining. See also fig. S15 and Table S5.

To test this hypothesis, we first co-cultured the isogenic EMT6 cells in the presence of lipopolysaccharide (LPS)-activated primary mouse astrocytes (**Fig. 3C**). Astrocytes promoted the proliferation of *Trp53*-null cancer cells significantly more than that of *Trp53*-WT cells (p=0.001; **Fig. 3D** and **fig. S13C**). To determine whether this effect required physical contact between the two cell types, or was mediated by factors secreted by the astrocytes, we next cultured the isogenic cells in an astrocyte-conditioned medium (ACM) (**Fig 3E**). ACM promoted the proliferation of *Trp53*-null cancer cells significantly more than that of *Trp53*-WT cells (p=0.04; **Fig. 3F,G** and **fig. S13D**). In addition, ACM significantly decreased apoptosis (by ∼30%-70%) in the BC cells, and this protective effect was stronger in the *Trp53*-null cells (**fig. S13E**). To test whether *Trp53*-null cells also proliferated faster *in vivo*, we stained slides from the mouse brain lesions for the proliferation marker Ki67. Indeed, *Trp53*-null BC cells proliferated significantly faster than *Trp53*-WT BC cells (p=3*10^-4^; **Fig. 3H,I**), and this *in vivo* proliferation difference was much stronger than the *in vitro* proliferation difference under various cell culture conditions (**fig. S13C,D** and **fig. S16E**).

We next examined whether astrocytes also affected the migration of the isogenic cells in a p53-dependent manner. Both astrocytes and ACM enhanced the migration of *Trp53*-null cells significantly more than that of *Trp53*-WT cells in a transwell migration assay (p=0.03 for astrocytes and p=0.005 for ACM; **Fig. 3J-L** and **fig. S13F,G**). Moreover, live cell imaging showed that ACM increased spontaneous cell motility to a much larger extent in *Trp53*-null cells in comparison to *Trp53*-WT cells (p=0.02; **Fig. 3M,N** and **fig. S13H**). In line with these results, H&E staining of slides from the mouse brains showed that *Trp53*-null-derived tumors were less compact and more sparse than *Trp53*-WT-derived tumors (p=0.02; **Fig. 3O,P**). Lastly, we generated 3D spheroids from the isogenic brain tumors, and followed the spheroid growth. Spheroids from *Trp53*-null tumors were larger and sparser than those from *Trp53*-WT tumors (p=0.001; **Fig. 3Q,R**). We conclude that p53-inactivated mouse BC cells survive, proliferate, and migrate faster in the presence of activated astrocytes, and suggest that factors that are secreted by astrocytes mediate, at least partially, these phenotypes.

To test the relevance of these findings to human cancer, we repeated the experiments with the isogenic human BC cells (CAL51). We co-cultured the isogenic human cells with starvation-activated primary human astrocytes, and found that the human astrocytes indeed promoted cell proliferation in a p53-dependent manner (**Fig. 14A**). Moreover, human ACM promoted cell proliferation of *TP53*-null cells significantly more than that of *TP53*-WT cells (p=9*10^-7^ at 24h; **fig. S14B,C**). In addition, astrocytes and ACM had a p53-dependent protective effect on cell viability: *TP53*-null cells exhibited improved survival and reduced cell death, as evaluated by live imaging and by flow cytometry using Sytox green and Annexin-V staining (**fig. S14D-H**). Similar to our observations with the mouse cells, *TP53*-null CAL51 cells migrated faster in the presence of astrocytes or ACM in comparison to *TP53*-WT cells (p=2*10^-5^ and p=5*10^-8^, respectively; **fig. S14I-M**). Live cell imaging also revealed that ACM preferentially increased the motility of *TP53*-null cells (**fig. S14N-P**). We conclude that astrocytes, and/or astrocyte-secreted factors, promote various metastatic behaviors in a p53-dependent manner in both mouse and human BCBM.

Finally, to ask whether astrocyte infiltration is associated with BCBM in human patients, we quantified p53, p21, and GFAP protein expression levels by IHC in 17 clinical BCBM samples. All samples presented altered expression of p53 and low levels of p21, in line with p53 inactivation (**Fig. 3S**, **fig. S15** and **Table S5**), and consistent with our genomic analyses of clinical human data. Moreover, astrocyte infiltration was evident in most of the tumor samples (**Fig. 3S**, **fig. S15,** and **Table S5**), consistent with our observations from the mouse tumors. Together, these results link p53 status to the BCBM-promoting activity of astrocytes.

### p53 inactivation increases FA uptake through CD36, thereby promoting cell proliferation and migration

We next set out to explore the molecular mechanism(s) underlying the ability of p53-inactivated cells to thrive in the BME. We first compared the gene expression profiles between clinical samples of BCBM and metastases from other organs. In three independent cohorts, BM exhibited upregulation of lipid metabolism, including FA synthesis, uptake, and activation (**fig. S16A,B** and **Table S6 and S7**). GSEA of BM vs. other metastases from the same patient also confirmed the upregulation of these pathways in the brain (**fig. S16C**). A comparison of BM to their corresponding primary BC also revealed upregulation of these pathways in the metastases (**fig. S16D** and **Table S6 and S8**). These results are in line with previous reports of altered lipid metabolism in BM (*13–15*). Intriguingly, we found that the same pathways of lipid metabolism were also upregulated in primary human BC that would metastasize to the brain in comparison to those that would metastasize elsewhere (**Fig. 4A** and **Table S6 and S9**), suggesting a cell-autonomous component to this metabolic difference. Importantly, the transcriptional signatures of lipid metabolism were elevated also when comparing *TP53*-perturbed vs. *TP53*-WT human primary BC (**Fig. 4B** and **Table S6 and S10**). Moreover, high metastatic capacity to the brain was associated with elevated lipid metabolism across human BC cell lines (**Fig. 4C** and **Table S6 and S9**), and the same pathways were also transcriptionally upregulated upon *TP53* knockout in CAL51 cells (**Fig. 4D, Table S6 and S11**). These results suggest that p53 inactivation might be associated with metabolic changes that promote BCBM.

**Figure 4.**
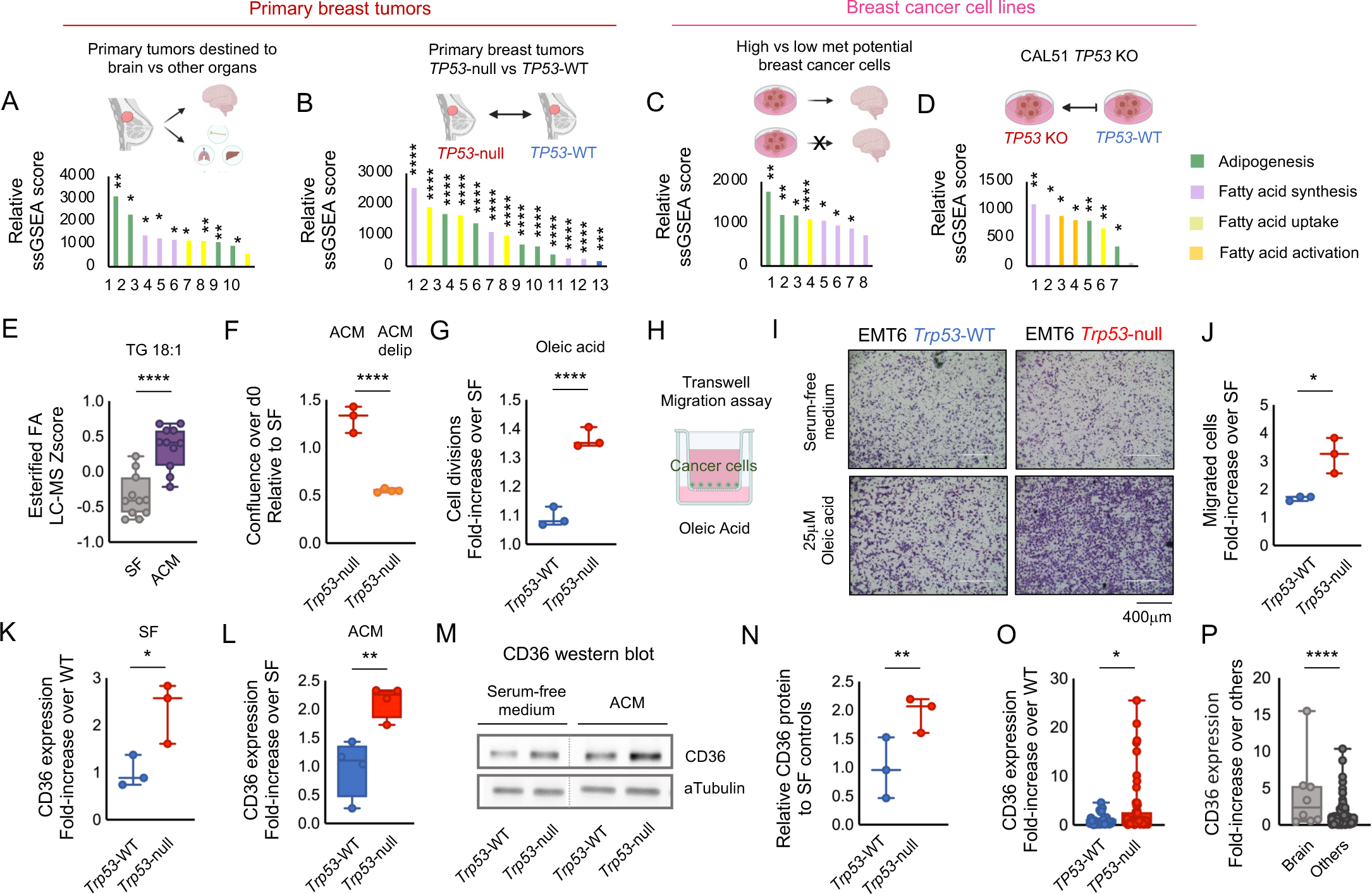
Increased fatty acid uptake through CD36 in p53-null BC cells promotes tumor cell proliferation and migration. **A-B,** Single-sample gene set enrichment analysis (ssGSEA) shows elevated gene expression signatures related to adipogenesis, FA synthesis, FA uptake, and FA activation in: **A,** primary tumors that metastasized to the brain compared to those that metastasized to other organs; **B,** primary tumors that carried biallelic *TP53* inactivation compared to tumors that were *TP53*-WT; **C,** BC cell lines with high brain metastatic potential in comparison to those with low brain metastatic potential; and **D,** human BC cells in which p53 was knocked-down in comparison to their control cells. Also see Tables S6-S11. **E,** Metabolomics quantification of oleic acid-containing triglycerides shows a significant enrichment in ACM. Two-tailed Student’s t-test p=2.3*10^-5^. SFM n=3, ACM n=3. OA-containing triglycerides n=11. Also see Table S12. **F,** Comparison of the proliferation of *Trp53*-null BC cells grown in ACM or delipidated ACM. Delipidation significantly reduced the proliferation-promoting effect of ACM, suggesting that lipids mediate this effect (at least in part). Proliferation shown as confluence relative to SFM after 3 days in culture. *Trp53*-null ACM vs *Trp53*-null delipidated, two-sided Student’s t-test p=1*10^-4^. **G,** Comparison of the effect of oleic acid (OA; 10μM) supplementation on the growth of *Trp53*-WT and *Trp53*-null BC cells. The proliferation-promoting effect of OA was significantly higher in *Trp53*-null cells in comparison to *Trp53*-WT cells. Shown as cell divisions in OA-supplemented over SF medium. One-sided Student’s t-test p=5*10^-4^; *Trp53*-WT n=3, *Trp53*-null n=3. **H,** Transwell migration assay to assess the migration of BC cells towards OA-containing SFM (25μM). **I,** Representative images of *Trp53*-WT or *Trp53*-null BC cells, stained in purple, showing their migration through the filter towards the lower chamber, which contains SFM with or without OA (25μM). **J,** Quantification of cell migration in the transwell assay at 20h. OA increased cell migration of both *Trp53*-WT and *Trp53*-null cells; its effect on *Trp53*-null cells was much more significant. Quantification of migrated cells is shown as the fold-increase of migrating cells in OA over serum-free conditions, two-sided Student’s t-test p=0.01. *Trp53*-WT n=3*, Trp53*-null n=3. **K,** Analysis of *CD36* mRNA expression by RT-qPCR. The mRNA expression levels of the fatty acid transporter CD36 were significantly higher in *Trp53*-null cells relative to *Trp53*-WT controls cultured in SFM. Shown as relative *CD36* expression levels fold-change over the average expression of *Trp53*-WT cells. Two-sided Student’s t-test p=0.03, *Trp53*-WT n=3, *Trp53-*null, n=3. **L,** Analysis of *CD36* mRNA expression by RT-qPCR. The mRNA expression levels of the fatty acid transporter CD36 were significantly higher in ACM-exposed *Trp53*-null cells relative to *Trp53*-WT controls. Shown as relative *CD36* expression levels fold-change over SFM, two-sided Student’s t-test p=0.007, *Trp53*-WT n=4, *Trp53*-null, n=4. **M,** Western blot CD36 protein quantification in isogenic *Trp53*-WT and *Trp53*-null cells cultured in SFM or in ACM. CD36 protein levels were moderately higher in *Trp53*-null cells cultured in SFM, and ACM exposure further elevated CD36 protein levels. **N,** Quantification of CD36 protein levels. The effect of ACM on CD36 protein expression was significantly stronger in *Trp53*-null cells in comparison to *Trp53*-WT cells. CD36 protein levels are shown as fold-change of ACM over SFM conditions. One-sided Student’s t-test, p=0.01, n=3. **O,** Comparison of CD36 mRNA expression levels between *TP53*-WT and *TP53*-null human primary breast carcinomas of the Luminal B molecular subtype, based on TCGA data. *TP53*-null tumors exhibited higher *CD36* mRNA expression levels in comparison to *TP53*-WT tumors. Shown as fold-increase *CD36* expression relative to the average expression of *TP53*-WT tumors. One-sided Student’s t-test p=0.02, *TP53*-WT n=30, *TP53*-null n=56. See also fig. S17D. **P,** Comparison of *CD36* mRNA expression levels between human primary breast carcinomas that metastasized to the brain and those that metastasized to other metastatic sites. Human primary tumors that metastasized to the brain exhibited increased *CD36* mRNA levels. Data source: GSE12276, One-sided Student’s t-test p=3*10^-7^, brain-metastasizing primary tumors n=8, primary tumors metastasizing to other organs n=188.

Lipid availability has been shown to be restrictive for cancer cell growth in the brain (*13–15*, *47–49*). We therefore hypothesized that astrocytes might support BC cells by secreting key FAs, and that the uptake of these FAs might be p53-dependent. To identify lipids that are secreted by astrocytes, we performed liquid chromatography-mass spectrometry (LC-MS) of serum-free medium (SFM) and of ACM. We found a significant increase in oleic acid (OA; 18:1)-containing triglycerides in ACM (p=2.3*10^-5^; **Fig. 4E** and **Table S12**), in line with previous reports showing high OA content in astrocytes (*12*, *50–52*). We then assessed the proliferation of isogenic *Trp53*-WT and *Trp53*-null EMT6 cells under lipid starvation, and found that p53 inactivation significantly improved cell proliferation under these conditions (p=3*10^-4^; **fig. S16E,F**). Delipidation of ACM largely prevented its proliferation-promoting effect (**Fig. 4F**), confirming the important role of lipids in this effect. Moreover, supplementation of OA to SFM preferentially increased the proliferation of *Trp53*-null cells (p=5*10^-4^; **Fig. 4G** and **fig. S16G**). Importantly, OA supplementation phenocopied not only the p53-dependent effect of ACM on proliferation, but also its differential effect on cell migration (**Fig. 4H-J**) and spontaneous cell motility (**fig. S16H**). These results demonstrate that astrocyte-secreted OA indeed affects breast cancer cells in a p53-dependent manner.

CD36 is an FA transporter that serves as the main transporter of FA into human cells (*53*). CD36 has been previously shown to promote metastasis, and to be induced by unsaturated FA such as OA (*51*, *54–57*). We therefore tested whether the upregulation of CD36 in p53-inactivated cells could explain their increased response to OA. We measured CD36 mRNA and protein levels by RT-qPCR and western Blot, respectively, and found that they were already elevated in *Trp53*-null vs. *Trp53*-WT under serum-free culture conditions (p=0.05 and p=0.02 for mRNA and protein expression levels, respectively; **Fig. 4K,M**). Importantly, CD36 levels were further upregulated when cells were cultured in ACM, and this upregulation was much more pronounced in *Trp53*-null cells (p=0.007 and p=0.01 for mRNA and protein expression levels, respectively; **Fig. 4L-N**). A similar preferential upregulation of CD36 was observed in ACM-cultured *TP53*-null vs. *TP53*-WT human CAL51 cells (**fig. S17A-C**). We then analyzed gene expression from clinical BC samples and found that CD36 expression was higher in *TP53*-null vs. *TP53*-WT primary tumors of the luminal molecular subtypes (p=0.05 and p=0.02 for LumA and LumB subtypes, respectively; **Fig. 4O** and **fig. S17C**). Furthermore, CD36 expression was significantly higher in primary tumors that would metastasize to the brain in comparison to those that would metastasize to other organs (p=3*10^-7^; **Fig. 4P**). Together, our findings reveal that p53 inactivation increases, via CD36 upregulation, the ability of BC cells to uptake OA (and, potentially, other FAs), which is made available to the cells by the surrounding activated astrocytes, thereby contributing to their improved proliferation and migration in the BME.

### p53 inactivation increases FAS, enabling cancer cells to utilize FAS-required metabolites secreted by astrocytes

The FAs in the BME cannot support the entire demand of cancer cells for lipids, making metastatic cancer cells dependent on FAS (*13–15*, *58*). Indeed, SREBP1, FASN and SCD1 have been recently implicated in metastasis from BC and leukemia to the CNS (*14*, *15*). SCD1 catalyzes the rate-limiting step of FAS, and is crucial for the synthesis of mono-unsaturated fatty acids (MUFAs), which then metabolize into various FAs that are required to support rapid cell proliferation (*59*). In line with the importance of FAS for BM, we found elevated SCD1 expression in BMs compared to their matched primary tumors (**fig. S18A**) and in BCBMs in comparison to metastases in other organs (**fig. S18B,C**), both in a published dataset (*27*) and in our own cohort.

p53 inactivation was previously suggested to increase FAS in adipocytes (*60*), prompting us to unravel the potential link between p53 and FAS in the context of BCBM. SCD1 mRNA and protein levels were upregulated in *TP53*-null vs. *TP53*-WT primary BCs, in both TCGA (p=2.2*10^-5^ and p=3*10^-4^, for mRNA and protein levels, respectively; **Fig. 5A,B**) and METABRIC (p=5*10^-4^ for mRNA levels; **fig. S18D**) datasets. Moreover, SCD1 gene expression levels were negatively correlated with the activity of the p53 pathway in primary human BCs that metastasized to the brain (Spearman’s rho=-0.62, p=0.01, GSE125989(*61*), **Fig. 5C)**. FASN expression was similarly elevated in *TP53*-null vs. *TP53*-WT primary BCs (Luminal B subtype, p=0.04 and p=0.002, for TCGA and METABRIC datasets, respectively; **fig. S18E)**, and significant correlations were observed between FASN expression and p53 pathway activity (**fig. S18F**). These results confirm a negative association between p53 activity and FAS in human cancer.

**Figure 5.**
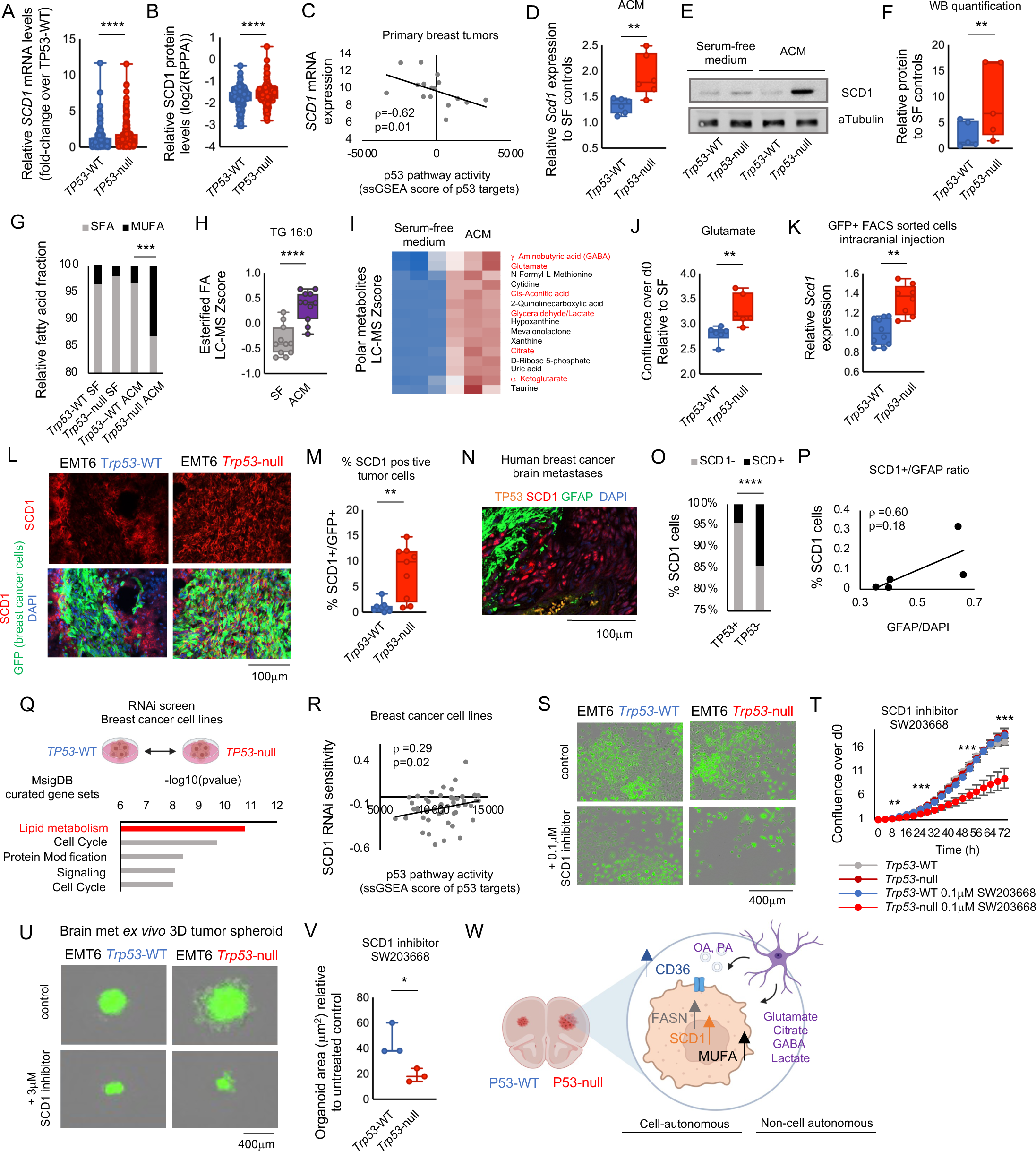
Increased fatty acid synthesis through SCD1 in *p53-*null BC cells is further stimulated by astrocytes and constitutes a cellular vulnerability. **A,** Comparison of *SCD1* mRNA expression levels between *TP53*-WT and *TP53*-null human primary breast tumors, based on TCGA data. Two-sided Student’s t-test p=2.2*10^-5^. *TP53*-WT n=259, *TP53*-null n=302. **B,** Comparison of SCD1 protein expression levels between *TP53*-WT and *TP53*-null human primary breast tumors, based on RPPA data from TCGA. One-sided Student’s t-test p=3*10^-4^. *TP53*-WT n=199, *TP53*-null n=250. **C,** The correlation between the p53 pathway activity score and the mRNA expression levels of *SCD1* across primary human breast tumors that metastasized to the brain (GSE125989). Low p53 pathway activity is associated with high *SCD1* mRNA expression levels. p53 pathway activity was evaluated by calculating a ssGSEA score for the MSigDBgene signature ‘KANNAN_p53_targets’. Spearman’s rho=-0.62, p=0.01. n=16 tumors. **D,** Analysis of *Scd1* mRNA expression by RT-qPCR. The mRNA expression levels of *Scd1* were significantly higher in ACM-exposed *Trp53*-null cells relative to *Trp53*-WT controls. Shown as relative *Scd1* expression levels fold-change over SFM. One-sided Student’s t-test p=0.004. *Trp53-*WT n=6, *Trp53*-null n=6. **E,** Western blot SCD1 protein quantification in isogenic *Trp53*-WT and *Trp53*-null cells cultured in SFM or in ACM. ACM exposure significantly elevated SCD1 protein levels in *Trp53*-null cells. **F**, Quantification of SCD1 protein levels following ACM treatment. The effect of ACM on SCD1 protein expression was significantly stronger in *Trp53*-null cells in comparison to *Trp53*-WT cells. *SCD1* protein levels are shown as fold-change of ACM over SF conditions. One-sided Student’s t-test p=0.02. *Trp53*-WT n=6, *Trp53*-null n=6. **G,** Analysis of FA content by gas chromatography. The content of saturated FAs (SFA) and mono-unsaturated fatty acids (MUFA) was compared between *Trp53*-WT and *Trp53-*null isogenic EMT6 cells exposed to SFM or ACM. Samples were taken after 24h of cell culture. *Trp53*-null cells treated with Astrocyte conditioned Medium show a shift in the saturated fatty acid to monounsaturated fatty acid fraction (SFA/MUFA), consistent with increased fatty acid biosynthesis, and specifically with SFA desaturation by SCD1. The relative fraction of MUFA is significantly higher in *Trp53*-null cells exposed to ACM. ACM treated *Trp53*-null vs -WT cells, One-tailed Chi-square test p=0.001. n=3. Also see Table S13. **H,** Lipid metabolomic analysis of ACM compared to SFM shows abundance of palmitic acid-containing triglycerides in ACM. The SFA palmitic acid is used as a substrate by SCD1 and converted into MUFA. Two-sided Student’s t-test p=0.001. SF n=3, CM n=3, 21 data points. Also see Table S12. **I,** Metabolomic analysis of polar metabolites comparing astrocyte conditioned medium to the base medium used for its generation. Shown is a heatmap of the most differentially abundant metabolites using LC-MS z-scores. Several metabolites that are important substrates utilized for fatty acid synthesis are more abundant in ACM (highlighted in red). Also see fig. S20A and Table S12. **J,** Proliferation of *Trp53*-WT and *Trp53*-null isogenic BC cells grown in medium supplemented with glutamate (75mM). Shown as confluence relative to starting point relative to respective serum-free controls after 2 days of culture. Two-sided Student’s t-test p=0.007. *Trp53*-WT n=6, *Trp53*-null n=6. **K,** Quantitative PCR measuring *Scd1* mRNA levels in *Trp53*-null and *Trp53*-WT cells (GFP+) isolated from brain tumors and FACS sorted for GFP+ tumor cells following intracranial injection into mice. Relative *Scd1* expression was significantly increased in *Trp53*-null cancer cells as compared to isogenic WT-controls. Two-sided Student’s t-test p=0.02. *Trp53*-WT n=10, *Trp53*-null n=8. **L,** Mouse brain tumor sections stained with antibodies against SCD1 in red and against BC cells in green (GFP) following intra-cranial brain injection of isogenic *Trp53*-WT and *Trp53-*null EMT6 cells. Antibodies were used to label SCD1 positive cells (red) and BC cells (green) in mouse brain sections following bi-lateral intracranial injection of EMT6 *Trp53*-WT and *Trp53*-null cells. Representative images showing increased SCD1 antibody staining in *Trp53-*null tumors relative to *Trp53*-WT controls (day 6). Note that the immunostainings shown in this panel were part of a triple staining for SCD1, GFAP and GFP, also shown in Fig.3A. **M,** Quantification of the % of SCD1-positive tumor cells in *Trp53*-WT and *Trp53*-null brain tumors established through intra-cranial injection of BC cells. A higher fraction of the *Trp53*-null cancer cells expresses SCD1 at 6 days post-injection. Data are shown as protein expression fold-change over the average of the *Trp53*-WT tumors. Two-sided Student’s t-test p=0.001. *Trp53*-WT n=9, *Trp53*-null n=9. **N,** Representative image of a human BCBM section labeled with antibodies against TP53 (orange), SCD1 (Red) and the astrocyte marker GFAP (Green). DAPI was used for nuclear staining. Five patient samples were analyzed. **O,** Quantification of the percentage of SCD1-positive cells in human brain metastatic lesions. Cells that lacked p53 expression were significantly more likely to express SCD1. Two-sided Fisher’s-exact test p<0.0001, n=5 tumor samples, 8540 cells were analyzed. See also fig. S21. **P,** Correlation between the intensity of GFAP (reflecting astrocyte infiltration) in human BCBM samples and the number of SCD1-positive cells. Shown as % SCD1-positive cells as a function of GFAP/DAPI. One-tailed Spearman’s correlation rho=0.60, p=0.18. n=5. **Q,** Comparison of the top 500 differential RNAi sensitivities between *TP53*-WT and *TP53*-null human BC cells. Shown are the top 5 enriched pathways. The genes that are more essential in *TP53*-null cell lines are strongly and significantly enriched for the lipid metabolism pathway ‘REACTOME_METABOLISM_OF_LIPIDS’, Ranked #1. p=2*10^-11^, FDR q-value = 5*10^-8^. CCLE BC cell lines, n=26. See also Table S14. Signatures, from top to bottom: REACTOME_METABOLISM_OF_LIPIDS, REACTOME_CELL_CYCLE, REACTOME_POST_TRANSLATIONAL_PROTEIN_MODIFICATION, REACTOME_CYTOKINE_SIGNALING_IN_IMMUNE_SYSTEM, REACTOME_CELL_CYCLE_MITOTIC. **R,** The correlation between the p53 pathway activity score and the sensitivity of human BC cell lines to RNAi-mediated knockdown of *SCD1*. Low p53 pathway activity is associated with high sensitivity to SCD1 inhibition (negative RNAi values represent increased sensitivity). p53 pathway activity was evaluated by calculating a ssGSEA score for the MSigDBgene signature ‘KANNAN_p53_targets_DN’. Spearman’s rho=0.29, p=0.02. CCLE BC cell lines n= 49. **S,** Representative images of GFP-labeled *Trp53*-WT and *Trp53*-null EMT6 cells treated with the SCD1 inhibitor SW203668 (0.1μM) or with control-DMSO for 72hr. *Trp53*-null cells were much more sensitive to the drug treatment. **T,** Live imaging-based proliferation curves of *Trp53*-WT and *Trp53*-null mouse BC cells treated with the SCD1 inhibitor SW203668 (0.1μM) or with control-DMSO for 72h. *Trp53*-null cells were significantly more sensitive to SCD1 inhibition than *Trp53*-WT controls. *Trp53*-null vs *Trp53*-WT p=0.01 at 8h, p=3*10^-4^ at 24h, p=2*10^-4^ at 48h, p=0.001 at 72h; n=3. **U,** Representative images of GFP-labeled BCBM *ex vivo* spheroids derived from *Trp53*-WT or *Trp53*-null isogenic EMT6 cells following 6 days of treatment with the SCD1 inhibitor SW203668 (3μM) or with DMSO-control. **V,** Live imaging-based comparison of spheroid size (µm^2^) between *Trp53*-WT and *Trp53*-null spheroids treated with the SCD1 inhibitor SW203668, relative to DMSO-control conditions, One-sided Student’s t-test p=0.02. See also fig. S22. **W,** A schematic model summarizing the study. p53 inactivation drives BM though adaptations in fatty acid metabolism. FA synthesis and transport are elevated in *TP53*-null primary tumors through p53-dependent expression of FASN, SCD1 and CD36 (cell-autonomous effect of p53 on lipid metabolism). FA synthesis and transport are further stimulated by increased supply of relevant metabolites provided by brain-resident astrocytes, which affect cancer cell growth and migration in a p53-dependent manner, thereby allowing *p53*-null BC cells to thrive in the brain environment.

We next evaluated the expression of *Scd1* in our isogenic mouse EMT6 cells. *Scd1* mRNA and protein levels were higher in *Trp53*-null vs. *Trp53*-WT cells under all culture conditions (**fig. S18G**), and ACM further increased *Scd1* expression preferentially in the *Trp53*-null cells (p=0.004 and p=0.02 for mRNA and protein levels, respectively; **Fig. 5D-F**). Similar elevation was observed with the key FAS enzyme *Fasn* (**fig. S18H**). SCD1 and FASN were similarly elevated in the *TP53*-null vs. *TP53*-WT human CAL51 cells (**fig. S19**). To confirm that the increased expression of SCD1 resulted in its increased activity, we analyzed the FA content of the *Trp53*-null and *Trp53*-WT cells under SFM and ACM conditions using gas chromatography (**Table S13**). The abundance of mono-unsaturated FAs – the product of SCD1 activity – was significantly higher in the *Trp53*-null cells following exposure to ACM (**Fig. 5G** and **Table S13**). Metabolomic profiling of ACM vs. SFM by LC-MS revealed a significant increase in the abundance of palmitic acid (16:0), an SCD1 substrate, in the ACM (**Fig. 5H** and **Table S12**). The metabolomic profiling further revealed that ACM was enriched in polar metabolites (**Fig. 5I**, and **Table S12**) that are required for FA synthesis (**fig. S20A**).

Glutamate and glutamine are important precursors of FAS and are particularly required for FAS under stressful conditions such as hypoxia (*62–64*). Glutamate levels were significantly higher in ACM than in SFM (**fig. S20B**). Interestingly, metabolomic analysis of 24 human BC cell lines (*65*) revealed a significantly lower glutamate content in *TP53*-null vs. *TP53*-WT cell lines (p=0.03; **fig. S20C**), suggesting that p53 inactivated cells might rely more on the glutamate secreted by the activated astrocytes. To ask whether this was indeed the case, we assessed the effect of glutamate supplementation in our isogenic *Trp53*-WT vs. *Trp53*-null EMT6 cells. Remarkably, glutamate supplementation resulted in increased proliferation of the BC cells, which partially mimicked the effect of ACM, and this effect was significantly stronger in *Trp53*-null cells (p=0.007; **Fig. 5J** and **fig. S20D,E**). Importantly, glutamate supplementation led to a significant increase of SCD1 and FASN protein levels, specifically in the *Trp53*-null cells (**fig. S20F-H**), indicating that p53 inactivation indeed increased the ability and/or need of the cancer cells to utilize glutamate for FAS. In line with glutamate transport being required for BM, we found that the expression of glutamate transporters was also upregulated in BCBM in comparison to primary BCs (**fig. S20I**). Specifically, *TP53*-null breast tumors exhibited elevated levels of three glutamate transporters (*66*, *67*) relative to *TP53*-WT breast tumors: *SLC1A*3, *SLC1A5,* and *SLC1A6* (**fig. S20J**). We validated by RT-qPCR that one of these transporters, Slc1a5, was significantly upregulated in the isogenic *Trp53*-null cells cultured under ACM (p=0.04; **fig. S20K**). Together, these results reveal that astrocytes secrete metabolites that are required for FAS, and that these metabolites get utilized by the cancer cells in a p53-dependent manner, resulting in increased FAS in p53-inactivated cancer cells.

We next examined the effect of p53 inactivation on FAS *in vivo*. *Scd1* mRNA and protein levels were significantly higher in BMs from *Trp53*-null vs. *Trp53*-WT BC cells (p=0.02 and p=0.001 for mRNA and protein levels, respectively; **Fig. 5K-M**). Immunofluorescence co-staining of 6 human BCBM samples for GFAP, TP53 and SCD1 confirmed a significant association between p53 status and SCD1 levels (**Fig. 5N,O** and **fig. S21**), and SCD1 levels were positively correlated with astrocyte infiltration (Spearman’s rho = 0.6, p=0.18; **Fig. 5P**). IHC of 17 additional clinical samples of human BCBM, which exhibited astrocyte infiltration and low p53 activity (**Fig. 3S**, **fig. S15** and **Table S5**), also confirmed high levels of SCD1 in these samples (**fig. S15**). These results confirm the clinical relevance of the association between p53 inactivation, astrocytes and SCD1 expression.

### p53 inactivation-induced FAS renders cells sensitive to SCD1 and FASN inhibition

The elevated FAS in p53-inactivated cancer cells raises the hypothesis that these cells may be more sensitive to disruption of this process. Several SCD1 and FASN inhibitors are currently ongoing clinical trials (*59*, *68–71*), emphasizing the clinical importance of identifying biomarkers for drug sensitivity. We therefore analyzed the sensitivity to SCD1 inhibition across 26 human BC cell lines. GSEA of the genes that are more essential in *TP53*-null vs. *TP53*-WT human BC cell lines exposed a strong enrichment for genes related to lipid metabolism (**Fig. 5Q** and **Table S14**). Moreover, low p53 pathway activity was significantly associated with increased sensitivity to SCD1 knockdown (Spearman’s rho = 0.29, p=0.02; **Fig. 5R**) or knockout (p=0.048; **fig. S22A**). To assess causality, we treated our isogenic *Trp53*-WT and *Trp53*-null EMT6 cells with the SCD1 inhibitor SW203668, and found that the p53-inactivated cells were significantly more sensitive to the drug (**Fig. 5S,T** and **fig. S22B,C**). The same differential drug response was identified in cells treated with another SCD1 inhibitor, A939572 (**fig. S22D**). Similarly, *TP53*-null CAL51 cells were more sensitive than their *TP53*-WT counterparts to the two SCD1 inhibitors (**fig. S22E-H**). We next inhibited SCD1 in 3D spheroids derived from the isogenic EMT6-induced BMs, and found that p53 inactivation increased the cellular sensitivity to SCD1 inhibition in this tumor-derived *ex vivo* model as well (**Fig. 5U,V** and **fig. S22I**). These results demonstrate a causal relationship between p53 inactivation and increased sensitivity to clinically-relevant SCD1 inhibitors.

Lastly, we repeated these experiments and analyses with FASN inhibition. *TP53*-null human BC cell lines were significantly more sensitive than *TP53*-WT cell lines to genetic (RNAi-mediated) knockdown and to pharmacological inhibition (with the drug C75) of FASN (**fig. S23 A,B**). Consistently, the isogenic *Trp53*-null cells were significantly more sensitive to the FASN inhibitor, C75, than *Trp53*-WT cells (**fig. S23C-E**), and so were p53-inactivated BCBM-derived 3D spheroids (**fig. S23F**). Together, these results demonstrate that p53 inactivation sensitizes BC cells to inhibition of both of these key FAS enzymes, SCD1 and FASN.

## Discussion

Understanding the molecular underpinning of metastasis organotropism is a key goal of cancer biology, as it may improve the therapeutic management of metastatic patients. Here, we identified p53 inactivation as a major driver of BCBM, and potentially of carcinoma BM in general. We found a cell-autonomous association between p53 inactivation and FA metabolism in BC cells and demonstrated that this association is causal and that it renders p53-inactivated brain-metastasizing cells more sensitive to FAS inhibitors. We also revealed a p53-dependent effect of astrocytes on BC cell survival, proliferation, and migration. Furthermore, we provided evidence for a p53-dependent, non-cell-autonomous effect on FA metabolism as an essential underlying pro-metastatic and brain-adapting mechanism (**Fig. 5W**).

*TP53* is the most studied gene in human cancer (*4*). Why has the overwhelmingly strong association between p53 inactivation and BCBM been previously overlooked? One reason might be that in recent years most of the focus has been on matched analyses of primary tumors and their derived metastases (*61*, *72*). Such analyses make for a clean system to study genomic alterations that arise during metastasis, but they would inevitably miss common drivers that pre-exist in the primary tumors. In the case of p53, we reveal that its status in the primary BC lesion is already strongly associated with the risk for BCBM, so this association would not come up in matched analyses. A second reason is that p53 inactivation is often achieved through loss of the entire chromosome arm (del17p). Aneuploidy is commonly ignored in genomic analyses of metastases, and when del17p is overlooked, the link between p53 inactivation and BCBM becomes much weaker (*18*). In contrast, when aneuploidy status is integrated into the analysis, the association between p53 perturbation and BCBM becomes strong and significant (**Fig. 1C,D**). Therefore, our study emphasizes the importance of integrating aneuploidy calls into cancer genomic analyses.

Our finding that BCBM cells upregulate both FA synthesis and uptake (**Fig. 5W**) is intriguing because these seem to be redundant mechanisms for obtaining the same goal. Why are both mechanisms at play here? Previous studies have begun to reveal the importance of metabolic adaptations – and of lipid metabolic adaptations in particular – in metastasis (*47, 73–75*). The balance between FA synthesis and FA uptake is not sufficiently understood, and could be of much clinical importance. We hypothesize that the low availability of lipids in the BME forces metastasizing cells to rely on both endogenous and exogenous FAs for their survival, migration, and proliferation. Several important questions remain, however: (1) Would CD36 inhibition affect cells in a p53-dependent manner, similarly to the FAS inhibitors? (2) Would BCBM cells respond better to inhibitors of FA synthesis or to inhibitors of FA uptake *in vivo*? (3) Is there functional redundancy between FA synthesis and uptake, enabling the development of drug resistance upon drug treatment? (*14*, *57*, *76–78*) (4) Do other lipids, such as cholesterol – whose biosynthesis is known to be regulated by p53 (*79–81*) – also promote BCBM in a p53-dependent manner?

The finding that FA metabolism is already altered in a p53-dependent manner in primary BCs is biologically important. It is in full agreement with a recent study that demonstrated that the metabolic programs of primary tumors were very similar to those of their metastases, arguing that cancer cells retain many aspects of a metabolic program that is defined by the tissue-of-origin even when exposed to a new metastatic niche (*82*). That study concluded that despite much evidence for specific metabolic adaptations within metastatic niches, the metabolic program of the primary tumors may constrain cancer metastasis (*82*). This is in line with older gene expression analyses showing that the metastatic potential of human tumors is encoded in the bulk of the primary tumor (*83*). We believe that our results provide a novel, concrete example of that idea, demonstrating the important interplay between cell-autonomous and non-cell-autonomous lipid metabolism in BCBM.

The establishment of p53 inactivation in the primary tumor as a significant risk factor for metastasis to the brain will also be of high clinical value, given the poor prognosis associated with BCBM, and the lack of molecular predictors of this devastating condition. What are the concrete clinical implications of our findings? First, we show that p53-inactivated cells are more sensitive to FAS inhibitors, and in particular to SCD1 inhibitors, suggesting p53 status as a potential biomarker of patient stratification for drug treatment. Our results also suggest FAS as a potential clinical liability of BMs. Second, our results suggest that BC patients with a *TP53* genetic alteration are considerably more likely to develop BCBM. Patients with elevated risk for BCBM can thus be identified based on the genomic evaluation of their primary tumors, which could affect patient care in at least two ways: (a) This particular subtype of patients may specifically benefit from frequent head MRIs – a medical procedure not routinely offered to BC patients in most healthcare systems. (b) For patients with tumors of the HER2-enriched molecular subtype, several existing targeted therapies differ both in their ability to penetrate the blood-brain barrier (BBB) and in their overall toxicity;(*84*) for example, the small molecule tyrosine kinase inhibitor, Neratinib, (as opposed to anti-HER2 antibodies) is a drug that penetrates relatively well to the brain but is associated with a relatively high toxicity (*85*). Knowing whether a patient is likely or not to develop BCBM could therefore affect the clinical cost-benefit analysis of such drugs.

Our study raises several intriguing questions for future exploration: (1) Is the identified axis of p53 inactivation – astrocyte infiltration – FA metabolism relevant also to primary brain tumors? Glioblastomas commonly inactivate p53 (*86*), and have been recently shown to depend on FAS(*49*, *87–90*), suggesting that our findings may be relevant to primary brain tumors as well. (2) What is the exact molecular mechanism by which p53 regulates FA metabolism in the context of BCBM? P53 inactivation was previously shown to affect lipid metabolism (*91–95*), and also to specifically upregulate FAS (*96*), however, the molecular interactions between the p53 pathway and SCD1 remain to be elucidated. (3) Here, we focused on the effect of p53 inactivation on FA metabolism, however, it is plausible that p53 inactivation contributes to BCBM through additional mechanisms. Likely candidates include altered chemokine signaling (*11*, *97–99*), as well as the regulation of matrix metalloproteinases (MMPs)(*100*, *101*) and Serpins (*102–104*), which have been previously implicated in BM and also been shown to be regulated by p53 (*101*). Another potential mechanism is mTOR signaling (*105–109*), which is upstream of FAS (*110*), is responsive to amino acids such as glutamate (*111*), and has been implicated in BM. It is thus tempting to speculate that astrocytes may affect the activity of the mTOR pathway in BCBM in a p53-dependent manner. Notably, several of these mechanisms are clinically targetable. (4) We report the loss of chromosome 17p, del17p, to be extremely common in BCBM. While it is clear from our results that p53 inactivation is a consequential outcome of del17p, this large-scale chromosome-arm loss may affect BCBM through additional, p53-independent mechanisms; whether such mechanisms exist – and might be targeted – remains unknown. (5) Lastly, we have found that aneuploidy-driven perturbation of a tumor suppressor gene could have a profound impact on metastatic organotropism. It will be important to further explore the role of aneuploidy in determining metastatic organotropism in BC and across all cancer types.

## Acknowledgements

We thank Ayelet Erez, Neta Erez, Amir Sonneblick, Giannino Del Sal, Shai Izraeli and Ofer Shoshani for helpful discussions; Maxim Itkin and Sergey Malitsky from the metabolomic facility at WIS for their assistance with metabolomics; Paula Ofek, Eilam Yeini and Yael Zilberstein for assistance with intracranial and intracardiac injections, monitoring and imaging of mice; Irena Shur and Daria Makarovsky for assistance with FACS sorting; Shai Israeli, Angela Savino, Angela Ruban, and Yona Goldshmit for the sharing of reagents; Wei Shi and Yang Liao for pre-processing of the RNAseq from the BROCADE project; and Paola Cassoni for funding support of the Bertero lab.

We acknowledge the patients who donated samples, including patients and their family who consented to donate to the BROCADE Rapid Autopsy Program. We are grateful for the support of BROCADE, initially funded by the Australian National Breast Cancer Foundation (NBCF) under infrastructure grant IF-14-001. In particular, we would like to thank Robin Anderson and Lisa Devereux, who manage BROCADE and helped with the identification, collection and processing of samples. We also acknowledge the Victorian Institute of Forensic Medicine that conducted the rapid autopsies; the Peter MacCallum, Kinghorn Centre, Garvan Institute, and Tobin Brothers Funeral Directors for providing transport of donors.

## Author contributions

U.B.-D. and K.L. conceived the study. U.B.-D. directed and supervised the overall study. R.S.-F. directed and supervised the *ex vivo* spheroid experiments and *in vivo* work. K.L. performed most of the computational analyses and *in vitro* experiments, assisted with *in vivo* experiments, and led the data analysis of all experiments. S.P. performed most of the *in vivo* animal experiments, spheroid experiments, and astrocyte derivation. Y.C.-S., T.W. and J.Z. performed *in vitro* experiments. T.W., Y.E. and K.L. performed genomic data analyses. S.D.-I. assisted with the animal experiments. A.I.L.-F. performed the gas chromatography experiments. A.A.R., L.B. and contributed clinical samples of BCBM and performed IHC experiments. I.B. contributed clinical samples of BCBM. A.S. and G.B. performed RNAseq of clinical samples of BCBM. J.B. and D.M. performed RNAseq of clinical samples of metastases from various organs. A.S. and H.M. provided experimental support. U.B.-D. and K.L. wrote the manuscript with input from all co-authors.

## Data and materials availability

RNA sequencing data of the metastasis cohort GSE245414 was deposited at the Gene Expression Omnibus (GEO). RNA sequencing data of patient autopsy samples is available upon request.

