## Supplementary Information for "Inactivation of p53 drives breast cancer brain metastasis by altering fatty acid metabolism"

#### Supplementary Materials

##### Material and Methods:

**Clinical genomic data sets.** TCGA PanCancer Atlas (*35*) https://www.cbioportal.org/study/summary?id=pancan_pcawg_2020, METABRIC (*6*, *112*, *113*) https://www.cbioportal.org/study/summary?id=brca_metabric, The Metastatic Breast cancer project (MBP) (*31*, *32*) https://www.cbioportal.org/study/summary?id=brca_mbcproject_2022, and MSK data sets (MSK-IMPACT 2017 (*37*) https://www.cbioportal.org/study/summary?id=msk_impact_2017, MSK-Cancer Cell 2018 (*17*), MSK-MET 2021 (*18*) https://www.cbioportal.org/study/summary?id=msk_met_2021) were obtained from cBioPortal (*114*). Additional cohorts of primary breast tumors with known metastatic sites (*33*) were downloaded from the Gene Expression Omnibus (GEO) repository (accession numbers GSE12276 (*33*) https://www.ncbi.nlm.nih.gov/geo/query/acc.cgi?acc=gse12276 and GSE76714 (*34*) https://www.ncbi.nlm.nih.gov/geo/query/acc.cgi?acc=GSE76714). Datasets containing expression data of metastasis samples were also downloaded from GEO (accession numbers GSE14017 https://www.ncbi.nlm.nih.gov/geo/query/acc.cgi?acc=GSE14017 and GSE14018 https://www.ncbi.nlm.nih.gov/geo/query/acc.cgi?acc=GSE14018)
(*27*), allowing for the comparison between metastatic sites. A third new cohort containing 6 brain and 26 other metastases was profiled using RNAseq as described below, and is available at GEO (GSE245414). Paired primary breast tumor and BM expression profiles were obtained from GEO (accession number GSE125989 https://www.ncbi.nlm.nih.gov/geo/query/acc.cgi?acc=GSE125989 (*61*)). Expression data from several metastatic sites of a single patient at autopsy was obtained as described below.

**Cancer cell line genomic data sets.** CAL51 *TP53*-KO RNAseq was taken from Redman-Rivera et al. (*115*). RNA expression data and metabolomics data (*65*) for 1019 human cancer cell lines was downloaded from the Cancer Cell Line Encyclopedia (CCLE, https://portals.broadinstitute.org/ccle/data). The metastatic potential of 489 human cancer cell lines to several metastatic sites (brain, lung, liver, bone and kidney) in mouse xenografts was taken from the MetMap500 paper (*14*). CRISPR (CERES), RNAi (Achille, DRIVE, Marcotte,

DEMETER2) dependency scores and drug screen data (GDSC1) were obtained from DepMap 22Q1 release (https://figshare.com/articles/dataset/DepMap_22Q1_Public/19139906).

**Assignment of *TP53* status.** *TP53* mutations and copy number alterations (CNA), including chromosome arm-level deletions (del17p) were assigned to clinical samples as follows. *TP53* mutational status and *TP53* copy number or copy number segments were downloaded for the following clinical datasets (TCGA (*35*), METABRIC (*6*, *113*), MSK-IMPACT (*37*), MSK-Cancer Cell 2018 (*37*), MSK-MET 2021 (*18*) and MBC (*32*) datasets) from cBioPortal (*114*). *TP53* mutations were extracted using the oncoprint function (HOMDEL, HETLOSS, MUT, AMP, GAIN). When *TP53* CN was not available, copy number segments were used to determine TP53 focal deletions defined at a threshold of ≤−0.3 for the loss of the *TP53* gene. For chromosome 17p arm-level calls, copy number segments were visualized using the IGV genome browser (https://software.broadinstitute.org/software/igv/). Loss of chromosome 17p (del17p) was defined at a threshold of ≤−0.3 (loss) across ≥70% of the chromosome arm. Alternatively, genes located on chromosome17p were used to determine chr17p status from WGS data. *TP53* status was then manually annotated and assigned to the following categories: Samples that contained no *TP53* mutations nor *TP53* copy number alterations were assigned '*TP53*-WT'. Tumors that carried any *TP53* mutation were classified as '*TP53*-mut', and further divided into '*TP53*-hotspot' mutations (R175H, R275H and R243) or '*TP53*-other' mutations. Samples that had no *TP53* mutations were assigned '*TP53*-no-mut'. Tumors that contained a heterozygous *TP53* deletion were labeled '*TP53*-del'. Tumors with homozygous *TP53* perturbations were labeled as '*TP53* biallelic inactivation' or '*TP53*-null', both when they contained any *TP53* mutation and an additional *TP53* copy number loss (*TP53*-mut/TP53-del), or in rare cases *TP53* deep deletions (*TP53*-del/del).

The *TP53* status of cancer cell lines was assigned in the same way. *TP53* mutational data and CN data were downloaded from https://portals.broadinstitute.org/ccle/data, DepMap 22Q1 release (https://figshare.com/articles/dataset/DepMap_22Q1_Public/19139906).

**Gene set enrichment analysis (GSEA).** Gene expression profiles were obtained from the above-mentioned data sets. Normalized matrix files were downloaded, and samples were manually annotated according to the provided sample information. Raw expression matrices were processed following a three-step protocol: (1) The lowest 10-20% expressed genes were defined and set as the expression threshold value. (2) Using this threshold value, genes not expressed in >20% of the samples were also removed. (3) Lastly, genes with lower expression values were raised to the threshold value. The GSEA tool (https://www.gsea-msigdb.org/gsea/index.jsp) (*116*) was used with the following parameters:: Number of permutations – 1000, collapse/remap to gene symbols – collapse, permutation type – geneset, Chip platform: ftp.broadinstitute.org://pub/gsea/annotations_versioned/Human_Gene_Symbol_with_Remapping_MSigDB.v2023.1.Hs.chip. Single cell gene set enrichment analysis (ssGSEA) was performed using GenePattern (https://www.genepattern.org/) (*117*).

**Correlating BM prevalence with *TP53* perturbation prevalence.** Across carcinomas: BM prevalence in different cancer types were taken from Riihimäki et al. (*36*). The analysis focused on carcinomas, and other tumor types were excluded. For lung and kidney, the highest-ranking subtypes were considered. *TP53* perturbation prevalence in the respective cancer types were obtained from cBioPortal as described above (TCGA data (*35*)). Across subtypes: For the analysis of the brain metastatic potential of BC subtypes, BM prevalence was taken from Smid et al, (*118*) and *TP53* perturbation prevalence in the respective molecular subtypes were calculated from METABRIC data (*6*, *112*, *113*).

**Cell culture conditions.** BC cell lines: EMT6 cells were obtained from ATCC and CAL51 cells were obtained from the Broad Institute cell line repository (*119*). Cells were cultured in DMEM (Life Technologies) with 10% fetal bovine serum (Sigma-Aldrich), 4mM glutamine, 100IU/ml penicillin and 100µg/ml streptomycin. Cells were maintained in culture for a maximum of two weeks. Human and Murine Astrocytes: Human astrocytes (ScienceCell Research Laboratory, Carlsbad, CA, USA, # 1800) and freshly isolated murine astrocytes were cultured in Astrocyte medium (ScienceCell, #1801) supplemented with 2% FBS, 1% astrocyte growth supplements, 100U/mL Penicillin and 100µg/mL Streptomycin. Astrocyte cultures were not propagated beyond passage 5**.** All cells were grown at 37°C in 5% $CO_2$ and routinely tested free for mycoplasma contamination with a mycoplasma detection kit (Biological Industries, Israel, or Lonza, Walkersville, MD, USA) according to the manufacturer's protocol. Culture of BC cells in ACM: EMT6 cells were seeded in a 6-well cell culture plate at a density of 80,000 cells per well or in a 6cm plate at a density of 250,000 cells per plate. The following day, cells were starved for 4h in serum-free DMEM or astrocyte medium. Following starvation, isogenic cells were grown in freshly prepared ACM for 16h to obtain RNA samples or slightly longer, 24-30h, for protein lysates. The number of cells was counted for an initial assessment of the effect of the ACM, and all samples were adjusted for cell number before being processed for RNA or protein extraction. Co-culture of BC cells and astrocytes: Astrocytes were seeded in a 96-well plate at a density of 1,000-3,000 cells per well. Cells attached overnight, and cancer cells were seeded the next morning at a ratio of 1:1 astrocytes to cancer cells. Cancer cells were allowed to attach and medium was then exchanged to serum-free astrocyte medium. Proliferation was assessed by live-cell imaging as described below.

**Generation of astrocytes conditioned medium (ACM).** For the preparation of human ACM, 0.5-1*$10^6$ human astrocytes were seeded in a 10cm cell culture dish in full astrocyte medium. After 24h, medium was exchanged to astrocyte medium without glutamine (ScienceCell, #1801-NG) astrocyte base medium containing only 100 IU/mL Penicillin and 100 µg/mL Streptomycin (AM-SF) and astrocytes were cultured under these starvation conditions for 24h. For the preparation of mouse ACM, 0.75-1*$10^6$ of isolated murine astrocytes were similarly seeded but incubated in AM-SF supplemented with 100 ng/mL LPS (*120*) for 24h. Following incubation, ACM was collected and filtered through a 0.22-µM filter and was used freshly on target cells.

**Fluorescent labeling of BC cell lines.** For red fluorescent labeling**,** EMT6 and CAL51 BC cells were labeled with pQC-mCherry retroviral particles, as previously described (*121*). For green fluorescent labeling, eGFP (pLEX_TRC206) was packaged using a 3rd generation lentiviral system. Briefly, lentiviral vector (1µg) and its packaging vectors (pMDLg/pRRE 0.65ug, pRSV.Rev 0.25ug, pMD.G (VSVG) 0.35µg) were transfected into 293T cells using JetPEI transfection reagent (Polyplus, #101-10N). The morning following transfection, the medium was replaced with fresh culture medium. 48 and 72h later, the lentivirus containing medium was collected, filtered through a 0.45-µm filter and the target cells were infected with the fresh lentivirus containing media (supplemented with 8 mg/ml polybrene (Sigma, # TR-1003-G)). The next day, the medium was replaced with fresh culture medium. Cells were expanded for two passages and then FACS-sorted to obtain a pure population of eGFP-positive cells.

**Generation of isogenic P53-WT/P53-null cells by CRISPR-Cas9 gene editing.** *Trp53* and *TP53* target sequences were cloned into lentiCRISPRv2(*122*) or lentiGuide vectors following standard protocols (*123*). sgRNAs targeting LacZ or EGFP were generated as non-targeting controls (*124*, *125*). Blast search confirmed no known target sequences in the mouse and human reference genomes. Constructs were validated by Sanger Sequencing. Targeting sequences can be found in Table S15. *TP53/Trp53* sgRNA- or control sgRNA-containing vectors were packaged in HEK293T cells using a 3rd generation lentiviral system. Lentiviral particles were obtained as described above. Target cells were infected with the fresh lentivirus containing media (supplemented with 8mg/ml polybrene). The next day, the medium was replaced with fresh culture medium containing selection antibiotics and stable clones were selected with antibiotics. Following antibiotic selection, EMT6 single cells were seeded into 96-well plates, using FACS, grown as single cell colonies and subsequent sanger sequencing confirmed the introduction of homozygous *Trp53* deletions. Five such *Trp53*-WT or *Trp53*-null clones were pooled, expanded and frozen as aliquots. These aliquots were used for all experiments and cells were further propagated in cell culture for up to two weeks. P53 deficiency was further validated using p53 western blot and qPCR for *Trp53* and its target genes. Similarly, CAL51 *TP53*-WT cells were transduced with either control sgRNA or two sgRNAs targeting two exons of the *TP53* gene. After antibiotic selection, knock-out was validated by western blot and qRT-PCR. Given the high efficiency of TP53 knockout at the population level, CAL51 cells were not single cell-cloned. Cells were propagated in culture for up to two weeks.

**Quantitative real time PCR (qRT-PCR).** Cells were collected using Bio-TRI® (Bio-Lab, # 009010233100) and RNA was extracted following manufacturer's protocol. cDNA was amplified using GoScript™ Reverse Transcription System (Promega, #A5001) following manufacturer's protocol. qRT-PCR was performed using Sybr® green (Applied Biosystems; AB-4367659), and quantification was performed using the $\Delta$CT method. *18s*, *Gapdh* and/or *Hprt* served as housekeeping genes for normalization purposes. Experiments involving mouse ACM were normalized against additional housekeeping genes, such as *Stx5* and *Ubc*, that were stable when exposed to LPS (*126*). Primer sequences are available in Table S15.

**Western Blot.** Cells were lysed in RIPA lysis buffer (20mM Tris-HCl (pH 7.5), 150mM NaCl, 1mM EDTA, 1mM EGTA, 1% NP40, 1% sodium deoxycholate) with the addition of protease inhibitor cocktail (Sigma-Aldrich #P8340). Protein lysates were resolved on 10% SDS-PAGE gels. Bands were detected using chemiluminescence (Millipore; WBLUR0500) on Alliance Q9 (UVITEC). Western Blot quantifications were performed using ImageJ®. Antibody details are available in Table S15.

**Live-cell imaging.** The viability, proliferation and motility of living cells were followed using an IncuCyte S3 Live Cell Analysis System (Sartorius) or the LiveCyte cell analysis system (Phasefocus). For assessment of cell proliferation, BC cells were plated in a 96-well plate (EMT6: 750-1,000 cells/well CAL51: 3,000 cells/well) in 3-5 technical replicates. Photos were taken every 2-6h until the culture became confluent, using an Incucyte (Satorius). To calculate the confluence, the Built-In program (2021A version) was used, applying a threshold of 1 and a minimum area of 140$\mu m^2$ to exclude the debris. For assessment of cell migration characteristics such as displacement, confinement, tracks and speed the LiveCyte was used. Briefly, BC cells were seeded at a density of 750-2,000 cells/well in microscopy-compatible 96-well plates (Corning). Starting the next day cells were recorded using video microscopy with an inverted microscope using a 10X objective (microscope placed in an incubation chamber maintained at 37°C with 5% CO2). Images were acquired using the LiveCyte acquisition software, and single-cell tracking, segmentation and analyses were performed using the LiveCyte analysis software (Phase Focus). Displacement, confinement and speed were calculated by the automatic LiveCyte analysis software (Phase Focus).

**Cell death analysis.** For cell death analysis by flow cytometry, EMT6 cells were cultured with either ACM or with SFM for a period of 48h, collected and stained with antibodies for Annexin-V and Propidium Iodide (PI). Briefly, cells were washed twice with cell staining buffer (Biolegend; #420201) before being resuspended in cell binding buffer (Biolegend; 422201). Cells were incubated with 2.5µl Annexin-V (Biolegend; #640920; 12µg/ml) and 1µl PI (Biolegend; #421301; 0.5mg/ml) for 15 mins in the dark at room temperature.

CAL51 cells were similarly treated with ACM or SFM for 72h, stained with Sytox green (S34860, Invitrogen, 1:1000) and Pacific Blue-AnnexinV Biolegend; #640918; 60µg/ml) for 15 min at room temperature. Data acquisition was performed using the Cytoflex flow cytometer (Beckman Coulter). Data analysis was performed using CytExpert analysis software 2.4 (Beckman Coulter). For cell death analysis by live-cell imaging, Sytox$^{TM}$ dead cell stain (Invitrogen; S34860) was used at a 1:10000 dilution and followed by live-imaging as described below.

**Cell migration assays.** Cell migration was assessed using a transwell migration assay (Costar Transwell Permeable Support, #3422). 50,000 astrocytes were seeded in full astrocyte growth medium in the bottom part of the transwell cell culture plate. After 24h, the medium was exchanged for 350µl serum-free astrocyte medium. 20,000-50,000 cancer cells were seeded in SFM into the insert and after 2h the insert was transferred onto the well containing the astrocytes. Cell migration was stopped by fixation in methanol or 4% PFA. BC cells were subsequently stained with Hemacolor (Merck) and/or detected by fluorescence. For EMT6 cells, migration was stopped after 20h, whereas for CAL51 cells (which migrated considerably more slowly) the assay was stopped after 48-72h. In a modification of this assay, the astrocytes were replaced by ACM as follows. 350µl fresh ACM was placed in the lower bottom part of the transwell set up and replaced every 24h. The insert containing the BC cells was placed immediately onto the ACM. The assay was developed as described above.

**Ethics and study approval.** Animal studies: All animal procedures were performed in compliance with Tel Aviv University and approved by the Institutional Animal Care and Use Committee (TAU IACUC protocol no. 01-20-074). Human tissue samples presented in Fig. 5N-P and fig. S21: Formalin-Fixed Paraffin-Embedded (FFPE) from clinical samples of BCBM (n=6) were obtained from Sheba Medical Center following an informed consent. The manipulation of the human samples for immunostaining was accepted by the ethics committees of Tel Aviv University and Sheba Medical Center, under an approved institutional review board (IRB no. 5727-18-SMC). Human tissue samples presented in Fig. 3S and fig. S15: A series of 17 BCBM with available histological material and follow-up data was retrieved from the pathology files of the University of Turin/Città della Salute e della Scienza Hospital of Torino between 2014 and 2017. The study was conducted in accordance with the Code of Ethics of the World Medical Association

(Declaration of Helsinki and following amendments) for experiments involving humans and within the guidelines and regulations defined by the University of Turin. Informed consent to surgical procedure and data collection was obtained from all subjects involved in the study. RNA sequencing of human metastases samples (GSE245414): The study was approved by the ethics committee and institutional review board of Regina Elena National Cancer Institute and appropriate regulatory authorities (approval no. IFO 1270/19); all patients provided written informed consent. The study was conducted in accordance with the Declaration of Helsinki and guidelines for Good Clinical Practice, as defined by the International Conference on Harmonization. The BROCADE rapid autopsy program (www.petermac.org/research/research-cohort-studies/brocade) operates under oversight of the PMCC HREC, approval 15-24. Donors provided written informed consent during life for donation of tissue soon after their death in a rapid autopsy procedure. Recruitment discussions are initiated by the treating clinician and involve the nominated next of kin.

**Astrocyte isolation from mice.** Brains from 8-10 week-old C57BL/6 mice were isolated, cut into small pieces and digested with Collagenase III/ Dispase solution (Worthington Biochemical Corporation, LS02104/ LS004186) for 50 min at 37°C under rotation. Myelin was removed using a percoll gradient and red blood cells were lysed. Microglia and endothelial cells were then removed from the cell suspension using CD11b and CD31 microbeads. Following negative selection, the remaining cell population was highly enriched for astrocytes (98% purity by FACS using anti GFAP antibody) and was plated in a 10 $cm^2$ dish and grown in astrocyte medium, as previously described (*99*).

**Intra-cardiac injection experiments.** To assess the metastatic potential of *Trp53*-WT / *Trp53*-null isogenic cells, $0.2*10^6$ GFP-labeled EMT6 BC cells were injected into the left ventricle of immunocompetent BALB/c mice under guidance of an ultrasound device (2 cohorts, 15 females/group). Ten days after injection, mice were euthanized, and their organs (brain, liver, spleen, femoral bone, and lungs) were harvested and imaged for GFP fluorescence using the Cambridge Research & Instrumentation, Inc. (Cri)-MAESTRO device (The University of Texas Southwestern Medical Center, Dallas, Texas, USA). Multispectral image-cubes were obtained through 550–800 nm spectral range in 10 nm steps using excitation (575–605 nm) and emission (645 nm longpass) filter sets. Mice autofluorescence and background signals were eliminated by spectral analysis and linear un-mixing algorithm. Tumor characteristics were quantified using image J. In a modified version of this experimental setup (competition assay), one group of mice (10 females) was similarly injected with both *Trp53*-WT (labeled with mCherry) and *Trp53*-null cells (labeled with eGFP), $0.5*10^6$ cells per genotype. After 8 days the animals were imaged by MRI, then sacrificed and their organs were analyzed for fluorescence using a Maestro device as described above.

**Intra-cranial injection experiments**. For brain growth adaptation experiments, EMT6 or CAL51 isogenic *Trp53*/*TP53*-WT/null cells ($1.5*10^4$ and $5*10^4$ cells/2µl, respectively) were inoculated stereo-tactically to the striatum (2 mm left and right from the Bregma (*Trp53/TP53*-null) and 3.5 mm depth (*Trp53*/*TP53*-WT)) of 8 to 10-week-old BALB/c or SCID female mice, respectively. Injections were either performed bilaterally into the two brain hemispheres, to compare tumor growth of *Trp53*/*TP53*-WT/null isogenic cells within the same brains (Cohort size: 10-15 mice/experiment), or unilaterally, allowing for the assessment of survival and the isolation of tumor cells from the brain. Mice body weight was monitored twice a week, and tumor growth was followed using a 4.7T/1H MRI. For EMT6 experiments, MRIs were taken every 3 days, usually starting on day 4. At endpoint, mice were euthanized and immediately perfused with 4% PFA in PBS and brains were harvested for immunohistochemistry (day 6 and 8). For CAL51, tumors were growing considerably slower and tumor growth was monitored by MRI on day 7, 14, 22, 29 and 36.

**Magnetic Resonance Imaging (MRI).** All mice were scanned with a conventional T1 or T2 protocol in the same scanning session on 4.7T MRI or 7 T MRI (MR Solutions, UK) using 32-channel head-coil, followed gadolinium (Soreq, Israel) intraperitoneal injection. T1-tomography data were analyzed using RadiAnt DICOM Viewer or MRIcro software, the tumor area was calculated per each axial scan and the tumor's volume was obtained by summing the tumor's areas of all slices.

**Isolation of BC cells from mouse brain tumors.** GFP-labeled EMT6 cells (*Trp53*-WT or *Trp53*-null) were intracranially inoculated as described above. Brains were harvested, tumors were resected, cut into small pieces, and incubated in rotation with Collagenase III (Worthington Biochemical Corporation, LS004186)/ Dispase (Worthington Biochemical Corporation,

LS02104) solution for 50 min at 37°C. Red blood-cells lysis was carried out followed by Percoll gradient for myelin separation.

**Extraction of RNA and cDNA synthesis from isolated mouse BM.** Following BC cell isolation from brain tumors on day 5 after intra-cranial injection into mice, *Trp53*-WT or *Trp53*-null cells were FACS-sorted based on their GFP fluorescence (AriaIII BD Bioscience, USA). 20,000 cells were collected for each sample and snap-frozen in liquid nitrogen. To extract RNA from these samples, a modified protocol of the phenol-chloroform method was used. Cell pellets were incubated with 100µl Bio-TRI® (Bio-Lab) at -80°C for 1h, after which chloroform was added at a dilution of 1:5 and tubes were vortexed well. Following a 3 min incubation at RT, samples were centrifuged at 12,000g at 4°C for 15 min to allow phase separation. The top clear phase was transferred to a fresh tube pre-prepared with 100µl isopropanol and 1µl glycogen (Thermo Fisher; R0561). Tubes were then incubated overnight at -20 °C. The following day, RNA was pelleted, washed and resuspended in DEPDC-treated water. cDNA was prepared using the qScript® cDNA Synthesis Kit (Quantabio; 95047-025) following the manufacturer's instructions.

**RNA sequencing from patient tumor samples.** Tumor material from patient's surgery or biopsy were collected in RNAlater (Thermo Fisher) and cryopreserved in liquid nitrogen for subsequent extraction of nucleic acids (RNA and DNA). RNA was extracted from fresh frozen tissue using the AllPrep DNA/RNA/miRNA Universal Kit (Qiagen, Valencia, CA, USA). The quality of the RNA was assessed with by Bioanalyzer using the Agilent RNA 6000 Nano Kit. Libraries for RNA sequencing were prepared using the TruSeq RNA Exome kit (Illumina) following the manufacturer's instructions. The quality of the resulting libraries was confirmed by Bioanalyzer (High Sensitivity DNA Kit), the intermediate library prior to exon enrichment was quantified by Bioanalyzer using the Agilent DNA 1000 kit, and the final library was assessed by qPCR. Samples were sequenced in paired-end mode, sequencing 76bp from each side on a NextSeq 500 instrument. RNA-seq data were analyzed with "rnaseq" version 3.9 pipeline of nf-core community RNA sequencing analysis pipeline (https://nf-co.re/rnaseq). The alignment was performed with the STAR algorithm and the gene quantification with the Salmon algorithm. The pipeline was run using the default parameters.

**RNA sequencing analysis of patient autopsy samples.** Patient 6071 was diagnosed with invasive ductal carcinoma with micropapillary features. Despite treatments, the patient deceased of advanced disease and metastases from the brain, lung (right and left), spine, lymph node, bone and the left rib were collected shortly after death. The mRNA was extracted from the patient autopsy samples using the miRNEasy kit (Qiagen, #217084) according to the manufacturer's recommendations. Briefly, after being crushed in liquid nitrogen with a mortar and pestle, the samples were resuspended in the lysis buffer and the mRNA was isolated using a guanidinium thiocyanate-phenol-chloroform extraction approach. Remaining genomic DNA was digested by RNase-Free DNase (Qiagen, #79254) on the purification column. The RNA was washed and eluted in water. The quantity and the quality of the isolated mRNA were assessed using the TapeStation (Agilent, 4200 TapeStation System), and 100 ng of the mRNA was used as an input for the library preparation using the TruSeq RNA Library Prep Kit (Illumina) according to the manufacturer's recommendations. The indexed libraries were pooled and diluted to 1.5pM for paired-end sequencing (2 × 81 cycles) on a NextSeq 500 instrument using the v2 150 cycle high output kit (Illumina) as per the manufacturer's instructions.

RNA-seq reads were aligned to the human reference genome GRCh38/hg38 using the Subread aligner (*127*). Read counts were generated for NCBI RefSeq human genes (build 38.2) using the featureCounts program (*128*). Genes were excluded from analysis if they failed to achieve a CPM (counts per million reads) value of 1 in at least one library. Genes with no official symbol were also removed from analysis. Counts were converted to log2-CPM and then quantile-normalized using voom function in limma package (*129*, *130*). Log2-CPM values were then further transformed to log2-RPKM (log2 reads per kilo bases per million reads) values.

**Immune staining on mouse brain tumors.** Tumor-bearing mice were anesthetized using ketamine (100 mg/kg) and xylazine (12 mg/kg) and perfused with 4% PFA in PBS. Mouse brains were resected, incubated with 4% PFA for 4h followed by 0.5M of D-Sucrose for 1h, and 1M D-Sucrose overnight. Tissues were then embedded in optimal cutting temperature (O.C.T.) on dry ice and stored at -80°C. Frozen O.C.T.-embedded tissues were cryo-sectioned into 5μm thick sections. Slides were fixed and permeabilized in acetone for 20 min at RT. Briefly, slides were incubated with goat serum (10% goat serum in PBS, 0.02% Tween-20) for 30 min. Slides were stained for morphology by hematoxylin and eosin (H&E), or immunostained with the following antibodies: anti-mouse/human/rat GFAP, anti-human p53, anti-mouse/human Iba1, anti-mouse

CD31, anti-mouse/human Ki67anti-mouse/human/rat SCD1, anti-GFP, and anti-mCherry. After 1h incubation, slides were incubated with secondary antibodies for an additional 1h. Nuclei were counterstained using Hoechst solution. The stained tissues were then fixed and mounted on a glass microscope slide using ProLong™ Gold antifade mounting. Fluorescence and brightfield images were captured using a fluorescence and brightfield illumination microscope (Evos FL Auto, life technologies, Cytation10) at 40X, 10X and 4X magnifications. Antibody details are available in Table S15.

**Immune staining on human BM**. Cohort of 6 BCBM, presented in Fig. 5N-P and fig. S21: Formalin-Fixed Paraffin-Embedded (FFPE) clinical samples of BCBM were sectioned into 5μm thick tissue sections. The sections were heated at 60°C for 30 min, deparaffinized using Dewax solution (Leica Biosystems) and treated with antigen retrieval solution (Leica Biosystem) for 20 min. Then, slides were incubated with 10% goat serum for 30 min and immunostained with the following antibodies: rabbit anti-mouse/human/rat SCD1, mouse anti-human p53, and rat anti-mouse/rat/human GFAP. All slides were processed and stained using the Bond RX automated stainer (Leica Biosystems). Nuclei were counterstained using Hoechst solution. The stained tissue sections were then fixed and mounted on a glass microscope slide using ProLong™ Gold antifade mounting. Fluorescence and brightfield images were taken with a fluorescence confocal microscope (Cytation10) at 40X magnification.

Cohort of 17 BCBM, presented in Fig. 3S and fig. S15: For each BCBM case, a representative FFPE tissue block was cut into 3 μm sections and stained with H&E or submitted to IHC performed by an automated platform (Ventana BenchMark AutoStainer, Ventana Medical Systems, Tucson, AZ, USA). Sections were heated overnight (60°C) and deparaffinized with EZ Prep Concentrate solution (Ventana Medical Systems, AZ, USA). Antigen retrieval was performed using ULTRA Cell Conditioning solution (ULTRA CC1, pH 8.5; Ventana Medical Systems, AZ, USA) at 95°C. FFPE sections were then immunostained with the following primary antibodies: anti-SCD1 (Abcam, ab19862), anti-p53 (clone DO.7), anti-p21$^{WAF1}$ (clone DCS-60.2), anti-Glial Fibrillary Acidic Protein (GFAP) (clone EP672Y). Further details on antibodies are provided in Table S15. UltraView universal DAB detection kit was used to stain for p53, p21 and GFAP; OptiView DAB IHC detection and optiView amplification kits were used for SCD1 stainings. For all reactions, ultraView universal HRP multimer was used as secondary antibody cocktail. Sections were counterstained with Hematoxylin and Bluing reagent (Ventana Medical Systems, AZ, USA).

**IHC quantification on mouse and human brains.** For the analysis of the cohort of 6 BCBM samples, images were analyzed using imageJ. For mouse tumors, the tumor area was determined through GFP fluorescence of BC cells. Staining intensity of GFAP, IBA1, CD31 and SCD1 immune stainings was measured within the GFP-positive tumor area. For each tumor, the mean intensity between all pictures taken (3-7 photos per tumor) was calculated. For the analysis of human BCBM, cells were divided based on their p53 status (p53-expressing cells vs non p53-expressing cells). Cells were then analyzed for SCD1 staining, and the percentage of SCD positive cells within the p53-expressing and non-expressing cell populations was compared. In addition, for each tumor the area of GFAP was normalized to the DAPI area. The association between GFAP/DAPI area to the percentage of SCD1 positive cells was analyzed using Spearman's correlation and linear regression.

For the analyses of SCD1, GFAP and TP53 in an additional cohort of 17 human BCBM, IHC stainings were evaluated as follows. For SCD1 IHC, cytoplasm staining was assessed using the H-score (range: 0-300), which represents the sum of the products between the staining intensity (classified as 1, 2, and 3) and the percentage of stained cells at each intensity level. For GFAP IHC, the presence of intermixed brain parenchyma within the metastasis was evaluated using the following score: 0: no intermixed GFAP-positive fibrils; 1: focal intermixed GFAP-positive fibrils, 2: multifocal intermixed GFAP-positive fibrils, 3: diffuse intermixed GFAP-positive fibrils. For p53 and p21, the prevalence of positive (nuclear) neoplastic cells was assessed.

**Fatty acid content analysis**. EMT6 cells were seeded in 6 cm plates at a density of 250,000 cells per plate in triplicate. The next day, cells were starved for 4h in SF-AM and then cultured for 20h in fresh ACM or SFM. Cell culture medium was then collected and snap frozen. Cells were trypsinized and $2*10^6$ cells per sample were analyzed. Lipids were extracted according to the procedure of Folch et al., (*131*). The total cell's lipid amount was calculated after aliquot evaporation to constant weight, as previously described (*132*, *133*). Extracted Lipids were processed for the analysis of fatty acids as follows: after extraction and weight, lipid representing 5 ml aliquot of the medium and the pellet corresponding to $2x10^6$ cells, kept frozen at -20ºC before use, were taken into a screw-capped tube (teflon-lined) containing 5µg heptadecanoic acid as internal standard. 1ml B3F in methanol was added for fatty acids esterification. The tubes were gassed with nitrogen and heated at 90ºC for 45 min with occasional shaking. After cooling, 1ml of hexane was added, the tubes' content was mixed and, after a short centrifugation, the hexane layer containing fatty acid methyl ester derivatives (FAME) was transferred into a new tube. The hexane extracts were concentrated by evaporation under nitrogen and one-twentieth of the final suspension was applied in 1µl hexane into the gas chromatograph. FAs were analyzed as methyl ester derivatives (FAME) by gas chromatography (GC) in a Varian, 3800 Series (Walnut Creek, CA) chromatograph (FID) with a fused silica SGE capillary column 30 x 0.025 and Varian Star Workstation Advance Application software, version 6x. The fatty acids' profiles were compared to that of a known mixture of fatty acids of animal source, PUFA2 (Supelco, USA) for identification and their amount quantitated by reference to the internal standard (*134*).

**Metabolomic profiling of astrocyte conditioned medium**. Extraction and analysis of lipids and polar metabolites was performed as previously described (*135*, *136*) with the following modifications: 100µl of medium samples (SFM and ACM) were extracted with 1ml of a pre-cooled (−20˚C) homogenous methanol:methyl-tert-butyl-ether (MTBE) 1:3 (v/v) mixture, containing following internal standards: 0.1µg/ml of phosphatidylcholine (17:0/17:0) (Avanti), 0.4µg/ml of phosphatidylethanolamine (17:0/17:0, 0.15nmol/ml of ceramide/sphingoid internal standard mixture II (Avanti, LM6005), 0.0267µg/ml d5-TG internal standard mixture I (Avanti, LM6000) and 0.1µg/ml palmitic acid-13C (Sigma, 605573). The tubes were vortexed and then sonicated for 30 min in ice-cold sonication bath (taken for a brief vortex every 10 min). Then, UPLC-grade water: methanol (3:1, v/v) solution (0.5ml), containing internal following standards: C13 and N15 labeled amino acids standard mix (Sigma, 767964, 1:500), was added to the tubes followed by vortex and centrifugation. The upper, organic phase, was transferred into 2ml eppendorf tube. The polar phase was re-extracted as described above, with 0.5 ml of MTBE. Both organic phases were combined and dried in speedvac and then stored at −80˚C until analysis. The lower, polar phase, used for polar metabolite analysis was treated in a similar way. The dried lipid extracts were re-suspended in 250µl mobile phase B (see below) and centrifuged at 13,000 rpm and 4˚C for 10 min. Then 225µL were transferred to the HPLC vials for injection. Polar dry samples were re-suspended in 100µl Methanol:DDW (50:50), centrifuged twice to remove the debris. 60µl were transferred to the HPLC vials for injection. LC-MS for lipidomics analysis: Lipid extracts were analyzed using a Waters ACQUITY I class UPLC system coupled to a mass spectrometer (Thermo exactive plus orbitrap) which was operated in switching positive and negative ionization mode. The analysis was performed using Acquity UPLC System combined with chromatographic conditions as described in Malitsky *et al*. (2016) with small alterations. Briefly, the chromatographic separation was performed on an ACQUITY UPLC BEH C8 column (2.1×100mm, i.d., 1.7µm) (Waters Corp., MA, USA). The mobile phase A consisted of DDW: Acetonitrile: Isopropanol 46:38:16 (v/v/v) with 1% 1M $NH_4Ac$, 0.1% acetic acid. Mobile phase B composition is DDW: Acetonitrile: Isopropanol 1:69:30 (v/v/v) with 1% 1M $NH_4Ac$, 0.1% acetic acid. The column was maintained at 40˚C and flow rate of mobile phase was 0.4ml/min. Mobile phase A was run for 1 min at 100%, then it was gradually reduced to 25% at 12 min, following decrease to 0% at 16 min. Then, mobile phase B was run at 100% till 21 min, and mobile phase A was set to 100% at 21.5 min. Finally, column was equilibrated at 100% A till 25 min. Lipid identification and quantification: Orbitrap data was analyzed using LipidSearch™ software (Thermo Fisher Scientific). The validation of the putative identification of lipids was performed by comparing to home-made library which contains lipids produced by various organisms and on the correlation between retention time and carbon chain length and degree of unsaturation. Relative levels of lipids were normalized to the internal standards and the protein amount in the examined samples. LC-MS polar metabolite analysis: Metabolic profiling of polar phase was done as described (*136*) with minor modifications described below. Briefly, analysis was performed using Acquity I class UPLC System combined with mass spectrometer Q Exactive Plus Orbitrap™ (Thermo Fisher Scientific) which was operated in a negative ionization mode. The LC separation was done using the SeQuant Zic-pHilic (150mm × 2. mm) with the SeQuant guard column (20mm × 2.1mm) (Merck). The Mobile phase B: acetonitrile and Mobile phase A: 20mM ammonium carbonate with 0.1% ammonia hydroxide in water: acetonitrile (80:20, v/v). The flow rate was kept at 200µl/min and gradient as follow: 0-2min 75% of B, 14 min 25% of B, 18 min 25% of B, 19 min 75% of B, for 4 min, 23 min 75% of B. Polar metabolites data analysis: The data processing was done using Progenesis QI (Waters), when detected compounds were identified by accurate mass, retention time, isotope pattern, fragments and verified using in-house-generated mass spectra library.

**Genome-wide RNAi, CRISPR and drug screens.** The differential sensitivities of *TP53*-WT and *TP53*-null BC cell lines in genome-wide RNAi screens were compared using gene-set enrichment analysis (GSEA). To this end, the top 500 significant hits were analyzed for enrichment of canonical pathways using the MsigDb database. RNAi and CRISPR sensitivity values for specific genes were correlated with ssGSEA scores for the MSigDB gene signature ‘KANNAN_p53_targets_DN as an approximation for p53 pathway activity. Drug sensitivities were similarly compared between *TP53*-WT and *TP53*-null human BC cell lines.

**Drug treatments.** Drug treatments were performed as previously described (*137*). For drug treatments, cancer cells were seeded in 96-well plates at 2000 cells/well (EMT6) or 4000 cells/well (CAL51) in 3-5 technical replicates. The next day, culture medium was replaced by fresh medium containing the drug, or a medium control that was adjusted for the drug solvent. Cell viability and proliferation were followed by live-cell imaging using a Livecyte or Incucyte, as described above, and cell viability was determined using the MTT assay at endpoint. Cell culture medium was replaced by fresh MTT solution (Sigma M2128, 0.5mg/ml) and incubated in a cell culture incubator until purple crystals within cells were clearly visible under a dissecting microscope. The reaction was stopped and MTT crystals were extracted with 10% Triton X-100 and 0.1N HCl in isopropanol and vigorous pipetting. Color absorption was quantified at 570nm and 630nm using a plate-reader ((BioTek Synergy H1 hybrid multi-mode reader, Agilent).

The following drugs and dilutions were used: SCD1 inhibitors SW203668(*138*) (#19379, Cayman) was prepared as a 10mM Stock in DMSO, concentrations between 3µM and 0.1µM were used for drug treatments. The drug was replaced every 24h for the duration of the treatment. SCD1 inhibitor A939572 (#1939572, Cayman) was prepared as a 1mg/ml Stock solution in DMSO and used as a working concentration of 75µM as previously described (*139*, *140*). The FASN inhibitor C75 (*141–143*) (#10005270, Cayman) was prepared as a Stock in DMSO at 20mM and used at a working concentration of 10-25µM. For Oleic acid treatments, Oleic acid (*144*) (#90260, Cayman) was prepared as a 5mM stock and used at a final concentration of 2.5-50µM. Glutamate (G1626-100G, Sigma) was used at a 75mM working concentration in glutamine-free astrocyte medium (ScienCell; 1801-NG).

**Multicellular 3D tumor spheroids**. Following brain tumor cell isolation on day 15, cell suspensions mainly containing GFP-labeled EMT6 cells (*Trp53*-WT or *Trp53*-null) were seeded (500 cells/well) into a U-shape 96-well plate (Corning) and incubated for 3 days. Cells were then treated with serial dilutions of SCD1 inhibitor (SW203668) (*138*), dilution 1:3 starting from 3µM, and the GFP signal fluorescence was monitored every 3h and imaged using a 10X objective. Results were calculated by the IncuCyte™ Software and presented as % of total red area covered (image/well) normalized to time zero for each condition.

**Statistical analyses.** Statistical analyses were performed using Microsoft excel and GraphPad PRISM® 9.1. Details of each statistical test, the number of analyzed samples and the number of independent experiments are specified in the Figure legends. Significance is indicated as customary: *p<0.05, ** p<0.01, *** p<0.001, ****<$1*10^{-4}$. Statistical significance between two groups was calculated by a two-tailed or one-tailed Student's t-test as appropriate. Paired student's t-test was performed for patient-matched expression analysis and bilateral intracranial injections that contained matched *Trp53*-WT and *Trp53*-null cells. The Mann-Whitney test was used to compare two groups when one or both groups showed a data distribution that was not normal. Two-sided Chi-square or Fisher's Exact test (when sample numbers were limited) were used to determine the significance of differences in the prevalence of events between two groups. Significance of GSEA was based on normalized enrichment score (NES) and the false discovery rate q-value (FDR q-value) as calculated by the GSEA software (version 4.03). Significance of ssGSEA was determined through a two-sided or one-sided Student's t-test between two groups as appropriate. Q-values for ssGSEA results were computed using the Benjamini–Hochberg method. Correlations and linear regression were calculated using GraphPad PRISM 9.1. Survival curves were calculated using the Kaplan Meyer method with GraphPad PRISM 9.1. Significance was determined with the Gehan-Breslow-Wilcoxon test, which prioritizes early events. Generally, a one-tailed statistical test was chosen when the directionality of the experiment was predicted, based on previous experimental results with a different data set or cell line.

In each presented box plot, all data points are shown and the internal line represents the median of the distribution. The box extends from the $25^{th}$ to $75^{th}$ percentiles. Whiskers extend to the highest and lowest data point.

**Supplementary figures:**

Fig.S1

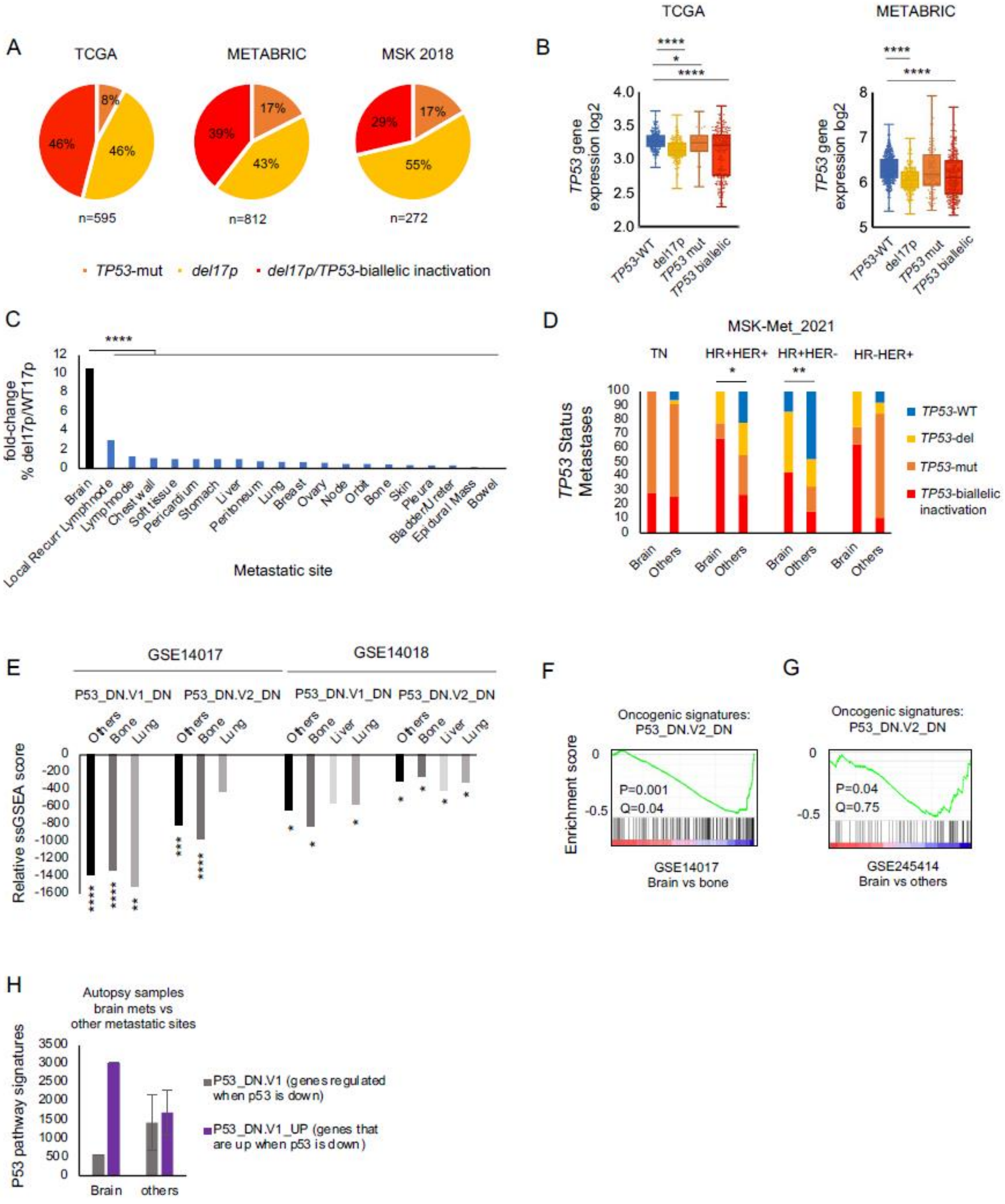

**Fig. S1: Clinical data analysis of *TP53* perturbation in BCBM (related to Fig. 1).**

**A,** Distribution of different types of *TP53* inactivation in primary breast tumors using data from TCGA, METABRIC and MSK_2018. Most tumors with *TP53* inactivating events lost a copy of *TP53* through deletion of chromosome 17p (del17p). Breast tumors with altered Chromosome 17p in %: TCGA 92%, METABRIC 82%, MSK_2018 84%.

**B,** *TP53* gene expression analysis. *TP53* gene expression is lower in breast tumors with *TP53* copy number alterations (with or without additional point mutations) in TCGA (left) and METABRIC (right) datasets. TCGA: *TP53*-WT vs del17p, $p=2.7*10^{-22}$; *TP53*-WT vs *TP53*-mut, p=0.02; *TP53*-WT vs biallelic inactivation, $p=2.2*10^{-12}$; *TP53*-WT n=259, del17p n=274, *TP53*-mut n=47, *TP53* biallelic inactivation n=247. METABRIC: *TP53*-WT vs del17p, $p=1.17*10^{-25}$; *TP53*-WT vs *TP53*-mut, p=0.29; *TP53*-WT vs biallelic inactivation, $p=2.4*10^{-10}$; *TP53*-WT n=592, del17p n=280, *TP53*-mut n=142, *TP53* biallelic inactivation n=320.

**C,** Prevalence of del17p in metastases from different metastatic sites shown as fold-change over WT17p tumors using data from Razavi et al., (*17*), Brain compared to other metastatic sites, Fisher's exact test, $p=1*10^{-5}$; Brain n=23, local recurrence n=4, lymph node n=83, chest wall n=56, soft tissue n=14, pericardium n=2, stomach n=2, lung n=24, breast n=10, ovary n=27, orbit n=3, bone n=98, skin n=15, pleura n=20, bladder/ureter n=4, epidural Mass n=7, bowel n=7.

**D,** Comparison of the prevalence of *TP53* alterations in BM and other metastatic sites originating from breast tumors of different subtypes: Triple-Negative (TN), HR+HER+, HR+HER-, HR-HER+. *TP53* inactivation vs *TP53*-WT, brain vs others, one-tailed Fisher's exact test, TN p=0.5, HR+HER+ p=0.18, HR+HER-p=0.05, HR- HER+ p=0.5, *TP53* biallelic inactivation vs *TP53*-WT, brain vs others, one-tailed Fisher-Exact test TN p=0.5, HR+HER+ p=0.04, HR+HER- p=0.002, HR- HER+ p=0.1. Sample number TN: Brain n=7, others n=136, HR+HER+: Brain n=9, others n=100, HR+HER-: Brain n=14, others n=569, HR- HER+: Brain n=8, others n=38.

**E,** Single-sample gene set enrichment analysis (GSEA) comparing brain metastases to other metastatic sites. p53 activity is lower in brain as compared to other metastatic sites (GSE14017 and GSE14018), supplementing Fig. 1F,G. GSE14017: P53_DN.V1_DN,
Brain vs others, $p=8*10^{-5}$; Brain vs bone, $p=7*10^{-4}$; Brain vs lung, p=0.002. P53_DN.V2_DN Brain vs others, $p=4*10^{-4}$; Brain vs bone, $p=9.8*10^{-5}$; Brain vs lung, p=0.22. GSE14018: GSEA signature P53_DN.V1_DN, Brain vs others, p=0.02; Brain vs bone, p=0.01; Brain vs liver, p=0.18; Brain vs lung, p=0.01. P53_DN.V2_DN Brain vs others, p=0.01; Brain vs bone p=0.05; Brain vs

liver p=0,04; Brain vs lung, p=0.04. GSE14017:  Brain n=15, bone n=10, lung n=4; GSE14018: Brain n=7, bone n=9, liver n=5, lung n=16.

**F,** GSEA plot showing lower TP53 target gene expression in brain when compared to bone mets. Oncogenic signature P53_DN.V2_DN p<0.001, q=0.04. Brain n=15, bone n=10. Table S1.

**G,** Analysis of a third, independent dataset (GSE245414), comparing gene expression profiles between brain metastatic tissue and other metastatic sites, confirms lower p53 activity in BM. GSEA Brain vs others, oncogenic signatures P53_DN.V2_DN p<0.04. ssGSEA p=0.01. Brain n=6, others n= 26 (lymph nodes n=5, liver n=1, lung n=3, pleura n=3, soft tissue n=8, spine n=6.) Table S1.

**H,** Single sample gene set enrichment analysis (single sample GSEA**)** comparing gene expression profiles across autopsy samples from different metastatic sites taken from the same patient. Gene expression signatures associated with the p53 pathway were lower in brain metastatic tissue relative to other metastatic samples. Gene expression signatures P53_DN.V1 and P53_DN.V1_UP; Brain n=1, others n=6.

Fig.S2

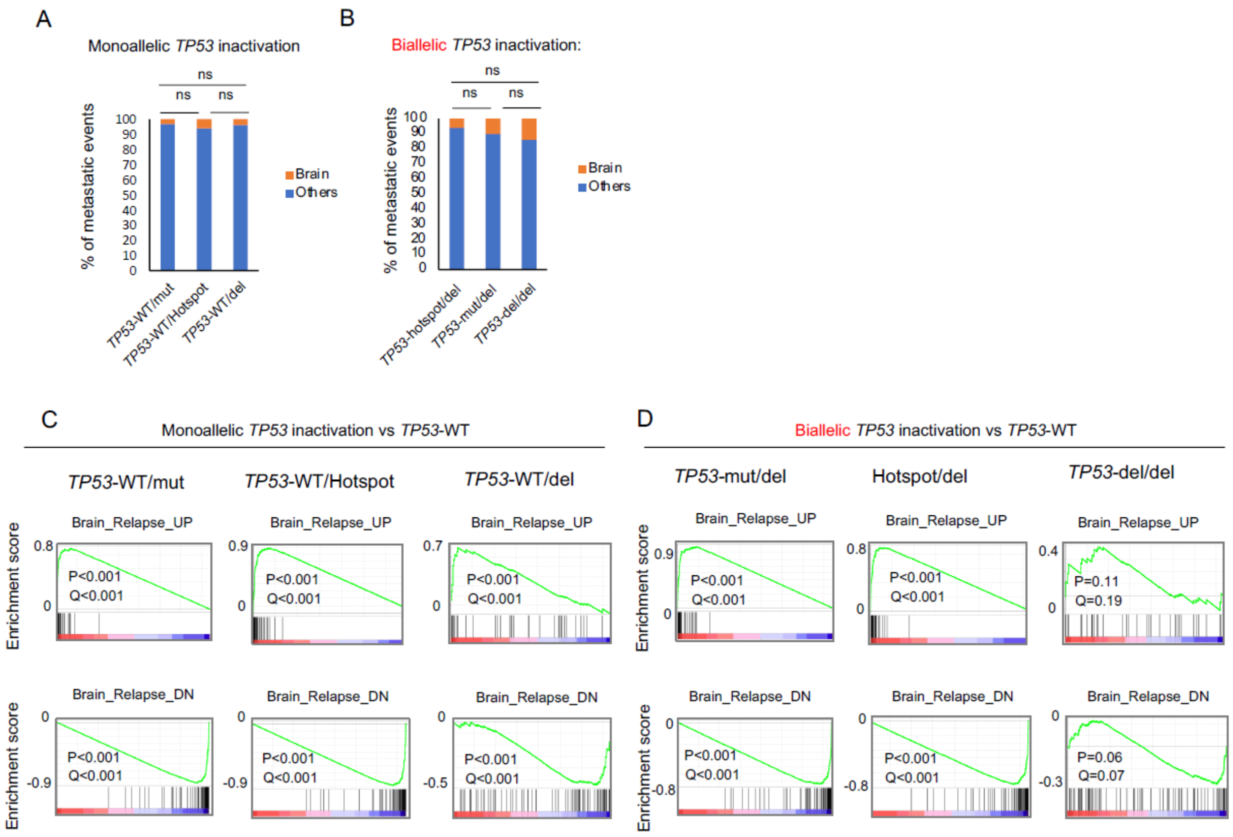

A
Monoallelic TP53 inactivation
% of metastatic events
ns
Brain
Others
TP53-WT/mut
TP53-WT/Hotspot
TP53-WT/del
B
Biallelic TP53 inactivation:
TP53-hotspot/del
TP53-mut/del
TP53-del/del
C
Monoallelic TP53 inactivation vs TP53-WT
D
Biallelic TP53 inactivation vs TP53-WT
TP53-mut/del
Hotspot/del
TP53-del/del
Brain_Relapse_UP
Brain_Relapse_DN
Enrichment score
P<0.001
Q<0.001
P=0.11
Q=0.19
P=0.06
Q=0.07

**Fig. S2: The association between *TP53* perturbation and BM is not due to specific hotspot mutations (related to Fig. 1).**

**A,B,** No difference in BM prevalence between different modes of (a) monoallelic and (b) biallelic *TP53* inactivation. *TP53* mutants with 3 common mutations, *TP53* mutants with all other mutations, and del17p tumors (with no known other *TP53* mutations) (data set MSK_Met_2021). Mutations were considered hotspot mutations if they involved the following transitions: R175H, R275H, R243any.

Monoallelic *TP53*-inactivation: *TP53*-WT/mut vs WT/Hotspot p= 0.23, TP53-WT/Mut vs *TP53*-WT/del p= 0.06, *TP53*-WT/ Hotspot vs *TP53*-WT/del= 1. Sample number: *TP53*-WT/mut n=218, *TP53*-WT/hotspot n=33, *TP53*-WT/del n=152.

Biallelic *TP53* inactivation: *TP53*-mut/del vs *TP53*-hotspot/del p=0.37, *TP53*-mut/del vs *TP53*-del/del p=0.64, *TP53*-hotspot/del vs del/del p=1. Sample number *TP53*-mut/del n=150, *TP53*-hotspot/del n=12, *TP53*-del/del n=14.

**C,D** Gene set enrichment analysis (GSEA). BM gene expression profiles were compared in tumors where *TP53* was altered through *TP53* hotspot mutations and/or deletion events. GSEA showed the same trend in c) monoallelic *TP53* inactivation vs *TP53*-WT and d) biallelic *TP53* inactivation vs *TP53*-WT independent of the mode of p53 inactivation. *TP53*-WT n=693, *TP53*-WT/mut n=168, *TP53*-WT/Hotspot n=30, *TP53*-WT/del n=505, *TP53*-mut/del n=405, *TP53*-Hotspot/del n=60, *TP53*-del/del n=5

Fig.S3

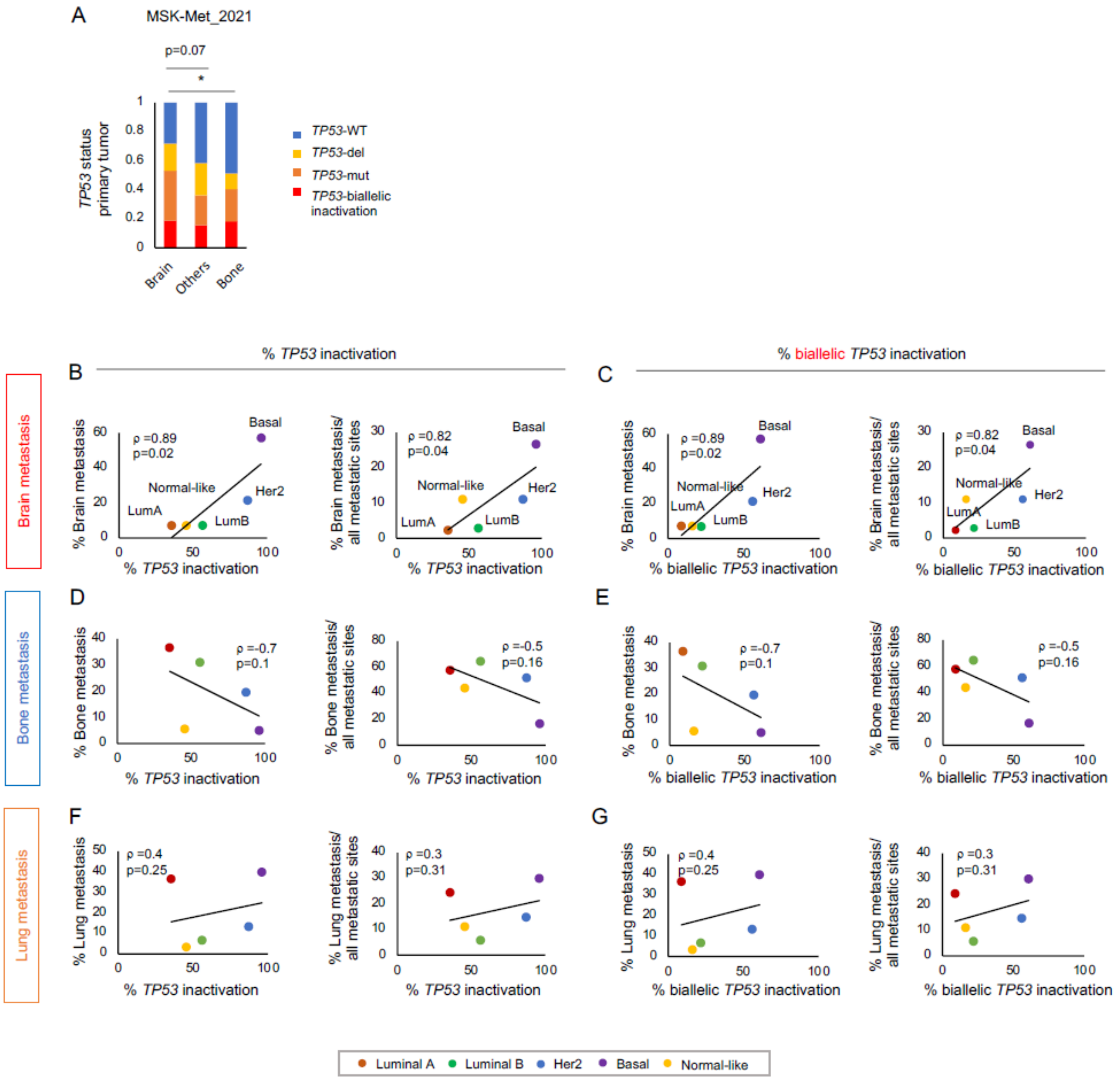
A
MSK-Met_2021
p=0.07
*
TP53 status primary tumor
Brain
Others
Bone
TP53-WT
TP53-del
TP53-mut
TP53-biallelic inactivation
% TP53 inactivation
% biallelic TP53 inactivation
B
C
D
E
F
G
Brain metastasis
Bone metastasis
Lung metastasis
% Brain metastasis
% Brain metastasis/ all metastatic sites
% Bone metastasis
% Bone metastasis/ all metastatic sites
% Lung metastasis
% Lung metastasis/ all metastatic sites
ρ =0.89
p=0.02
ρ =0.82
p=0.04
ρ =-0.7
p=0.1
ρ =-0.5
p=0.16
ρ =0.4
p=0.25
ρ =0.3
p=0.31
Basal
Her2
LumB
LumA
Normal-like
Luminal A
Luminal B
Her2
Basal
Normal-like

**Fig. S3: Prevalence of *p53* inactivation is associated with metastasis to the brain, but not to bone or lung, across BC subtypes (related to Fig. 1).**

**A,** Frequency of *TP53* alterations in primary breast tumors that metastasize to the brain compared to other metastatic sites. Data set used: MSK-Met 2021. *TP53*-inactivation vs *TP53*-WT, one-tailed Fisher's exact test, brain vs others p=0.07. Brain vs bone p=0.02, brain n=32, others n=958, bone n=235. (Tumors that metastasized to >4 sites were excluded from the analysis.) **B,C** *TP53* inactivation in BC subtypes. Significant positive correlation for *TP53* inactivation with BM occurrence across BC subtypes. Data source for *TP53* inactivation prevalence: METABRIC; data source for BM prevalence: Smid et al., 2008(*118*). **b,** Percentage of *TP53* inactivation. Left: Spearman's correlation: % all *TP53* inactivation with BM prevalence in %, rho= 0.89, p=0.02. Right: Spearman's correlation: % all TP53 inactivation with % BM / all metastasis of same subtype in %, rho = 0.82, one-tailed p=0.04 **c,** Percentage of *TP53* biallelic inactivation. Spearman's correlation: % all *TP53* inactivation with BM prevalence in %, rho= 0.89, p=0.02. Spearman's correlation: % all *TP53* inactivation with percent BM/ all metastasis of same subtype in %, rho= 0.82, one-tailed p =0.04.

**D-G,** No significant correlation of *TP53* inactivation and the metastasis to other metastatic sites, as shown for two common metastatic sites of BC, bone and lung.

**D,E** Trending negative correlation for the presence of *TP53* inactivation events with bone metastasis prevalence. **D,** Percent *TP53* inactivation. Spearman's correlation: % all *TP53* inactivation with bone metastasis prevalence in %, rho = -0.7, p=0.1. Spearman's correlation: % all *TP53* inactivation with % bone metastasis/ all metastasis of same subtype in %, rho= 0.5, p =0.16. **E,** Trending negative correlation of percentage *TP53* biallelic inactivation with bone metastasis prevalence in %, rho=-0.7, p =0.1. Spearman's correlation: % *TP53* biallelic inactivation with percent bone metastasis/ all metastasis of same subtype in %, rho =-0.5, p =0.16. **F,G,** No significant correlation between loss of *TP53* and lung metastasis prevalence. Spearman's correlation: % all *TP53* inactivation with lung metastasis prevalence in %, rho= 0.4, p=0.25. Spearman's correlation: % all *TP53* inactivation with percent lung metastasis/ all metastasis of same subtype in %, rho=0.31, p=0.3. **G,** percent *TP53* biallelic inactivation. Spearman's correlation: % all p53 inactivation with lung metastasis prevalence in %, rho = 0.4, p =0.25. Spearman's correlation: % all *TP53* inactivation with percent lung metastasis/ all metastasis of same subtype in %, rho= 0.31, p =0.3. (Note that the Sperman correlation coefficients and p-values are identical in panels ‘B’ vs. ‘C’, ‘D’ vs. ‘E’, and ‘F’ vs. ‘G’, due to the same ranking of observations.)

### Fig.S4

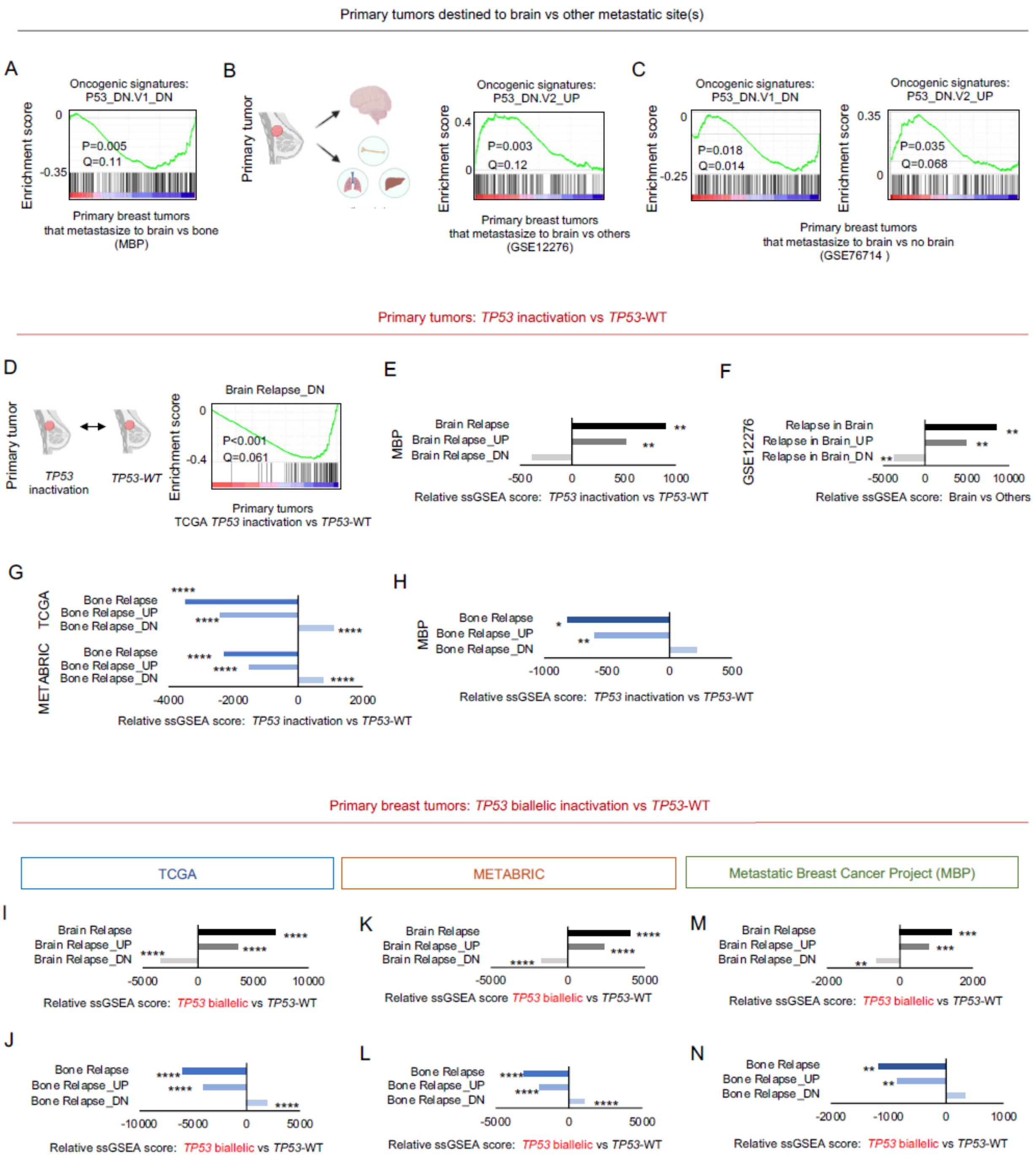

Primary tumors destined to brain vs other metastatic site(s)
A
Oncogenic signatures: P53_DN.V1_DN
Enrichment score
0
-0.35
P=0.005
Q=0.11
Primary breast tumors that metastasize to brain vs bone (MBP)
B
Primary tumor
Oncogenic signatures: P53_DN.V2_UP
0.4
0
P=0.003
Q=0.12
Primary breast tumors that metastasize to brain vs others (GSE12276)
C
Oncogenic signatures: P53_DN.V1_DN
-0.25
P=0.018
Q=0.014
Oncogenic signatures: P53_DN.V2_UP
0.35
P=0.035
Q=0.068
Primary breast tumors that metastasize to brain vs no brain (GSE76714 )
Primary tumors: TP53 inactivation vs TP53-WT
D
TP53 inactivation
TP53-WT
Brain Relapse_DN
-0.4
P<0.001
Q=0.061
Primary tumors TCGA TP53 inactivation vs TP53-WT
E
MBP
Brain Relapse
Brain Relapse_UP
Brain Relapse_DN
-500 0 500 1000
Relative ssGSEA score: TP53 inactivation vs TP53-WT
F
GSE12276
Relapse in Brain
Relapse in Brain_UP
Relapse in Brain_DN
-5000 0 5000 10000
Relative ssGSEA score: Brain vs Others
G
TCGA
METABRIC
Bone Relapse
Bone Relapse_UP
Bone Relapse_DN
-4000 -2000 0 2000
Relative ssGSEA score: TP53 inactivation vs TP53-WT
H
MBP
Bone Relapse
Bone Relapse_UP
Bone Relapse_DN
-1000 -500 0 500
Relative ssGSEA score: TP53 inactivation vs TP53-WT
Primary breast tumors: TP53 biallelic inactivation vs TP53-WT
TCGA
METABRIC
Metastatic Breast Cancer Project (MBP)
I
Brain Relapse
Brain Relapse_UP
Brain Relapse_DN
-5000 0 5000 10000
Relative ssGSEA score: TP53 biallelic vs TP53-WT
K
-5000 0 5000
Relative ssGSEA score TP53 biallelic vs TP53-WT
M
-2000 0 2000
Relative ssGSEA score: TP53 biallelic vs TP53-WT
J
Bone Relapse
Bone Relapse_UP
Bone Relapse_DN
-10000 -5000 0 5000
Relative ssGSEA score: TP53 biallelic vs TP53-WT
L
-5000 0 5000
Relative ssGSEA score: TP53 biallelic vs TP53-WT
N
-2000 -1000 0 1000
Relative ssGSEA score: TP53 biallelic vs TP53-WT

**Fig. S4: Gene set enrichment analysis of gene expression data from primary human breast tumors (related to Fig. 1).**

**A-C,** Gene set enrichment analysis (GSEA) of datasets comparing gene profiles between primary tumors of patients who developed brain metastases and those who developed metastases in other organs.

**A**, GSEA plot showing downregulation of a p53 expression signature in primary tumors that metastasized to the brain as compared to bone, supplementing plot shown in Fig. 1H (data source: Metastatic breast cancer project (MBP). Oncogenic signatures p53_DN.V1_DN, brain vs bone p=0.005, brain n=7, bone=24. Table S2.

**B,** GSEA plot showing upregulation of a signature indicative of lower p53 signaling in primary breast tumors that metastasized to the brain in comparison to other sites in a second dataset (GSE12276). Brain vs others, oncogenic signatures P53_DN.V2_UP, p=0.003, brain n=7, others n=121. Table S2.

**C,** GSEA plot showing downregulation of a signature indicative of increased p53 signaling (left) and upregulation of a signature indicative of decreased p53 signaling (right) in triple-negative primary breast tumors that metastasized to the brain vs. those that did not (data source GSE76714, Duchnowska, Breast Cancer 2017). Brain vs others, Oncogenic signatures: P53_DN.V1_DN p=0.018, Oncogenic signatures: P53_DN.V2_UP p= 0.035, brain n=21, others n=38.

**D,** GSEA of primary breast carcinomas from TCGA, comparing tumors with *TP53* inactivation to *TP53*-WT tumors. SMID_BREAST_CANCER_RELAPSE_IN_BRAIN_DN p<0.001. *TP53* inactivation n=623, *TP53*-WT n=259.

**E,** ssGSEA of primary breast carcinomas comparing tumors with *TP53* inactivation to *TP53*-WT tumors in a third dataset, supporting the data shown in Fig. 1M. Tumors with *TP53* inactivation showed increased expression of signatures positively correlated with BM. *TP53* inactivation vs *TP53*-WT, The metastatic breast cancer project MBP, SMID_BREAST_CANCER_RELAPSE_IN_BRAIN p=0.01, _UP p=0.003, _DN p=0.07. Table S4.

**F,** ssGSEA validation of MsigDB brain relapse signature using primary breast tumors that metastasized to brain compared to those that metastasize elsewhere. GSE12276. Brain vs others, SMID_BREAST_CANCER_RELAPSE_IN_BRAIN p= 0.007, ranked 1, SMID_BREAST_CANCER_RELAPSE_IN_BRAIN_UP p=0.008 ranked 8,

SMID_BREAST_CANCER_RELAPSE_IN_BRAIN_DN p=0.01 ranked 8, brain n=7, others n=121. Table S4.

**G,H,** ssGSEA analysis comparing human primary breast carcinomas that carry *TP53* inactivation with *TP53*-WT tumors, supplementing Fig. 1M. Gene expression signatures associated with bone metastasis are decreased in primary tumors that harbor *TP53* alterations. Note this is the opposite trend observed with BM associated gene signatures. *TP53* inactivation vs *TP53*-WT, TCGA, SMID_BREAST_CANCER_RELAPSE_IN_BONE $P=1.8*10^{-42}$, _UP $P=9.6*10^{-53}$, _DN $P=5.9*10^{-54}$; METABRIC, SMID_BREAST_CANCER_RELAPSE_IN_BONE $P=1.2*10^{-19}$, _UP $P=4.4*10^{-25}$, _DN $P=2.6*10^{-24}$ .MBP, **S**MID_BREAST_CANCER_RELAPSE_IN_BONE p=0.03, _UP p=0.008, _DN p=0.23.

**I-N,** ssGSEA. Comparison of primary BCs with *TP53* biallelic inactivation to tumors that are *TP53*-WT using expression data from TCGA (**I,J**), METABRIC (**K,L**) and MBP(**M,N**) cohorts. Gene expression signatures compared are associated with BM and colored in grey or associated with bone metastasis shown in shades of blue. Note opposing trends. Table S3.

TCGA: SMID_BREAST_CANCER_RELAPSE_IN_BRAIN $p=9.7*10^{-50}$, _UP $p=2*10^{-46}$,_DN $p=3.9*10^{-48}$.SMID_BREAST_CANCER_RELAPSE_IN_BONE $p=9.3*10^{-52}$, _UP $P=2.6*10^{-53}$_DN, $p=6*10^{-42}$.

METABRIC: SMID_BREAST_CANCER_RELAPSE_IN_BRAIN $p= 2.3*10^{-74}$, BRAIN _UP $p= 1.7*10^{-76}$, BRAIN _DN $p=1.4*10^{-58}$, SMID_BREAST_CANCER_RELAPSE_IN_BONE $p=6.5*10^{-68}$, BONE _UP $P=1*10^{-65}$, BONE _DN $p=7*10^{-53}$.

MBP: SMID_BREAST_CANCER_RELAPSE_IN_BRAIN $p= 7*10^{-4}$, BRAIN _UP $p=2*10^{-4}$, BRAIN _DN p=0.01, SMID_BREAST_CANCER_RELAPSE_IN_BONE p=0.004, BONE _UP p=0.002, BONE _DN p=0.11.

Fig.S5

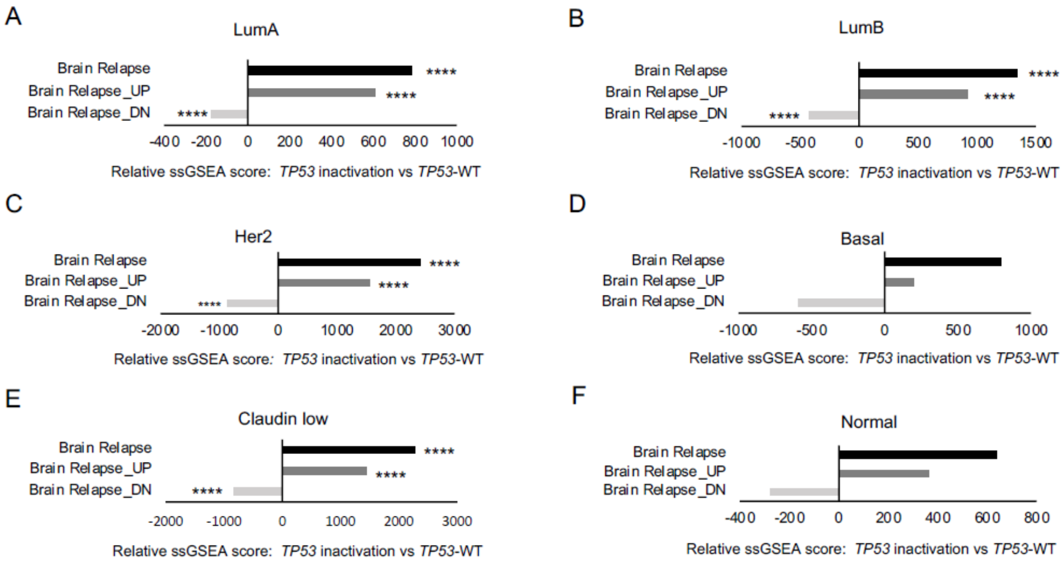
A
LumA
Brain Relapse
Brain Relapse_UP
Brain Relapse_DN
****
****
****
-400 -200 0 200 400 600 800 1000
Relative ssGSEA score: TP53 inactivation vs TP53-WT
B
LumB
Brain Relapse
Brain Relapse_UP
Brain Relapse_DN
****
****
****
-1000 -500 0 500 1000 1500
Relative ssGSEA score: TP53 inactivation vs TP53-WT
C
Her2
Brain Relapse
Brain Relapse_UP
Brain Relapse_DN
****
****
****
-2000 -1000 0 1000 2000 3000
Relative ssGSEA score: TP53 inactivation vs TP53-WT
D
Basal
Brain Relapse
Brain Relapse_UP
Brain Relapse_DN
-1000 -500 0 500 1000
Relative ssGSEA score: TP53 inactivation vs TP53-WT
E
Claudin low
Brain Relapse
Brain Relapse_UP
Brain Relapse_DN
****
****
****
-2000 -1000 0 1000 2000 3000
Relative ssGSEA score: TP53 inactivation vs TP53-WT
F
Normal
Brain Relapse
Brain Relapse_UP
Brain Relapse_DN
-400 -200 0 200 400 600 800
Relative ssGSEA score: TP53 inactivation vs TP53-WT

**Fig. S5: BM signatures are elevated in *TP53*-deficient primary BCs across all molecular subtypes (related to Fig. 1).**

**A,** Gene set enrichment analysis comparing primary BCs that carried any *TP53* perturbation to *TP53*-WT tumors. Tumors were stratified by subtype to rule out subtype specific effects. BM signatures were regulated consistent with higher brain metastatic capacity of *TP53*-deficient tumors across all subtypes.

METABRIC, *TP53* inactivation vs *TP53*-WT, two-sided Student's t-test,

LumA: SMID_BREAST_CANCER_RELAPSE_IN_BRAIN $p=9.7*10^{-11}$, _UP $p=3.7*10^{-10}$, DN $p= 6.4*10^{-16}$;

LumB: SMID_BREAST_CANCER_RELAPSE_IN_BRAIN $p=2.3*10^{-13}$, _UP $p=2.5*10^{-12}$,_DN $p=7.5*10^{-10}$;

Her2: SMID_BREAST_CANCER_RELAPSE_IN_BRAIN $p=5.9*10^{-6}$ , _UP $p=5.8*10^{-6}$ ,_DN $p=4*10^{-4}$;

Basal: SMID_BREAST_CANCER_RELAPSE_IN_BRAIN p=0.4, _UP p=0.6,_DN p=0.3;

Claudin-low: SMID_BREAST_CANCER_RELAPSE_IN_BRAIN $p=9.6*10^{-7}$, _UP $p=4*10^{-9}$,_DN p=0.001;

Normal: SMID_BREAST_CANCER_RELAPSE_IN_BRAIN p=0.1, _UP p= 0.14,_DN p=0.13.

Number of tumors compared in analysis: LumA: *TP53* inactivated n=177, *TP53*-WT n=323 LumB: *TP53* inactivated n=163, *TP53*-WT n=128 Her2: *TP53* inactivated n=120, *TP53*-WT n=18 Basal: *TP53* inactivated n=142, *TP53*-WT n=6 Claudin-low: *TP53* inactivated n=94, *TP53*-WT n=62 Normal: *TP53* inactivated n=45, *TP53*-WT n=54.

Fig.S6

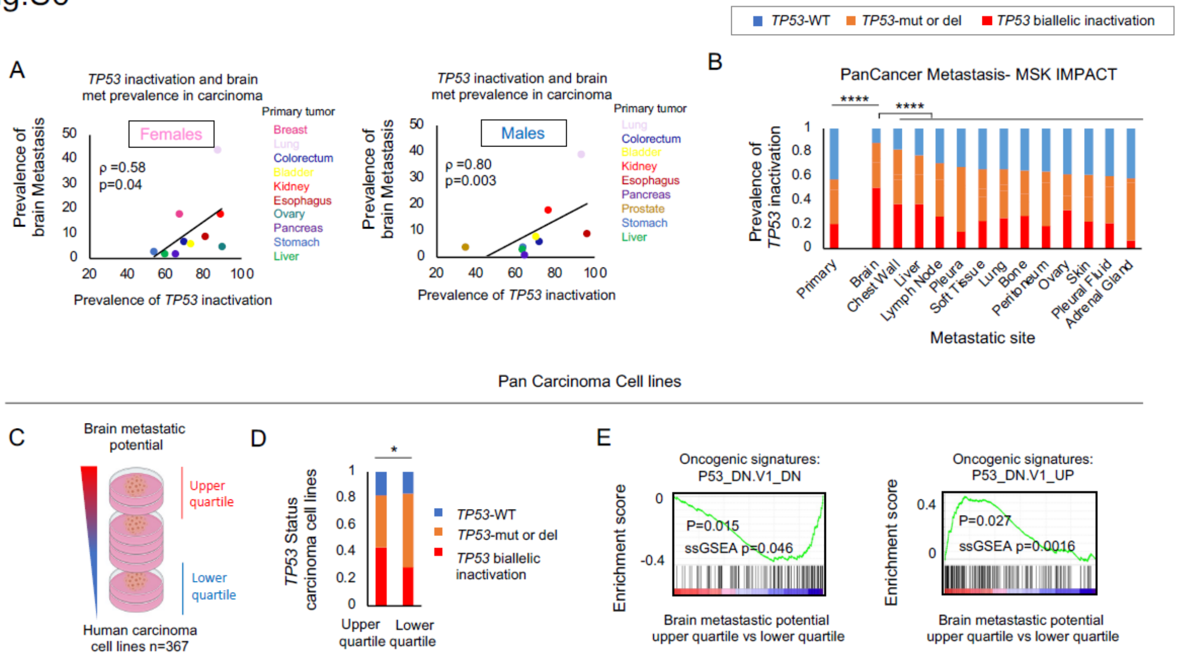

TP53-WT
TP53-mut or del
TP53 biallelic inactivation
A
TP53 inactivation and brain met prevalence in carcinoma
Females
Males
Primary tumor
Prevalence of brain Metastasis
Prevalence of TP53 inactivation
ρ =0.58
p=0.04
ρ =0.80
p=0.003
Breast
Lung
Colorectum
Bladder
Kidney
Esophagus
Ovary
Pancreas
Prostate
Stomach
Liver
B
PanCancer Metastasis- MSK IMPACT
Prevalence of TP53 inactivation
Primary
Brain
Chest Wall
Liver
Lymph Node
Pleura
Soft Tissue
Lung
Bone
Peritoneum
Ovary
Skin
Pleural Fluid
Adrenal Gland
Metastatic site
Pan Carcinoma Cell lines
C
Brain metastatic potential
Upper quartile
Lower quartile
Human carcinoma cell lines n=367
D
TP53 Status carcinoma cell lines
Upper quartile
Lower quartile
TP53 biallelic inactivation
E
Oncogenic signatures: P53_DN.V1_DN
Oncogenic signatures: P53_DN.V1_UP
Enrichment score
P=0.015
ssGSEA p=0.046
P=0.027
ssGSEA p=0.0016
Brain metastatic potential upper quartile vs lower quartile

**Fig. S6: *TP53* inactivation is associated with BM across primary carcinomas (related to Fig. 1).**

**A,** Correlation of *TP53* inactivation with BM prevalence across carcinomas, for female (left) and male (right) tumors. Each dot represents a primary carcinoma (for lung and kidney, the highest-ranking subtypes were considered). Female: one-tailed Spearman's correlation rho = 0.58, p= 0.04. male: one-tailed Spearman's correlation: rho=0.80, p= 0.003.

**B,** Prevalence of *TP53* alterations in metastatic sites in a pan-cancer analysis, using data from MSK-IMPACT(*37*). Brain Metastases are significantly enriched for *TP53* alterations, and for biallelic *TP53* inactivation, in comparison to primary tumors and to other metastatic sites. *TP53* inactivation vs *TP53*-WT: BM vs primary tumor, Fisher's exact test $p<1*10^{-5}$, BM vs other metastasis, Fisher's exact test p=0. *TP53* biallelic inactivation vs *TP53*-WT; BM vs primary tumor, Fisher's exact test, $p<1*10^{-5}$, BM vs other metastasis $p<1*10^{-5}$. Primary n=2947, brain n=126, chest wall n=79, liver n=658, lymph node n=571, pleura n=111, soft tissue n=80, lung n=265, bone n=252, peritoneum n=48, ovary, skin n=47, pleural fluid n=33, adrenal gland n=34.

**C,** 367 carcinoma cell lines were grouped by their brain metastatic potential, and *TP53* status and activity were compared between the groups.

**D,** Comparison of the *TP53* status in the carcinoma cell lines with the highest brain metastatic potential (top 25%) to those with the lowest brain metastatic potential (bottom 25%) shows a significant enrichment of *TP53* biallelic inactivation in cell lines with high potential to metastasize to the brain, p=0.03.

**E,** GSEA of gene expression profiles from highly brain metastatic carcinoma cells (top 25%) compared to carcinoma cells with low brain metastatic potential (bottom 25%) shows decreased p53 signaling in brain metastatic cells. Brain metastastic potential upper quartile vs lower quartile, Oncogenic signatures: P53_DN.V1_DN, p=0.015, ranked 3. P53_DN.V1_UP p=0.027, ranked 4. Table S2.

### Fig.S7

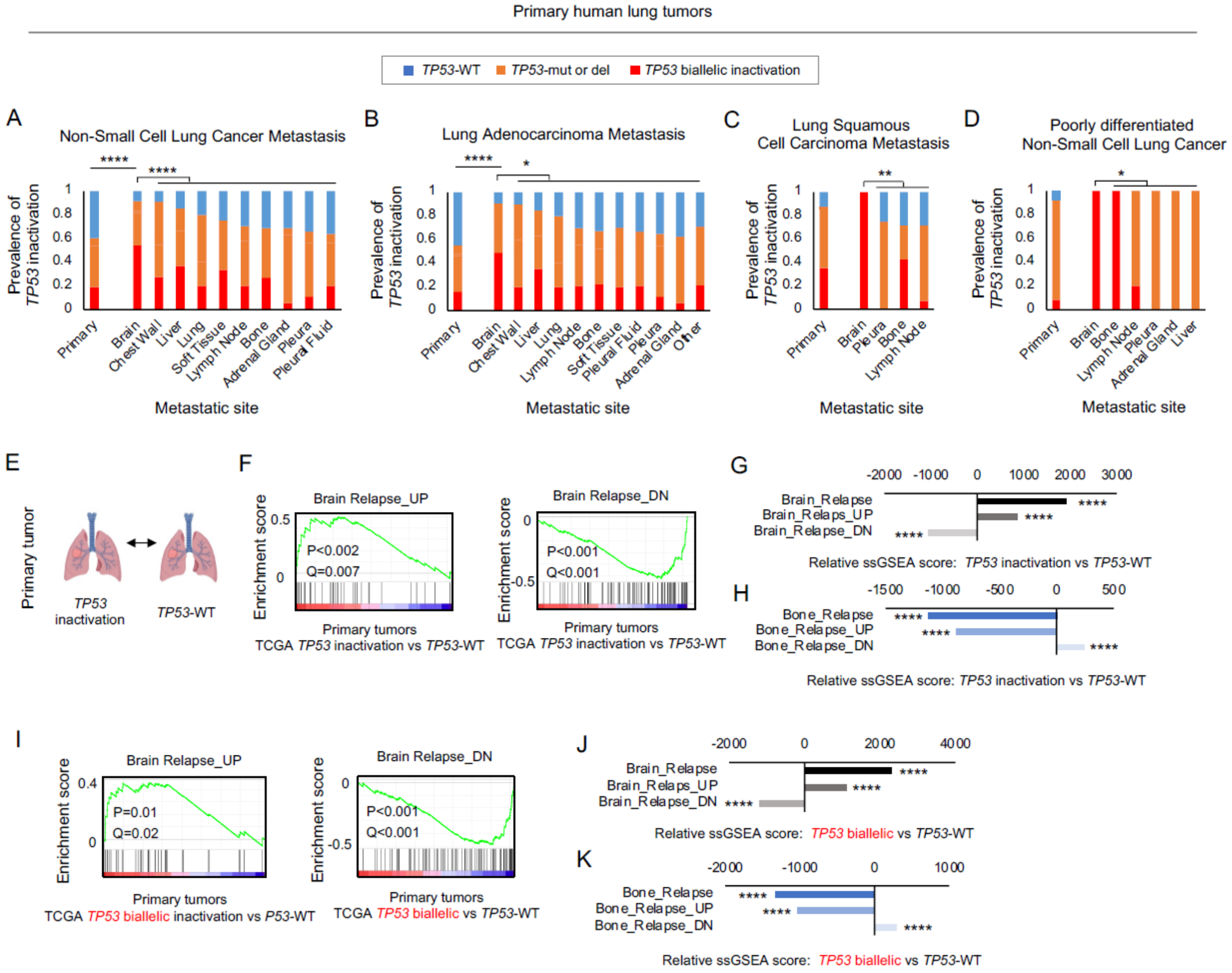

Primary human lung tumors
TP53-WT
TP53-mut or del
TP53 biallelic inactivation
A
Non-Small Cell Lung Cancer Metastasis
Prevalence of TP53 inactivation
Metastatic site
Primary
Brain
Chest Wall
Liver
Lung
Soft Tissue
Lymph Node
Bone
Adrenal Gland
Pleura
Pleural Fluid
B
Lung Adenocarcinoma Metastasis
Other
C
Lung Squamous Cell Carcinoma Metastasis
D
Poorly differentiated Non-Small Cell Lung Cancer
E
Primary tumor
TP53 inactivation
TP53-WT
F
Brain Relapse_UP
P<0.002
Q=0.007
Brain Relapse_DN
P<0.001
Q<0.001
Enrichment score
Primary tumors
TCGA TP53 inactivation vs TP53-WT
G
Brain_Relapse
Brain_Relaps_UP
Brain_Relapse_DN
Relative ssGSEA score: TP53 inactivation vs TP53-WT
H
Bone_Relapse
Bone_Relapse_UP
Bone_Relapse_DN
Relative ssGSEA score: TP53 inactivation vs TP53-WT
I
Brain Relapse_UP
P=0.01
Q=0.02
Brain Relapse_DN
P<0.001
Q<0.001
Primary tumors
TCGA TP53 biallelic inactivation vs P53-WT
TCGA TP53 biallelic vs TP53-WT
J
Relative ssGSEA score: TP53 biallelic vs TP53-WT
K
Relative ssGSEA score: TP53 biallelic vs TP53-WT

**Fig. S7: Clinical data analysis of *TP53* perturbation in lung cancer brain metastases (related to Fig. 1).**

**A-D.** *TP53* alterations in metastasis from primary lung tumors: **a,** Non-small cell lung cancer metastasis, *TP53* inactivation vs *TP53*-WT: BM vs primary tumors, Fisher's exact test $p<1*10^{-5}$, BM vs other metastasis, Fisher's exact test $p=8*10^{-4}$. *TP53* biallelic inactivation vs *TP53*-WT, BM vs primary tumors, Fisher's exact test, $p<1*10^{-5}$, BM vs other metastasis $p=<1*10^{-5}$. Primary n=915, brain n=59, chest wall n=11, liver n=68, lung n=10, soft tissue n=12, lymph node n=211, bone n=48, adrenal gland n=19, pleura n=73, pleural fluid n=25.

**B,** Lung adenocarcinoma metastasis, *TP53* inactivation vs *TP53*-WT: BM vs primary tumors, Fisher's exact test $p< 1*10^{-5}$, BM vs other metastasis, Fisher's exact test p=0.0017. *TP53* biallelic inactivation vs *TP53*-WT, BM vs primary tumor, Fisher's exact test, $p<1*10^{-5}$, BM vs other metastasis $p<1*10^{-4}$. Primary n=781, brain n=53**,** chest wall n=10, liver n=65, lung n=, lymph node n=152, bone n=40, soft tissue n=10, pleural fluid n=24, pleura n=68, adrenal gland n=16, other n=425.

**C,** Lung squamous cell carcinoma metastasis, *TP53* inactivation vs *TP53*-WT: BM vs primary tumors, Fisher's exact test p=0.13. BM vs other metastasis, Fisher's exact test p=0.55. *TP53* biallelic inactivation vs *TP53*-WT, BM vs primary tumor, Fisher's exact test, p=0.57. BM vs other metastasis p=0.01. Primary n=122, brain n=3, pleura n=4, bone n=7, lymph node n=14.

**D,** Poorly differentiated, non-small cell lung cancer, *TP53* inactivation vs *TP53*-WT: BM vs primary tumors, Fisher's exact test p=1, BM vs other metastasis, Fisher's exact test p=1. *TP53* biallelic inactivation vs *TP53*-WT, BM vs primary tumors, Fisher's exact test, p=0.4. BM vs other metastasis p=0.05. Primary n=12, brain n=3, bone n=1, lymph node n=5, pleura n=1, adrenal gland n=1, liver n=1.

**E,** Comparison of gene expression profiles from primary lung tumors with and without *TP53* alterations.

**F,** GSEA comparing gene signatures of BM between primary *TP53*-deficient and *TP53*-WT lung adenocarcinomas (source: TCGA, PanCancer Atlas) (*35*). *TP53*-deficient primary tumors express gene signatures associated with BM.

*TP53* inactivation vs *TP53*-WT, SMID_BREAST_CANCER_RELAPSE_IN_BRAIN_UP p=0.002, _DN p=0. *TP53* inactivation n=366, *TP53*-WT n=185.

**G,** Single-Sample GSEA confirms that *TP53* inactivation is associated with increased expression levels of brain-specific metastasis signatures. SMID_BREAST_CANCER_RELAPSE_IN_BRAIN $p=2.1*10^{-23}$, BRAIN _UP $p=1.5*10^{-11}$, BRAIN _DN $p=2.7*10^{-18}$.

**H,** Single-sample GSEA shows that *TP53* inactivation is associated with decreased expression levels of bone-specific metastasis signatures.

SMID_BREAST_CANCER_RELAPSE_IN_BONE $p=1.2*10^{-9}$, BONE _UP $p=4.8*10^{-8}$, BONE _DN $p=5.9*10^{-5.}$

**I,** GSEA comparing gene signatures of BM between primary *TP53*-null (biallelic inactivation) and TP53-WT lung adenocarcinomas (source: TCGA, PanCancer Atlas). SMID_BREAST_CANCER_RELAPSE_IN_BRAIN_UP p=0.01, BRAIN _DN $p<1*10^{-4}$. *TP53*-null n=169, *TP53*-WT n=185.

**J,** Single-Sample GSEA confirms that *TP53* biallelic inactivation is associated with increased expression levels of brain-specific metastasis signatures. SMID_BREAST_CANCER_RELAPSE_IN_BRAIN $p=2.3*10^{-27}$, BRAIN_UP $p=8.2*10^{-15}$, BRAIN_DN $p=1.1*10^{-19}$.

**K,** Single-sample GSEA shows that *TP53* inactivation is associated with decreased expression levels of bone-specific metastasis signatures. SMID_BREAST_CANCER_RELAPSE_IN_BONE $p=1.69*10^{-10}$, BONE_UP $p=1.75*10^{-8}$, BONE_DN $p=1.7*10^{-5}$.

Fig.S8

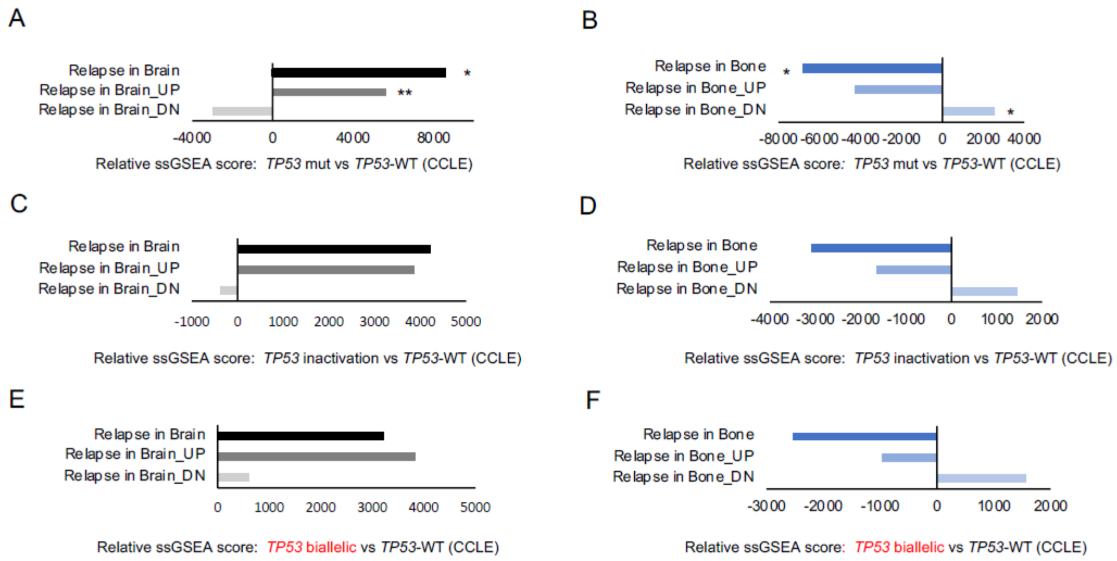

A
Relapse in Brain
Relapse in Brain_UP
Relapse in Brain_DN
*
**
-4000
0
4000
8000
Relative ssGSEA score: TP53 mut vs TP53-WT (CCLE)
B
Relapse in Bone
Relapse in Bone_UP
Relapse in Bone_DN
*
*
-8000 -6000 -4000 -2000 0 2000 4000
Relative ssGSEA score: TP53 mut vs TP53-WT (CCLE)
C
Relapse in Brain
Relapse in Brain_UP
Relapse in Brain_DN
-1000 0 1000 2000 3000 4000 5000
Relative ssGSEA score: TP53 inactivation vs TP53-WT (CCLE)
D
Relapse in Bone
Relapse in Bone_UP
Relapse in Bone_DN
-4000 -3000 -2000 -1000 0 1000 2000
Relative ssGSEA score: TP53 inactivation vs TP53-WT (CCLE)
E
Relapse in Brain
Relapse in Brain_UP
Relapse in Brain_DN
0 1000 2000 3000 4000 5000
Relative ssGSEA score: TP53 biallelic vs TP53-WT (CCLE)
F
Relapse in Bone
Relapse in Bone_UP
Relapse in Bone_DN
-3000 -2000 -1000 0 1000 2000
Relative ssGSEA score: TP53 biallelic vs TP53-WT (CCLE)

**Fig. S8: Analysis of gene expression profiles between CCLE BC cells lines with or without *TP53* inactivation (related to Fig. 2).**

**A-F** Single sample gene expression analysis (ssGSEA) comparing human BC cell lines that carry *TP53* inactivation with *TP53*-WT BC cell lines (n=43). **A,B,** Increased expression of signatures associated with BM and decreased expression of signatures associated with bone metastasis. *TP53* mut versus *TP53*-WT**,** one-sided student's t-test SMID_BREAST_CANCER_RELAPSE_IN_BRAIN p=0.02, BRAIN _UP p=0.01, BRAIN _DN p=0.11, SMID_BREAST_CANCER_RELAPSE_IN_BONE P=0.04, BONE _UP p= 0.07, BONE _DN p=0.02. **C-F** The same trend was observed when comparing *TP53* inactivation to *TP53*-WT (c,d,) and *TP53* biallelic inactivation to *TP53*-WT (e,f,), although it did not reach significance. **C,D,** *TP53* inactivation vs *TP53*-WT, SMID_BREAST_CANCER_RELAPSE_IN_BRAIN p=0.18, BRAIN _UP p=0.07, BRAIN _DN p=0.43. SMID_BREAST_CANCER_RELAPSE_IN_BONE p=0.22, BONE _UP p=0.28, BONE _DN p=0.16. **E,F,** *TP53* inactivation vs *TP53*-WT, SMID_BREAST_CANCER_RELAPSE_IN_BRAIN p=0.25, BRAIN _UP p=0.09, BRAIN _DN P=0.38. SMID_BREAST_CANCER_RELAPSE_IN_BONE p=0.25, BONE _UP p=0.36, BONE _DN p=0.15. BC cell lines: *TP53*-WT n=4, *TP53*-mut n=20, *TP53*-null n=16, *TP53* inactivated n=43.

Fig.S9

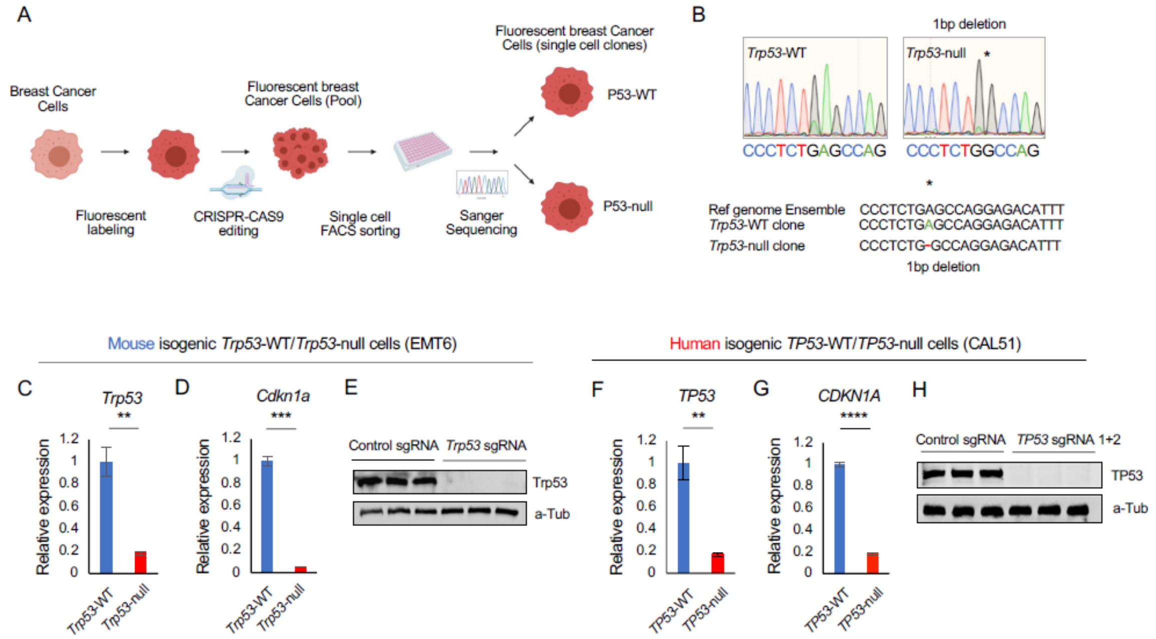

A
Breast Cancer Cells
Fluorescent labeling
CRISPR-CAS9 editing
Fluorescent breast Cancer Cells (Pool)
Single cell FACS sorting
Sanger Sequencing
Fluorescent breast Cancer Cells (single cell clones)
P53-WT
P53-null
B
1bp deletion
Trp53-WT
Trp53-null
CCCTCTGAGCCAG
CCCTCTGGCCAG
Ref genome Ensemble CCCTCTGAGCCAGGAGACATTT
Trp53-WT clone CCCTCTGAGCCAGGAGACATTT
Trp53-null clone CCCTCTG-GCCAGGAGACATTT
1bp deletion
Mouse isogenic Trp53-WT/Trp53-null cells (EMT6)
Human isogenic TP53-WT/TP53-null cells (CAL51)
C
Trp53
**
D
Cdkn1a
***
E
Control sgRNA
Trp53 sgRNA
Trp53
a-Tub
F
TP53
**
G
CDKN1A
****
H
Control sgRNA
TP53 sgRNA 1+2
TP53
a-Tub
Relative expression
1.2
1
0.8
0.6
0.4
0.2
0
Trp53-WT
Trp53-null
TP53-WT
TP53-null

**Fig. S9: Generation of mouse and human isogenic *Trp53*/*TP53*-null and *Trp53*-*TP53*-WT BC cells (related to Fig. 2).**

**A,** Workflow of isogenic cell line generation of *Trp53*-WT and *Trp53*-null BC cells. Murine EMT6 cells were fluorescently labeled, subjected to CRISPR-CAS9 editing for *Trp53* and subsequently sorted and grown as single cell clones. *Trp53* status was assessed by Sanger Sequencing. Human *TP53*-null CAL51 cells were generated in the same way, except without single-cell sorting, resulting in a heterogeneous population of *TP53*-null cells.

**B,** Sanger Sequencing confirmed *Trp53*-WT and *Trp53*-null status of EMT6 single cell clones. 5 independent single cell clones were mixed per genotype to obtain the *Trp53*-WT/*Trp53*-null isogenic system.

**C,** qPCR measurement of the relative *Trp53* expression showing a strong reduction of *Trp53* transcript levels in *Trp53*-null EMT6 cells. sgRNA *Trp53* vs sgRNA control, one-sided t-test p=0.003.

**D,** qPCR measurement of the relative expression of the p53 target p21 (*Cdkn1A*), showing that it is drastically decreased in EMT6 *Trp53*-null cells, one-tailed t-test, $p=2*10^{-4}$.

**E,** Western blot analysis of p53 showing abundant p53 protein levels in EMT6 *Trp53*-WT cells, but no p53 protein in the *Trp53*-null cells.

**F-H,** Validation of human CAL51 isogenic *TP53*-WT/-null system. **F,G,** Strong reduction of both *TP53* and *CDKN1A* gene expression in CAL51 BC cells treated with *TP53* sgRNA guides. sgRNA *TP53* vs CAL51 control, one-tailed Student's t-test *TP53* p=0.003, *CDKN1A* $p=2*10^{-6}$**.** **H,** Western blot analysis of p53 showing abundant p53 protein levels in CAL51 *TP53*-WT cells, but no p53 protein in the *TP53*-null cells.

Fig.S10

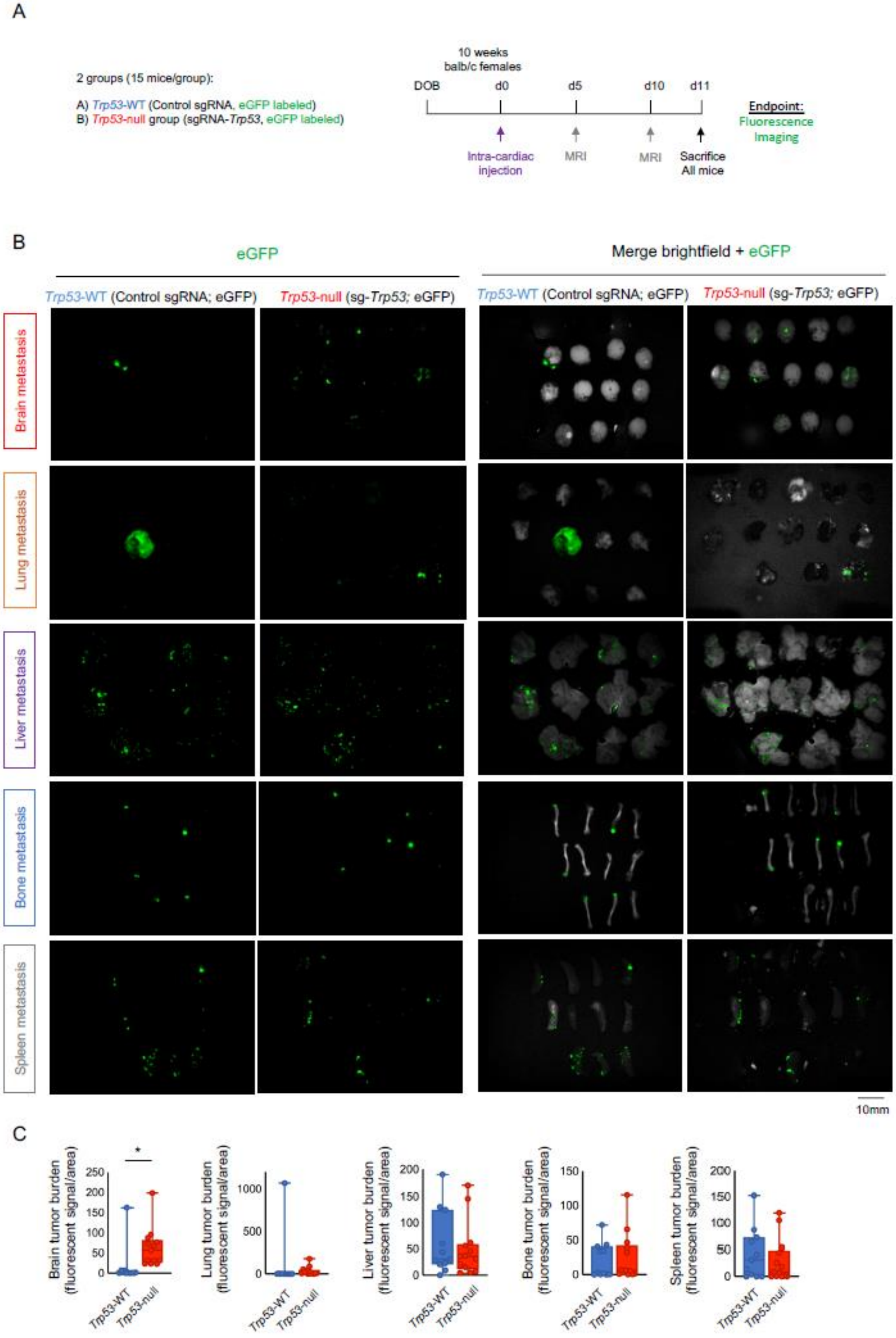
A
2 groups (15 mice/group):
A) Trp53-WT (Control sgRNA, eGFP labeled)
B) Trp53-null group (sgRNA-Trp53, eGFP labeled)
10 weeks
balb/c females
DOB
d0
d5
d10
d11
Intra-cardiac injection
MRI
MRI
Sacrifice All mice
Endpoint:
Fluorescence Imaging
B
eGFP
Merge brightfield + eGFP
Trp53-WT (Control sgRNA; eGFP)
Trp53-null (sg-Trp53; eGFP)
Trp53-WT (Control sgRNA; eGFP)
Trp53-null (sg-Trp53; eGFP)
Brain metastasis
Lung metastasis
Liver metastasis
Bone metastasis
Spleen metastasis
10mm
C
Brain tumor burden (fluorescent signal/area)
Lung tumor burden (fluorescent signal/area)
Liver tumor burden (fluorescent signal/area)
Bone tumor burden (fluorescent signal/area)
Spleen tumor burden (fluorescent signal/area)
*
Trp53-WT
Trp53-null

**Fig. S10: Organotropism of mouse *Trp53*-WT and *Trp53*-null EMT6 isogenic cells following intra-cardiac injection into separate immune-competent mice (related to Fig. 2).**

**A,** Experimental timeline of intra-cardiac injections of EMT6 *Trp53*-WT/ and *Trp53*-null isogenic cells. Two groups with 15 mice each were injected with either *Trp53*-WT or *Trp53*-null cells, both fluorescently labeled with eGFP. The entire cohort was sacrificed when mice started to die at day 11. At endpoint, organs were dissected (brain, lung, liver, bone, spleen) and eGFP fluorescence and brightfield images were taken.

**B,** Raw images showing eGFP fluorescence images (left) or a merge of eGFP fluorescence and brightfield images (right) in five organs dissected from mice that had been injected with *Trp53*-WT or *Trp53*-null isogenic cells. Shown are brain, lung, liver, bone and spleen. *Trp53*-WT n=11, *Trp53*-null n=13.

**C,** Additional quantification of tumor burden following intra-cardiac injection of mouse *Trp53*-WT or *Trp53*-null cells, supplementing Fig. 2H. Here, brain tumor burden was calculated as intensity of fluorescent signal relative to organ size. *Trp53*-null vs *Trp53*-WT injected mice, two-tailed Student's t-test for indicated organs: brain p=0.03, lung p=0.48, liver p=0.63, bone p=0.75, spleen p=0.49. *Trp53*-WT n=11, *Trp53*-null n=13.

Fig.S11

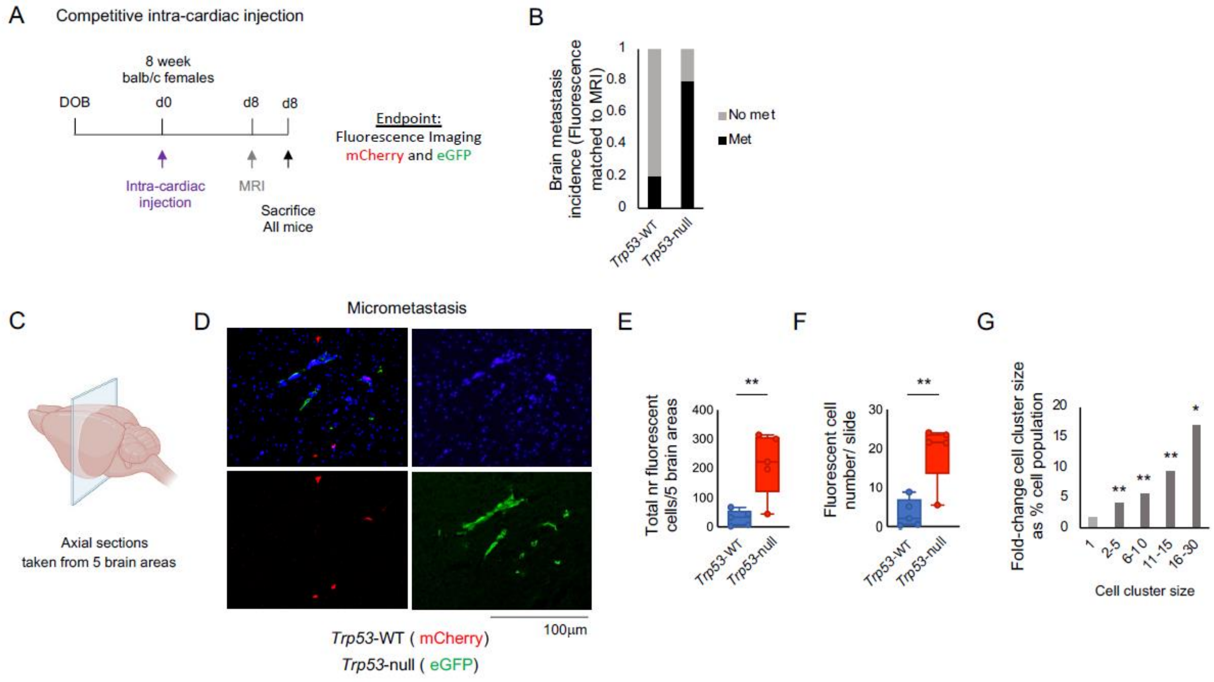

A
Competitive intra-cardiac injection
8 week
balb/c females
DOB
d0
d8
d8
Intra-cardiac injection
MRI
Sacrifice
All mice
Endpoint:
Fluorescence Imaging
mCherry and eGFP
B
Brain metastasis incidence (Fluorescence matched to MRI)
No met
Met
Trp53-WT
Trp53-null
C
Axial sections
taken from 5 brain areas
D
Micrometastasis
100µm
Trp53-WT ( mCherry)
Trp53-null ( eGFP)
E
Total nr fluorescent cells/5 brain areas
**
F
Fluorescent cell number/ slide
**
G
Fold-change cell cluster size as % cell population
Cell cluster size
1
2-5
6-10
11-15
16-30

**Fig. S11: Organotropism of mouse *Trp53*-WT and *Trp53*-null EMT6 isogenic cells following intra-cardiac injection into the same immune-competent mice (competition assay) (related to Fig. 2).**

**A,** Experimental timeline of intracardiac injections of EMT6 *Trp53*-WT/ and *Trp53*-null isogenic cells in a competition assay. Fluorescently-labeled *Trp53*-WT (red) and *Trp53*-null (green) EMT6 isogenic cells were injected into the same mice. The entire cohort was sacrificed after the first mice died. Brains were harvested, sectioned and subjected to immunofluorescence imaging. Mice n=5.

**B,** Fluorescence- and MRI-based quantification of BM incidence by isogenic *Trp53*-WT and *Trp53*-null cells following intra-cardiac injection into the same mice. Increased number of brain lesions were detected with *Trp53*-null eGFP-labeled cells in comparison to *Trp53*-WT (sgRNA control) mCherry-labeled cells. Fisher's exact test, brain p=0.20. n=5 mice.

**C,** Mouse brains were processed and tissue sections were obtained from 5 areas of each brain.

**D,** Immunofluorescent images of brain sections visualizing isogenic cancer cells. *Trp53*-null are labeled in green and *Trp53*-WT cells in red, sections were counterstained with DAPI to visualize cell nuclei.

**E-G,** Fluorescence-based quantification of micro metastasis derived from *Trp53*-WT and *Trp53*-null cells. **E,** The number of *Trp53*-null cells throughout the brain was significantly higher. Two-tailed Student's t-test p=0.006. **F,** The number of fluorescent *Trp53-null* cells per slide was significantly higher. Two-tailed Student's t-test p=0.003. **G,** Mouse brains contain bigger cell clusters of *Trp53*-null cells following competitive injection of labeled *Trp53*-null BC cells and their isogenic WT controls. Quantification of *Trp53*-null and *Trp53*-WT cell cluster size in brain sections from five brain areas. Fold-change *Trp53*-null over *Trp53*-WT as % cell population. Two-tailed Student's t-test Cluster size 2-5 cells p= 0.002, 6-10 cells p= 0.006, 11-15 cells p=0.003, 16-30 cells p=0.02. Mice n=5.

Fig.S12

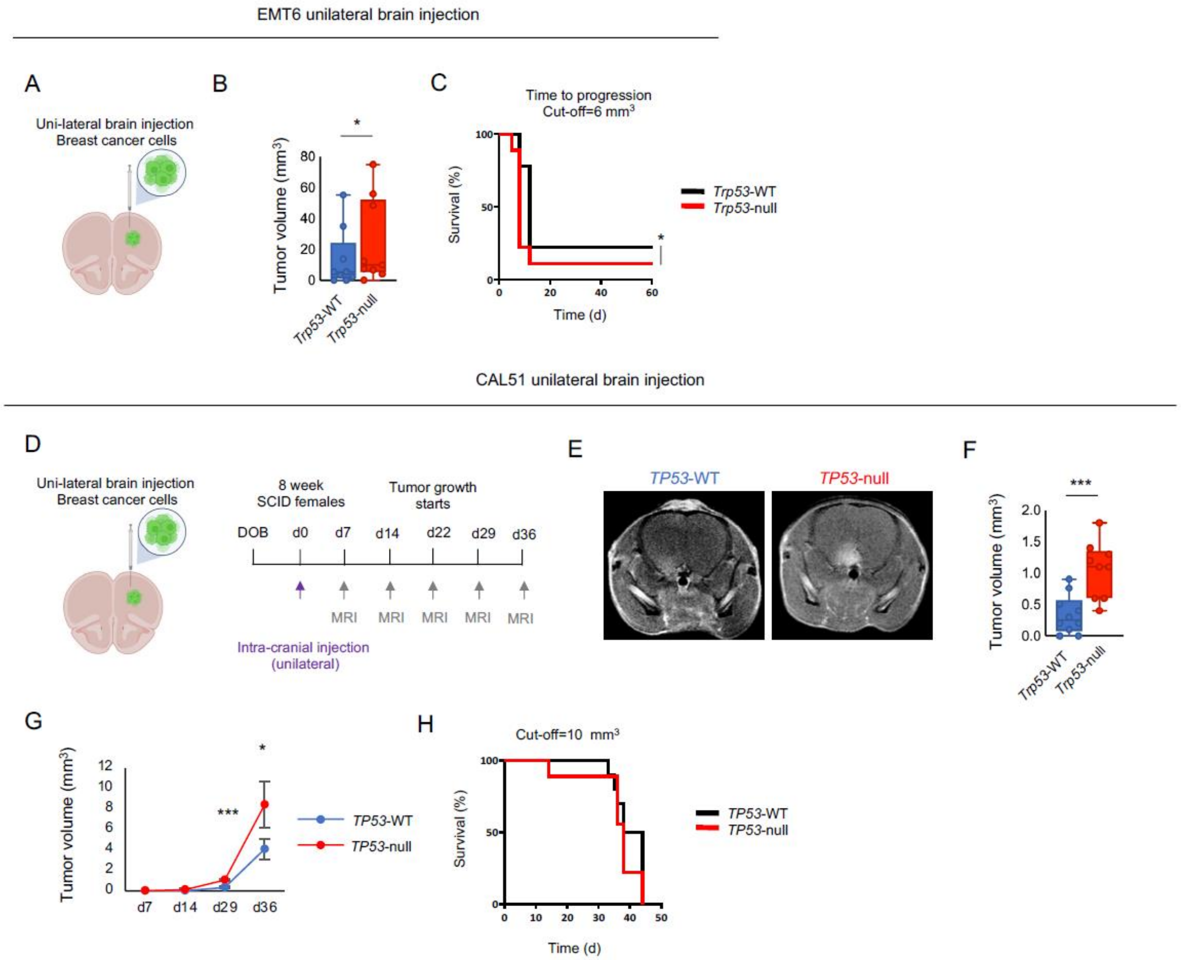
EMT6 unilateral brain injection
A
Uni-lateral brain injection
Breast cancer cells
B
Tumor volume (mm3)
Trp53-WT
Trp53-null
C
Time to progression
Cut-off=6 mm3
Survival (%)
Time (d)
Trp53-WT
Trp53-null
CAL51 unilateral brain injection
D
Uni-lateral brain injection
Breast cancer cells
8 week
SCID females
Tumor growth
starts
DOB
d0
d7
d14
d22
d29
d36
MRI
Intra-cranial injection
(unilateral)
E
TP53-WT
TP53-null
F
Tumor volume (mm3)
Trp53-WT
Trp53-null
G
Tumor volume (mm3)
TP53-WT
TP53-null
H
Cut-off=10 mm3
Survival (%)
Time (d)
TP53-WT
TP53-null

**Fig. S12: Unilateral intra-cranial injection of mouse and human *Trp53*-WT/*TP53*-WT and *Trp53*-null/*TP53*-null cells (related to Fig. 2).**

**A,** Unilateral intra-cranial injection of mouse isogenic *Trp53*-WT and *Trp53*-null cells into immune-competent mice. Mice were injected with either *Trp53*-WT or *Trp53*-null EMT6 isogenic BC cells directly into the brain (two mouse cohorts, n=10 per group).

**B,** MRI-based quantification of tumor volume 8 days after intra-cranial injection. Brain tumors that developed from injected EMT6 *Trp53*-null cells were larger than those from *Trp53*-WT cells. One-tailed Mann-Whitney test p=0.05. *Trp53*-WT n=9, *Trp53*-null n=9.

**C,** *Trp53*-null tumors reached a size threshold faster than their isogenic controls, time to progression with a cut-off of $6mm^3$, two-tailed Gehan-Breslow-Wilcoxon test p=0.035, *Trp53*-WT n=9 , *Trp53*-null n=9.

**D** Unilateral intra-cranial injection of human isogenic *TP53*-WT and *TP*53-null cells into immune-deficient (SCID) mice. Mice were injected with either *TP53*-WT or *TP53*-null CAL51 isogenic BC cells directly into the brain (two mouse cohorts, n=10 per group). The experimental timeline is shown, brains were imaged by MRI throughout a five-week period.

**E,** Representative MRI images of mouse brains 36 days after unilateral intra-cranial injection of the isogenic cells

**F,G,** MRI-based quantification of tumor volume 29 days after intra-cranial injection (**G**), also shown as a growth curve over time (**G**). One-sided Student's t-test d29 $p=4*10^{-5}$, d36 p=0.05. *TP53*-WT n=10, *TP53*-null n=9.

**H,** *TP53*-null tumors reached a size threshold faster than their isogenic controls, time to progression with a cut-off of $10mm^3$. Gehan-Breslow-Wilcoxon test p=0.37, *TP53*-WT n=10, *TP53*-null n=9.

Fig.S13

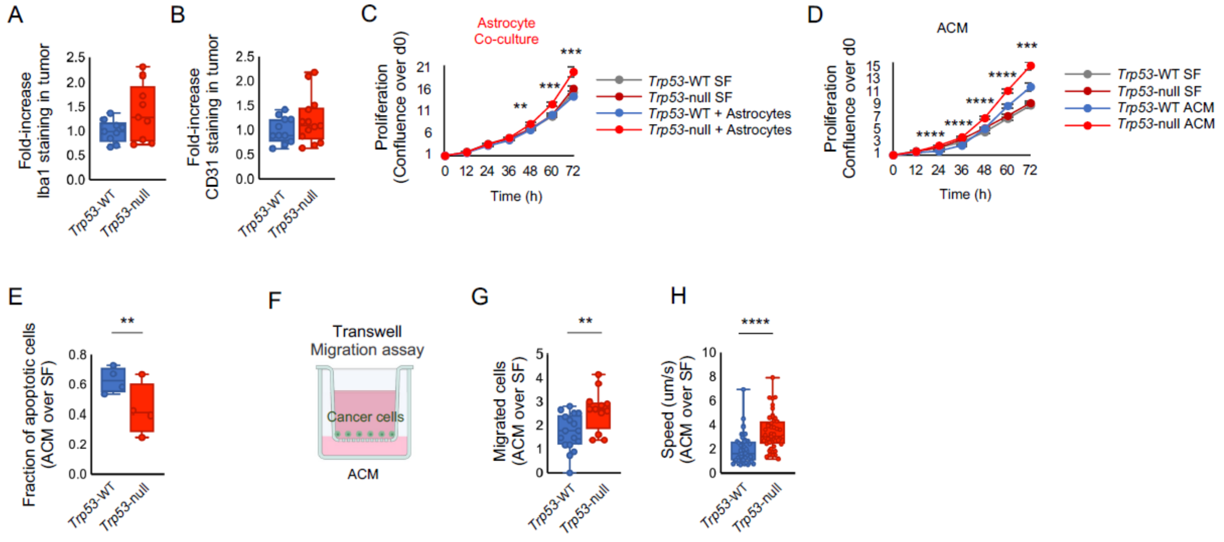
A
Fold-increase Iba1 staining in tumor
Trp53-WT
Trp53-null
B
Fold-increase CD31 staining in tumor
Trp53-WT
Trp53-null
C
Astrocyte Co-culture
Proliferation (Confluence over d0)
Time (h)
Trp53-WT SF
Trp53-null SF
Trp53-WT + Astrocytes
Trp53-null + Astrocytes
D
ACM
Proliferation Confluence over d0
Time (h)
Trp53-WT SF
Trp53-null SF
Trp53-WT ACM
Trp53-null ACM
E
Fraction of apoptotic cells (ACM over SF)
Trp53-WT
Trp53-null
F
Transwell Migration assay
Cancer cells
ACM
G
Migrated cells (ACM over SF)
Trp53-WT
Trp53-null
H
Speed (um/s) (ACM over SF)
Trp53-WT
Trp53-null

**Fig. S13: Mouse astrocytes infiltrate p53-null brain metastases and promote the survival, proliferation and migration of p53-null mouse BC cells in the brain (related to Fig. 3).**

**A,** Fluorescence-based quantification of microglia in the EMT6-derived tumors. Iba1 staining intensity was similar in tumors growing from *Trp53*-null and *Trp53*-WT BC cells. Data are shown as fold-increase in staining intensity over *Trp53*-WT, two-sided Student's t-test p=0.08, Tumor sections evaluated *Trp53*-WT n=9, Trp53-null n=9, mice n=3.

**B,** Fluorescence-based quantification of endothelial cells in the EMT6-derived tumors. CD31 staining intensity was similar in tumors growing from *Trp53*-null and *Trp53*-WT BC cells. Data are shown as fold-increase in staining intensity over *Trp53*-WT, two-tailed Student's t-test p=0.23, Tumor sections evaluated *Trp53*-WT n=11, *Trp53*-null n=13, mice n=3.

**C,** Live imaging-based proliferation curves of BC EMT6 cells co-cultured with mouse astrocytes. The effect of co-culture on the growth of *Trp53*-null BC cells is stronger than that of *Trp53*-WT cells. Two-sided t-test, *Trp53*-null vs *Trp53*-WT,
48h p=0.002, 60h p= p=$7*10^{-4}$, 72h p= p=$2*10^{-5}$. *Trp53*-WT n=8, Trp53-null n=8.

**D,** Live imaging-based proliferation curves of BC EMT6 cells cultured with ACM. The effect of ACM on the growth of *Trp53*-null BC cells is stronger than that of *Trp53*-WT cells. Two-sided t-test, *Trp53*-null vs *Trp53*-WT, 24h p=$1.2*10^{-10}$, 36h p=$1*10^{-10}$, 48h p=$4.5*10^{-7}$, 60h p=$1*10^{-5}$, 72h p=$2*10^{-4}$. *Trp53*-WT n=7, Trp53-null n=7.

**E,** Flow cytometry analysis of BC cells cultured with ACM. The fraction of cells in apoptosis is lower in *Trp53*-null than in *Trp53*-WT cells. AnnexinV-positive cells as fold-change relative to serum-free controls. Two-sided student's t-test, *Trp53*-null vs *Trp53*-WT, p=0.03. n=4.

**F,** Transwell migration assay to assess the migration of BC cells towards ACM. BC cells were seeded in SFM in the upper compartment and ACM was placed in the lower compartment.

**G,** Quantification of cell migration in the transwell assay after 24h of co-culture of BC cells and ACM. The effect of ACM on cell migration was significantly stronger in *Trp53*-null cells. Quantification of migrated cells is shown as the fold-increase of migrating cells in ACM over SFM, two-tailed Student's t-test p=0.005. Fields counted *TP53*-WT n=17, *TP53*-null n=12.

**H,** Live-imaging- based quantification of the effect of ACM on cell velocity of *Trp53*-WT and *Trp53*-null cells. Cell velocity shown as fold-increase in ACM over SF, two-sided Student's t-test p=$1*10^{-6}$. Each data point represents a velocity measurement.

Fig.S14

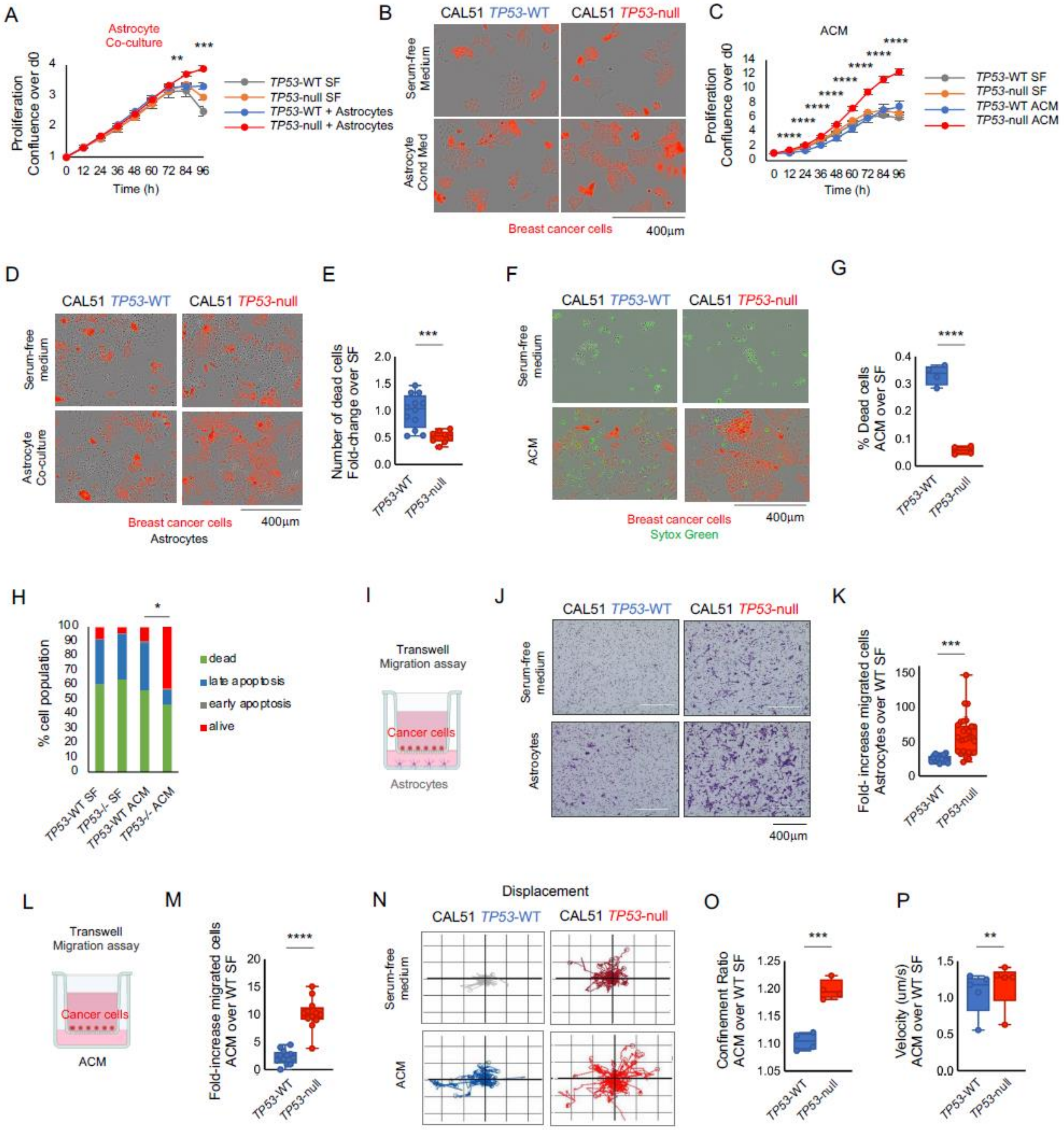

A
Astrocyte Co-culture
Proliferation Confluence over d0
Time (h)
TP53-WT SF
TP53-null SF
TP53-WT + Astrocytes
TP53-null + Astrocytes
B
CAL51 TP53-WT
CAL51 TP53-null
Serum-free Medium
Astrocyte Cond Med
Breast cancer cells
400µm
C
ACM
Proliferation Confluence over d0
Time (h)
TP53-WT SF
TP53-null SF
TP53-WT ACM
TP53-null ACM
D
CAL51 TP53-WT
CAL51 TP53-null
Serum-free medium
Astrocyte Co-culture
Breast cancer cells
Astrocytes
400µm
E
Number of dead cells Fold-change over SF
TP53-WT
TP53-null
F
CAL51 TP53-WT
CAL51 TP53-null
Serum-free medium
ACM
Breast cancer cells
Sytox Green
400µm
G
% Dead cells ACM over SF
TP53-WT
TP53-null
H
% cell population
dead
late apoptosis
early apoptosis
alive
TP53-WT SF
TP53-/- SF
TP53-WT ACM
TP53-/- ACM
I
Transwell Migration assay
Cancer cells
Astrocytes
J
CAL51 TP53-WT
CAL51 TP53-null
Serum-free medium
Astrocytes
400µm
K
Fold- increase migrated cells Astrocytes over WT SF
TP53-WT
TP53-null
L
Transwell Migration assay
Cancer cells
ACM
M
Fold-increase migrated cells ACM over WT SF
TP53-WT
TP53-null
N
Displacement
CAL51 TP53-WT
CAL51 TP53-null
Serum-free medium
ACM
O
Confinement Ratio ACM over WT SF
TP53-WT
TP53-null
P
Velocity (um/s) ACM over WT SF
TP53-WT
TP53-null

**Fig. S14: Human astrocytes infiltrate p53-null brain metastases and promote the survival, proliferation and migration of p53-null human BC cells in the brain (related to Fig. 3)**

**A,** Live imaging-based proliferation curves of human BC CAL51 cells co-cultured with activated human astrocytes. The effect of the astrocytes is stronger for the *TP53*-null cells in comparison to the *TP53*-WT cells. One-sided t-test at indicated time points: 84h, p=0.002; 96h, p=$1*10^{-4}$. *TP53*-WT n=8, *TP53*-null n=8.

**B,** Representative images of *TP53*-WT and *TP53*-null CAL51 cells grown for 48h in SFM or in ACM. *TP53*-null cells cultured in ACM had a significant growth advantage over their isogenic *TP53*-WT and serum-free controls.

**C,** Live imaging-based proliferation curves of human BC CAL51 cells cultured in ACM or in SFM. One-tailed t-test at indicated time points: 12h, p=$1*10^{-7}$; 24h p= $5*10^{-7}$; 36h p= $3*10^{-5}$; 48h p= $5*10^{-6}$; 60h p=$4*10^{-6}$; 72h p= $5*10^{-6}$; 84h p= $5*10^{-6}$; 96h p= $3*10^{-5}$. *TP53*-WT n=20, *TP53*-null n=20.

**D,** Representative images of *TP53*-WT and *TP53*-null CAL51 cells grown for 48h in SFM or co-cultured with activated human astrocytes. *TP53*-null cells cultured in ACM exhibited significantly reduced cell death in comparison to their isogenic *TP53*-WT and serum-free controls.

**E,** Morphology-based quantification of cell death in human BC CAL51 cells co-cultured with activated human astrocytes or cultured in SFM (at 72h). *TP53*-null cells exhibit significantly reduced cell death in comparison to *TP53*-WT cells. One-sided Student's t-test P=$4*10^{-4}$. Fields counted, *TP53*-WT n=10, *TP53*-null n=7.

**F,** Representative images of *TP53*-WT and *TP53*-null CAL51 cells grown for 5 days in SFM or in ACM. *TP53*-null cells cultured in ACM exhibited significantly reduced cell death in comparison to their isogenic *TP53*-WT and serum-free controls, as measured by Sytox green staining of dying cells (green).

**G,** Fluorescence-based quantification of cell death co-cultured with activated human astrocytes or cultured in SFM (at 120h). ACM rescued cell death to a greater extent in *T53*-null cells. Shown are dying cells (green) and living cells (red) as percentage of cell population. p=$1*10^{-5}$. *TP53*-WT n=4, *TP53*-null n=4.

**H,** Flow cytometry analysis of apoptosis in CAL51 *TP53*-WT and *TP53*-null cells cultured in ACM for 3 days. Cells were labeled with AnnexinV and with Sytox-green. Quantification of cells that are alive (red), in early apoptosis (grey), late apoptosis (blue) and dead (green). *TP53*-null vs *TP53*-WT cells, dead vs alive, Chi-square test, p=0.03. *TP53*-WT n=3, *TP53-null* n=3.

**I,** Transwell migration assay to assess the migration of human BC cells towards human activated astrocytes.

**J,** Representative images of human *TP53*-WT and *TP53*-null BC CAL51 cells, stained in purple, showing their migration through the filter towards the lower chamber, which contains either SFM or human astrocytes.

**K,** Quantification of cell migration in the transwell assay. *TP53*-null cells exhibit increased migratory potential already under serum-free conditions, and their migratory capacity is considerably increased in the presence of astrocytes. *TP53*-null vs *TP53*-WT grown with astrocytes, $P=2*10^{-5}$. Fields counted *TP53*-WT n=26, *TP53*-null- n=28.

**L,** Transwell migration assay to assess the migration of human BC cells towards human ACM.

**M,** Quantification of cell migration in the transwell assay. *TP53*-null cells exhibit augmented migratory potential upon exposure to ACM. Shown as fold-increase of migratory cells in ACM over SFM. One-sided Student's t-test $5*10^{-8}$. Fields counted *TP53*-WT n=16, *TP53*-null n=28.

**N,** Representative live cell imaging-based tracking of cell movement of individual human BC cells. Displacement plots show that ACM stimulated cell movement both in *TP53*-WT and in *TP53*-null cells, with a larger effect on *TP53*-null cells.

**O,** Live-imaging- based quantification of the effect of ACM on the confinement ratio (degree of cell motility) of *TP53*-WT and *TP53*-null cells. Confinement ratio shown as fold-increase in ACM over WT SF, one-sided Student's t-test p=0.0005, n=4.

**P,** Live-imaging- based quantification of the effect of ACM on cell velocity of *TP53*-WT and *TP53*-null cells. Cell velocity shown as fold-increase in ACM over WT SF, one-sided Student's t-test p=0.004, *TP53*-WT n=5, *TP53*-null n=5.

Fig.S15

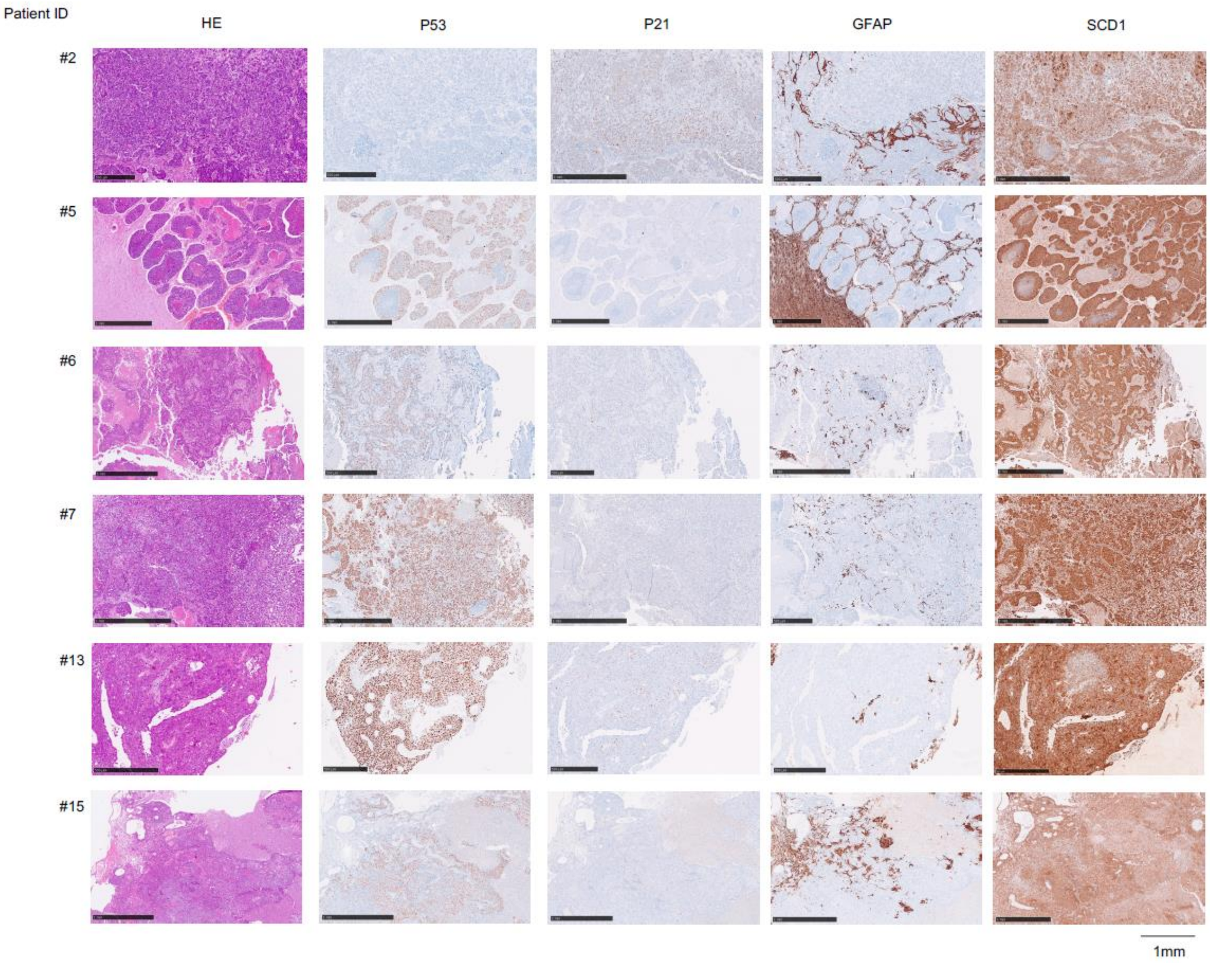

**Fig. S15: Human BCBM exhibit low levels of p53 signaling, astrocyte tumor infiltration, and high levels of SCD1 expression.**

**A,** Representative images of BCBM sections from 6 patients out of an analyzed cohort of 17 patient samples (Source: Luca Bertero), detailed in Table S5. Sections were stained for haematoxylin-eosin (H&E) or with antibodies labeling p53, p21 GFAP or SCD1 protein. All sections showed very low or no detectable p21, staining indicating a strong reduction or absence of p53 activity. Astrocyte infiltration could be detected in many tumors but the degree of infiltration was highly variable. Strong SCD1 staining was detected across samples, consistent with upregulation of SCD1 level in BCBM. For protein quantification in all samples, see Table S5.

Fig.S16

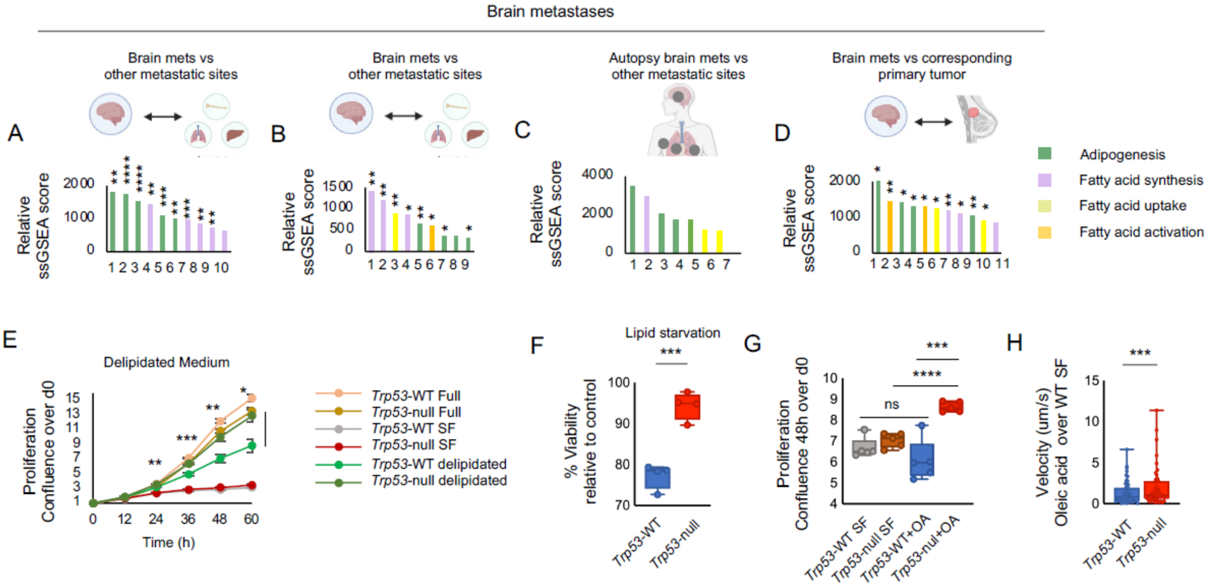

Brain metastases
A
Brain mets vs
other metastatic sites
Relative
ssGSEA score
B
Brain mets vs
other metastatic sites
C
Autopsy brain mets vs
other metastatic sites
D
Brain mets vs corresponding
primary tumor
Adipogenesis
Fatty acid synthesis
Fatty acid uptake
Fatty acid activation
E
Delipidated Medium
Proliferation
Confluence over d0
Time (h)
Trp53-WT Full
Trp53-null Full
Trp53-WT SF
Trp53-null SF
Trp53-WT delipidated
Trp53-null delipidated
F
Lipid starvation
% Viability
relative to control
Trp53-WT
Trp53-null
G
Proliferation
Confluence 48h over d0
ns
Trp53-WT SF
Trp53-null SF
Trp53-WT+OA
Trp53-nul+OA
H
Velocity (um/s)
Oleic acid over WT SF
Trp53-WT
Trp53-null

**Fig. S16: p53-dependent fatty acid metabolism in BC cells (related to Fig. 4).**

**A-D,** Single-sample gene set enrichment analysis (ssGSEA) shows elevated gene expression signatures related to adipogenesis, fatty acid (FA) synthesis, FA uptake and FA activation in: **A,B,** brain metastases compared to metastases in other organs (GSE14018, Giovanni Blandino) Table S7; **C,** BM compared to metastases in other organs, using transcriptomic data from samples taken at autopsy from BROCADE patient 6071; **D,** brain metastases compared to their matched primary breast tumors (data from GSE125989 (*61*)). Table S8.

**E,** Live imaging-based proliferation curves of *Trp53*-WT and *Trp53*-null mouse BC cells cultured in full medium, SFM or delipidated full medium. *Trp53*-null EMT6 cells grow better than their isogenic *Trp53*-WT controls when lipid levels are limited, in line with increased fatty acid synthesis in *Trp53*-null cells. *Trp53*-null vs *Trp53*-WT cells under delipidated conditions, at indicated time points: 24h p=0.005, 36h p=0.001, 48h p=0.009, 60h p=0.02. Two-tailed Student's t-test. *Trp53*-WT n=4, *Trp53*-null n=4.

**F,** Comparison of the effect of lipid starvation on the growth of *Trp53*-WT and *Trp53*-null BC cells. *Trp53*-null cells survive better under lipid starvation conditions, two-sided Student's t-test $p=3*10^{-4}$, *Trp53*-WT n=4, *Trp53*-null n=4.

**G,** Live imaging-based proliferation of *Trp53*-WT and *Trp53*-null mouse BC cells cultured in SFM with or without OA (10µM) for 48h. OA promotes the growth of *Trp53*-null cells more than that of *Trp53*-WT cells. Two-sided Student's t-test *Trp53*-WT vs. *Trp53*-WT cells treated with OA p=0.26; *Trp53*-null vs. *Trp53*-null cells treated with OA $p=3*10^{-5}$, OA treated *Trp53*-null vs *Trp53*-WT cells relative to SF controls p= 0.001, *Trp53*-WT n=4, *Trp53*-null n=4.

**H,** Live-imaging- based quantification of the effect of OA on cell migration (velocity) of *Trp53*-WT and *Trp53*-null cells. Cell velocity shown as fold-increase in OA over SF. The effect of OA on cell velocity was significantly stronger in *Trp53*-null cells. *Trp53*-null vs *Trp53*-WT, two-sided Student's t-test $p= 3*10^{-91}$.

### Fig.S17

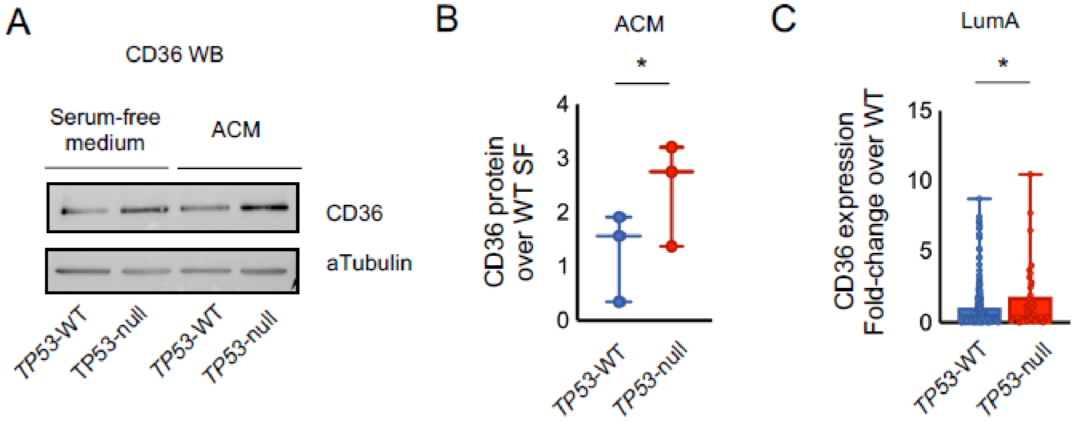
A
CD36 WB
Serum-free medium
ACM
CD36
aTubulin
TP53-WT
TP53-null
TP53-WT
TP53-null
B
ACM
*
CD36 protein over WT SF
4
3
2
1
0
TP53-WT
TP53-null
C
LumA
*
CD36 expression Fold-change over WT
15
10
5
0
TP53-WT
TP53-null

**Fig. S17: The fatty acid transporter CD36 is elevated in *TP53*-null human BC cells (related to Fig. 4).**

**A,** Western blot CD36 protein quantification in isogenic *TP53*-WT and *TP53*-null CAL51 cells cultured in SFM or in ACM. CD36 protein levels were significantly higher in ACM-exposed *TP53*-null cells.

**B,** Quantification of CD36 protein levels from three separate western blot experiments. The effect of ACM on CD36 protein expression was significantly stronger in *TP53*-null CAL51 cells in comparison to *TP53*-WT cells. CD36 protein levels are shown as fold-change of ACM over WT SF conditions. One-sided Student's t-test, p=0.02, n=3.

**C,** Comparison of *CD36* mRNA expression levels between *TP53*-WT and *TP53*-null human primary breast carcinomas of the Luminal A molecular subtype, based on TCGA data. *TP53*-null tumors exhibited higher *CD36* mRNA expression levels in comparison to *TP53*-WT tumors. Shown as fold-increase *CD36* expression relative to the average expression of *TP53*-WT tumors. One-sided Student's t-test p=0.05, *TP53*-WT n=206, *TP53*-null n=41.

Fig.S18

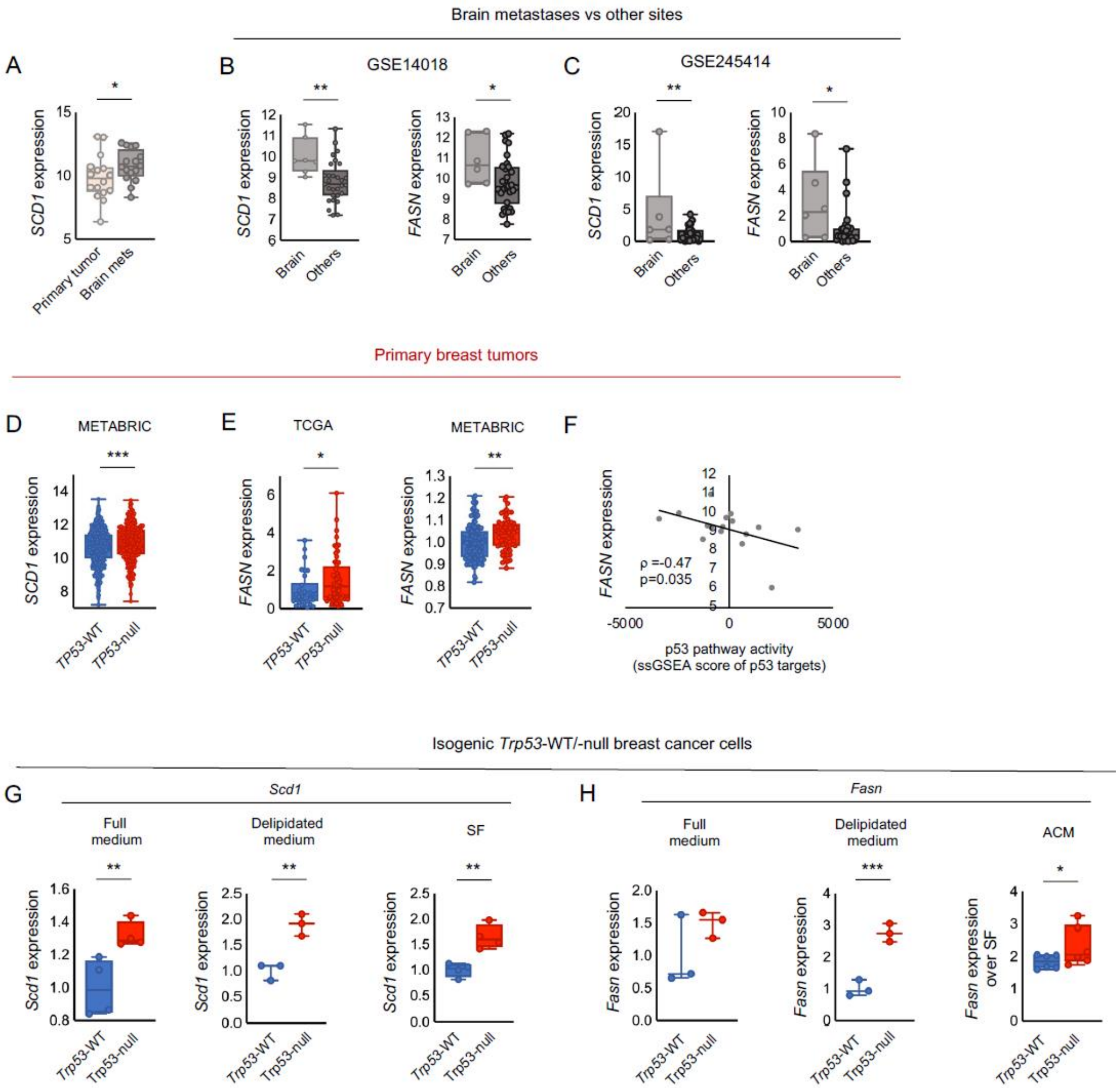
Brain metastases vs other sites
A
B
GSE14018
C
GSE245414
SCD1 expression
FASN expression
Primary tumor
Brain mets
Brain
Others
Primary breast tumors
D
METABRIC
E
TCGA
METABRIC
F
TP53-WT
TP53-null
ρ =-0.47
p=0.035
p53 pathway activity
(ssGSEA score of p53 targets)
Isogenic Trp53-WT/-null breast cancer cells
G
Scd1
H
Fasn
Full medium
Delipidated medium
SF
ACM
Scd1 expression
Fasn expression
Fasn expression over SF
Trp53-WT
Trp53-null

**Fig. S18: Fatty acid synthesis is upregulated in p53-null BC cells (related to Fig. 5).**

**A,** Comparison of *SCD1* mRNA expression levels between primary tumors to their matched brain metastases (GSE125989). SCD1 expression is significantly higher in the BM in comparison to their primary tumors of origin. Paired two-sided Student's t-test p=0.05. n=16 matched primary tumors and brain metastases.

**B,** Comparison of *SCD1*and *FASN* mRNA expression levels between BM and metastases in other organs (GSE1418), showing elevated SCD1 expression BM. Brain vs others, two-sided Student's t-test *SCD1* p=0.006, *FASN* p=0.05. BM n=7, metastases in other sites n=29.

**C,** Comparison of the mRNA expression levels of the fatty acid synthesis enzymes *SCD1*, and *FASN* between BM and metastases in other organs (GSE245414). The expression of both enzymes is significantly increased in BM in comparison to others metastatic sites. One-sided Student's t-test, *SCD1* p=0.01, *FASN* p=0.02. BM n=6, metastases in other metastatic sites n=26.

**D,** Comparison of *SCD1* mRNA expression levels between *TP53*-WT and *TP53*-null human primary breast tumors, based on METABRIC data. One-sided Student's t-test $p=5*10^{-4}$. *TP53*-WT n=592, *TP53*-null n=320.

**E,** Comparison of *FASN* mRNA expression levels between *TP53*-WT and *TP53*-null human primary breast tumors of the Luminal B molecular subtype, based on TCGA and METABRIC data. *TP53*-null tumors exhibited higher *FASN* mRNA expression levels in comparison to *TP53*-WT tumors. Shown as fold-increase *FASN* expression relative to the average expression of *TP53*-WT tumors. One-sided Student's t-test TCGA p=0.04, METABRIC p= 0.002. TCGA *TP53*-WT n=30, *TP53*-null n=56; METABRIC *TP53*-WT n=128, *TP53*-null n=63.

**F,** The correlation between the p53 pathway activity score and the mRNA expression levels of FASN across primary human breast tumors that metastasized to the brain (GSE125989). Low p53 pathway activity is associated with high FASN mRNA expression levels. p53 pathway activity was evaluated by calculating an ssGSEA score for the MSigDBgene signature 'KANNAN_p53_targets'. Spearman's rho=-0.47, p=0.035, n=16.

**G,** RT-qPCR-based analysis of *Scd1* mRNA expression in *Trp53*-WT and *Trp53*-null mouse BC EMT6 cells cultured in full medium (left), delipidated medium (center) or SFM (right). *Scd1* expression was significantly higher in *Trp53*-null cells under all tested conditions. One-sided Student's t-test, full medium p=0.01, delipidated medium p=0.003, and SFM p=0.003. Full medium n=4, delipidated medium n=3, SFM n=4.

**H,** qPCR-based analysis of *Fasn* mRNA expression in *Trp53*-WT and *Trp53*-null mouse BC EMT6 cells cultured in full medium (left), delipidated medium (center) or ACM (right). *Fasn* expression was significantly higher in *Trp53*-null cells under all tested conditions. One-tailed Student's t-test, full medium p=0.11, delipidated medium $p=5*10^{-4}$, and ACM p=0.05; full medium n=3, delipidated medium n=3, ACM n=6 and n=3 for Trp53-WT and Trp53-null, respectively.

Fig.S19

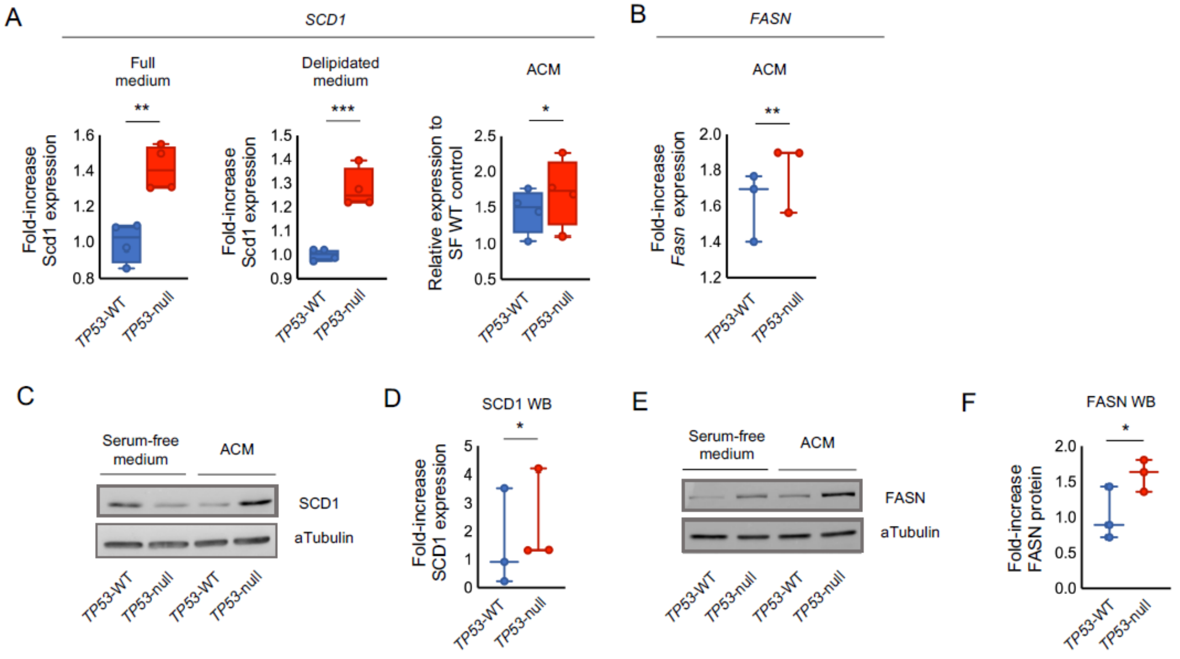
A
SCD1
Full medium
Delipidated medium
ACM
Fold-increase Scd1 expression
Relative expression to SF WT control
TP53-WT
TP53-null
B
FASN
ACM
Fold-increase Fasn expression
C
Serum-free medium
ACM
SCD1
aTubulin
D
SCD1 WB
Fold-increase SCD1 expression
E
FASN
F
FASN WB
Fold-increase FASN protein

**Fig. S19: Fatty acid synthesis is upregulated in *TP53*-null human BC cells, and is further stimulated by astrocyte conditioned medium (related to Fig. 5).**

**A,** qPCR-based analysis of *SCD1* mRNA expression in *TP53*-WT and *TP53*-null human BC CAL51 cells cultured in full medium (left, one-tailed Student's t-test p=0.003), delipidated medium (center, one-tailed $p=6*10^{-4}$) or ACM (right, one-tailed Student's t-test p=0.04). *SCD1* expression was significantly higher in *TP53*-null cells under all tested conditions. Full medium n=4, delipidated medium n=4, ACM n=3.

**B,** qPCR-based analysis of *FASN* mRNA expression in *TP53*-WT and *TP53*-null human BC CAL51 cells cultured in ACM. *FASN* expression was significantly higher in *TP53*-null cells. One-sided Student's t-test, p=0.008, n=3.

**C,** Western blot SCD1 protein quantification in isogenic *TP53*-WT and *TP53*-null CAL51 cells cultured in SFM or in ACM. ACM exposure significantly elevated SCD1 protein levels in *TP53*-null cells.

**D**, Quantification of SCD1 protein levels from three separate western blot experiments. The effect of ACM on SCD1 protein expression was significantly stronger in *TP53*-null cells in comparison to *TP53*-WT cells. SCD1 protein levels are shown as fold-change of ACM over SF conditions. One-sided Student's t-test p=0.03, *TP53*-WT n=3, *TP53*-null n=3.

**E,** Western blot FASN protein quantification in isogenic *TP53*-WT and *TP53*-null CAL51 cells cultured in SFM or in ACM. ACM exposure significantly elevated FASN protein levels in *TP53*-null cells.

**F**, Quantification of FASN protein levels from three separate western blot experiments. The effect of ACM on FASN protein expression was significantly stronger in *TP53*-null cells in comparison to *TP53*-WT cells. FASN protein levels are shown as fold-change of ACM over SF conditions. One-sided Student's t-test p=0.02, *TP53*-WT n=3, *TP53*-null n=3.

Fig.S20

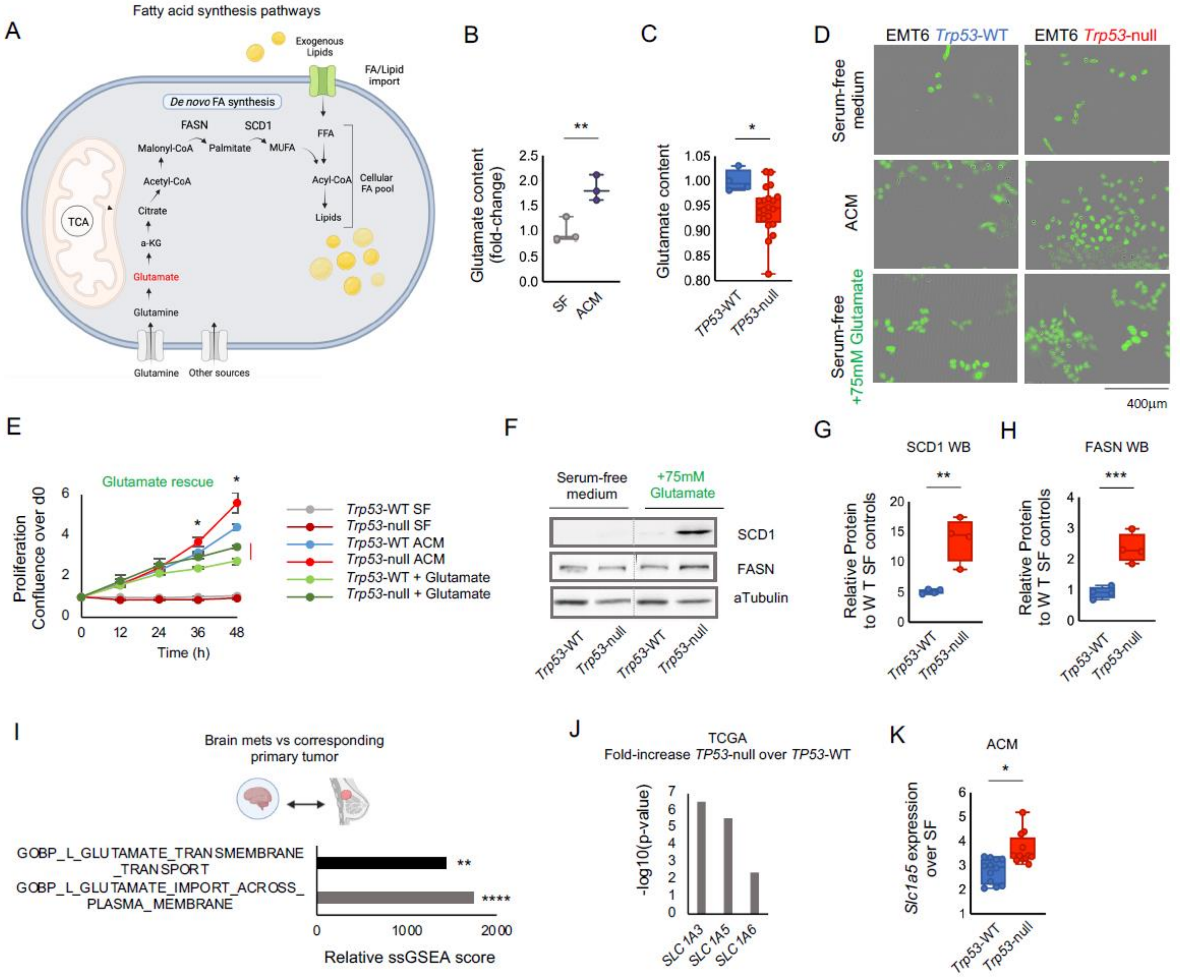

A
Fatty acid synthesis pathways
Exogenous Lipids
FA/Lipid import
De novo FA synthesis
FASN
SCD1
Malonyl-CoA
Palmitate
MUFA
FFA
Acetyl-CoA
Acyl-CoA
Cellular FA pool
Citrate
Lipids
TCA
a-KG
Glutamate
Glutamine
Glutamine
Other sources
B
Glutamate content (fold-change)
SF
ACM
**
C
Glutamate content
TP53-WT
TP53-null
*
D
EMT6 Trp53-WT
EMT6 Trp53-null
Serum-free medium
ACM
Serum-free +75mM Glutamate
400µm
E
Glutamate rescue
Proliferation Confluence over d0
Time (h)
Trp53-WT SF
Trp53-null SF
Trp53-WT ACM
Trp53-null ACM
Trp53-WT + Glutamate
Trp53-null + Glutamate
F
Serum-free medium
+75mM Glutamate
SCD1
FASN
aTubulin
Trp53-WT
Trp53-null
Trp53-WT
Trp53-null
G
SCD1 WB
Relative Protein to W T SF controls
**
H
FASN WB
Relative Protein to W T SF controls
***
I
Brain mets vs corresponding primary tumor
GOBP_L_GLUTAMATE_TRANSMEMBRANE_TRANSPORT
GOBP_L_GLUTAMATE_IMPORT_ACROSS_PLASMA_MEMBRANE
Relative ssGSEA score
J
TCGA
Fold-increase TP53-null over TP53-WT
-log10(p-value)
SLC1A3
SLC1A5
SLC1A6
K
ACM
Slc1a5 expression over SF
Trp53-WT
Trp53-null

**Fig. S20: Glutamate is secreted from astrocytes and stimulates SCD1 expression, FAS and cell proliferation (related to Fig. 5).**

**A,** Schematic illustration of FAS pathways and the metabolites that are utilized as fatty acid building blocks, such as glutamate. Image adapted from Tumanov et al. 2015 (*145*).

**B,** Liquid Chromatography-Mass spectrometry (LC-MS) analysis of polar metabolites reveals higher glutamate levels in ACM. Shown as fold-increase of glutamate abundance in ACM over SFM, two-sided Student's t-test p=0.01. SFM n=3, ACM n=3. Table S12.

**C,** Analysis of metabolic profiling data (*65*) across human BC cell lines reveals lower intra-cellular glutamate levels in *TP53*-null vs. *TP53*-WT BC cells. Glutamate content is shown as fold-change over *TP53*-WT. Two-sided Student's t-test p=0.03. *TP53*-WT n=4 *TP53*-null n=20.

**D,** Representative images of GFP-labeled *Trp53*-WT and *Trp53*-null EMT6 cells cultured in SFM (top), in ACM (middle) or in SFM supplemented with glutamate (75mM) for 48h. Glutamate supplementation improved the proliferation of the cells in SFM, and the effect was stronger in *Trp53*-null cells.

**E,** Live imaging-based proliferation curves of *Trp53*-WT and *Trp53*-null mouse BC cells cultured in SFM, ACM or SFM supplemented with glutamate (75mM) for 48h. Glutamate partially phenocopied the proliferation-promoting effect of the astrocytes. *Trp53*-null vs *Trp53*-WT cells cultured in medium supplemented with 75mM glutamate, shown as confluence over starting point, two-sided Student's t-test p=0.02 (at 36h), p=0.04 (at 48h). n=3.

**F,** Western blot protein quantification of SCD1 and FASN in isogenic *Trp53*-WT and *Trp53*-null cells cultured in SFM with or without glutamate (75mM). ACM exposure significantly elevated SCD1 and FASN protein levels in *Trp53*-null cells.

**G,H,** Quantification of SCD1 (**G**) and FASN (**H**) protein levels from three separate western blot experiments. Glutamate strongly stimulated SCD1 and FASN protein expression, specifically in *Trp53*-null cells. The effect of glutamate on SCD1 protein expression was significantly stronger in *Trp53*-null cells in comparison to *Trp53*-WT cells. Protein levels are shown as fold-change of glutamate-containing vs. SF medium. One-sided Student's t-test p=0.005 and $p=5*10^{-4}$ for SCD1 and FASN, respectively. *Trp53*-WT n=2, *Trp53*-null n=2.

**I,** Gene set enrichment analysis (GSEA) comparing the gene expression signatures of glutamate transport between primary human BCs and their matched brain metastases (GSE125989). Gene expression signatures associated with glutamate uptake are over-expressed in BM.

‘GOBP_L_GLUTAMATE_TRANSMEMBRANE_TRANSPORT’ p=0.001, ‘GOBP_LGLUTAMATE_IMPORT_ACROSS_PLASMA_MEMBRANE $p=6*10^{-4}$. n=16 tumor samples and their matched metastases. Also see Table S8.

**J,** Comparison of the mRNA expression levels of three glutamates receptors, *SLC1A3*, *SLC1A5* and *SLC1A6* between *TP53*-WT and *TP53*-null human primary breast tumors (based on TCGA data). Shown are the p-values of the over-expression.

**K**, qPCR-based analysis of *SLC1A5* mRNA expression in *Trp53*-WT and *Trp53*-null BC EMT6 cells cultured in ACM. *SLC1A5* expression was significantly higher in *Trp53*-null cells. Shown as fold-change increase in ACM relative to SFM. One-sided Student’s t-test p=0.04. *Trp53*-WT n=3, *Trp53*-null n=3.

Fig.S21

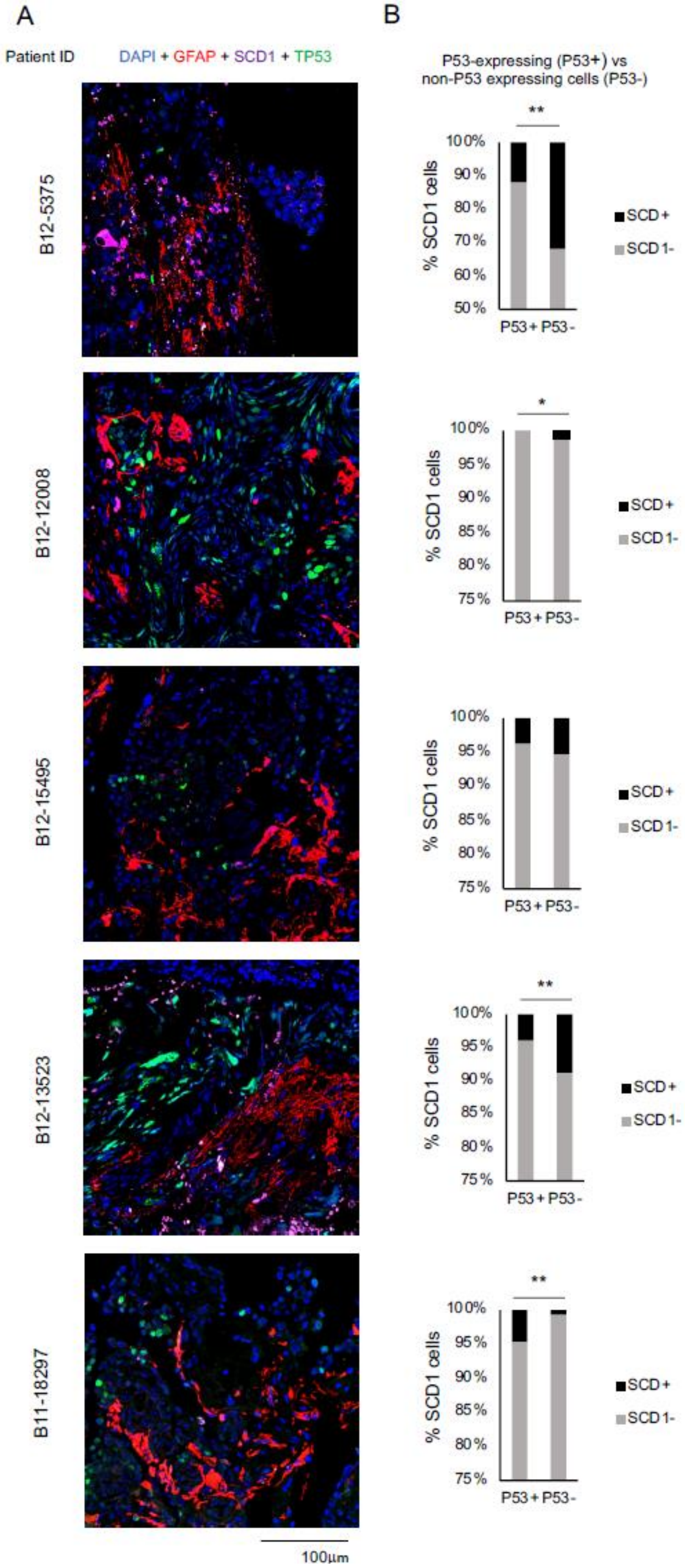
A
Patient ID
DAPI + GFAP + SCD1 + TP53
B12-5375
B12-12008
B12-15495
B12-13523
B11-18297
100µm
B
P53-expressing (P53+) vs
non-P53 expressing cells (P53-)
**
*
% SCD1 cells
100%
90%
80%
70%
60%
50%
95%
85%
75%
P53+ P53-
SCD+
SCD1-

**Fig. S21: *P53* inactivation is associated with SCD1 protein expression in BCBM from human patients (related to Fig. 5).**

**A,** Representative immunofluorescence images of human brain metastases from five BC patients, stained with antibodies against TP53 (green), SCD1 (purple), and GFAP (red), as well as with DAPI nuclear staining (blue). Data source: Iris Barshack.

**B,** Comparison of the percentage of SCD1-positive cells between p53-expressing and non-p53-expressing tumor cells. p53-negative cells were significantly enriched for SCD1 expression in 4 out of 5 metastases. Fisher's exact test, patient ID: B12-5375 $p=0.004$, B12-12008 $p=0.04$, B12-15495 $p=0.42$, B12-13523 $p=0.003$, B11-18297 $p=0.002$.

Fig.S22

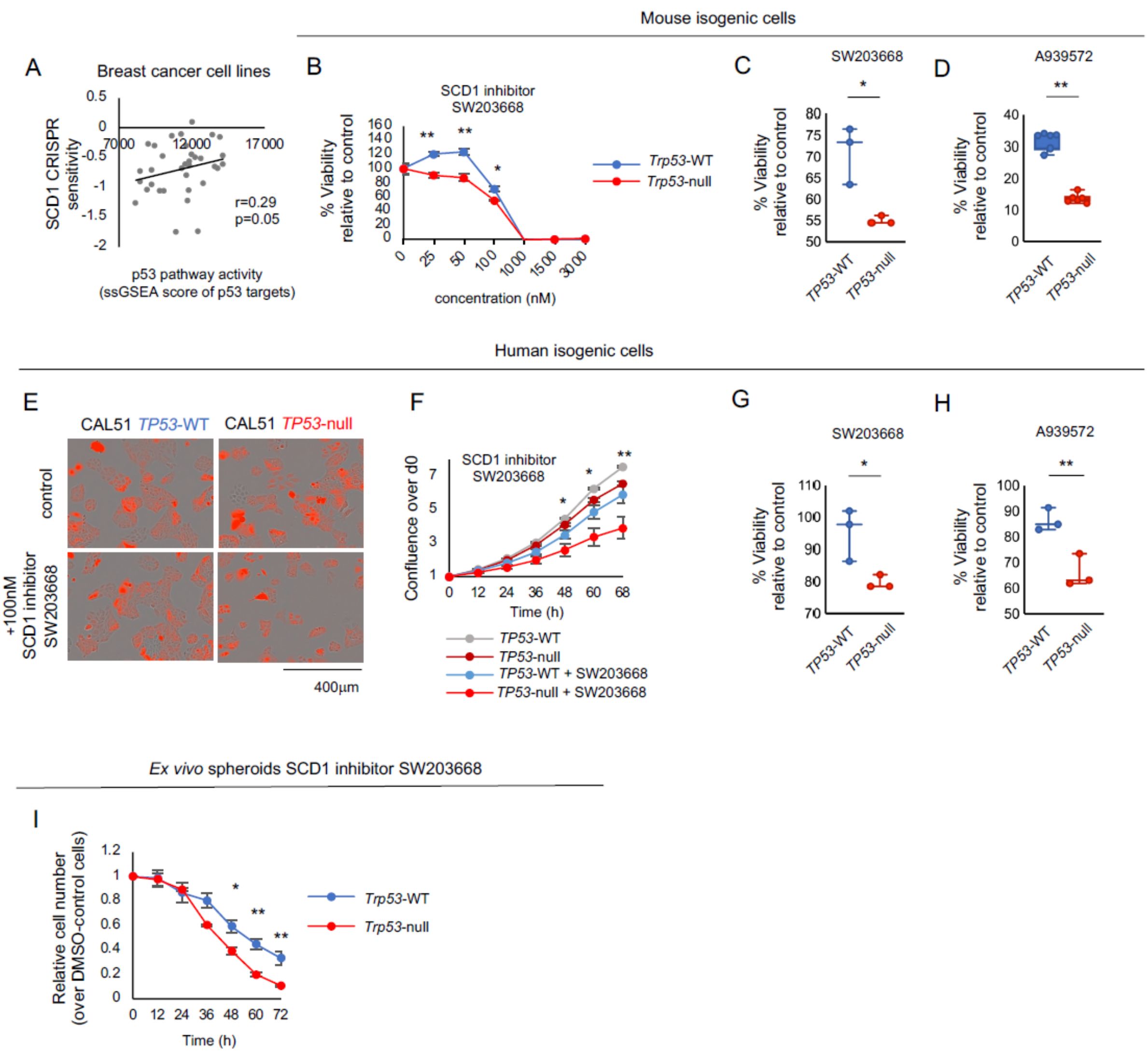

Mouse isogenic cells
A
Breast cancer cell lines
SCD1 CRISPR sensitivity
p53 pathway activity (ssGSEA score of p53 targets)
r=0.29
p=0.05
B
SCD1 inhibitor SW203668
% Viability relative to control
concentration (nM)
Trp53-WT
Trp53-null
C
SW203668
D
A939572
TP53-WT
TP53-null
Human isogenic cells
E
CAL51 TP53-WT
CAL51 TP53-null
control
+100nM SCD1 inhibitor SW203668
400µm
F
SCD1 inhibitor SW203668
Confluence over d0
Time (h)
TP53-WT
TP53-null
TP53-WT + SW203668
TP53-null + SW203668
G
SW203668
H
A939572
Ex vivo spheroids SCD1 inhibitor SW203668
I
Relative cell number (over DMSO-control cells)
Time (h)
Trp53-WT
Trp53-null

**Fig. S22**: ***P53* inactivation renders cells more sensitive to SCD1 inhibition (related to Fig. 5).**

**A,** The correlation between the p53 pathway activity score and the sensitivity of human BC cell lines to CRISPR/Cas9-mediated knockout of *SCD1*. Low p53 pathway activity is associated with high sensitivity to SCD1 inhibition (negative KO values represent increased sensitivity). p53 pathway activity was evaluated by calculating an ssGSEA score for the MSigDBgene signature 'KANNAN_p53_targets_DN'. Spearman's rho=0.29, p= 0.05 (one-tailed). BC cell lines n=34.

**B,** Live imaging-based dose response curves of *Trp53*-WT and *Trp53*-null EMT6 BC cells treated with the SCD1 inhibitor SW203668 or with control-DMSO for 48h. *Trp53*-null cells were significantly more sensitive to SCD1 inhibition than *Trp53*-WT controls. Two-sided Student's t-test p=0.01 for 25nM, p=0.004 for 50nM, p=0.02 for 100nM. *Trp53*-WT n=3, *Trp53*-null n=3.

**C,** Comparison of the relative viability of *Trp53*-WT and *Trp53*-null EMT6 cells to 48h of exposure to the SCD1 inhibitor SW203668 (100nM), quantified by the MTT assay. *Trp53*-null cells are significantly more sensitive to the drug. Two-sided Student's t-test p=0.02; *Trp53*-WT n=3, *Trp53*-null n=3.

**D,** Comparison of the relative viability of *Trp53*-WT and *Trp53*-null EMT6 cells to 48h of exposure to a second SCD1 inhibitor, A939572 (75nM), quantified by the MTT assay. *Trp53*-null cells are significantly more sensitive to the drug. Two-sided Student's t-test p=0.002, T*rp53*-WT n=3, *Trp53*-null n=3.

**E,** Representative images of mCherry-labeled *TP53*-WT and *TP53*-null human CAL51 BC cells treated with the SCD1 inhibitor SW203668 (100nM) or with control-DMSO for 48h. *TP53*-null cells were much more sensitive to the drug treatment.

**F,** Live imaging-based proliferation curves of *TP53*-WT and *TP53*-null human CAL51 BC cells treated with the SCD1 inhibitor SW203668 (100nM) or with control-DMSO for 72h. *TP53*-null cells were significantly more sensitive to SCD1 inhibition than *TP53*-WT controls. One-sided Student's t-test significant at indicated time points: 48h p=0.03, 60h p=0.02, 68h p=0.01. For each condition, n=3, 15 fields imaged.

**G,** Comparison of the relative viability of *TP53*-WT and *TP53*-null CAL51 cells to 72h of exposure to the SCD1 inhibitor SW203668 (100nM), quantified by the MTT assay. *TP53*-null cells are significantly more sensitive to the drug. One-sided Student's t-test p=0.02; *TP53*-WT n=3, *TP53*-null n=3.

**H,** Comparison of the relative viability of *TP53*-WT and *TP53*-null CAL51 cells to 72h of exposure to another SCD1 inhibitor, A939572 (75nM), quantified by the MTT assay. *TP53*-null cells are significantly more sensitive to the drug. One-sided Student's t-test p=0.005; *TP53*-WT n=3, *TP53*-null n=3.

**I**, Comparison of the effect of treatment with the SCD1 inhibitor SW203668 (3μM) on the growth of *Trp53*-WT and *Trp53*-null tumor derived 3D spheroids. *Trp53*-null spheroids are more sensitive to SCD1 inhibition than their *Trp53*-WT controls. Response is shown as confluence relative to DMSO-treated control cells. One-sided Student's t-test p=0.02 at 36h, p=0.02 at 48h, p=0.003 at 60hr, p=0.01 at 72h. *Trp53*-WT n=3, *Trp53*-null n=3.

Fig.S23

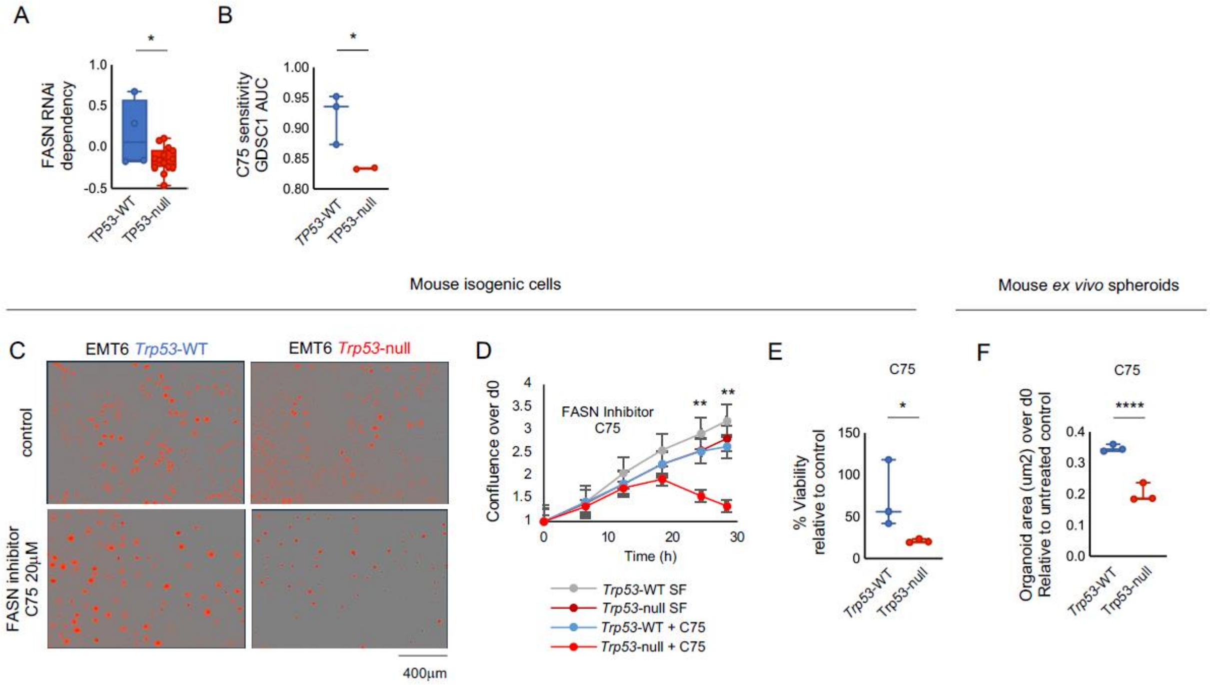

A
*
FASN RNAi dependency
1.0
0.5
0.0
-0.5
TP53-WT
TP53-null
B
*
C75 sensitivity GDSC1 AUC
1.00
0.95
0.90
0.85
0.80
TP53-WT
TP53-null
Mouse isogenic cells
Mouse ex vivo spheroids
C
EMT6 Trp53-WT
EMT6 Trp53-null
control
FASN inhibitor C75 20μM
400μm
D
Confluence over d0
FASN Inhibitor C75
**
**
Time (h)
Trp53-WT SF
Trp53-null SF
Trp53-WT + C75
Trp53-null + C75
E
C75
*
% Viability relative to control
150
100
50
0
Trp53-WT
Trp53-null
F
C75
****
Organoid area (um2) over d0 Relative to untreated control
0.4
0.3
0.2
0.1
0.0
Trp53-WT
Trp53-null

**Fig. S23**: ***P53*-null cancer cells are more sensitive to FASN inhibition that p53-WT controls (related to Fig.5)**

**A,** Comparison of the sensitivity of *TP53*-WT and *TP53*-null human BC cell lines to RNAi-mediated knockdown of *FASN*. *TP53*-null cells are more sensitive to the genetic inhibition of *FASN* than *TP53*-WT cells. One-sided Student's t-test p=0.02. CCLE BC cell lines: *TP53*-WT n=4, *TP53*-null n=14.

**B,** Comparison of the sensitivity of *TP53*-WT and *TP53*-null human BC cell lines to the FASN inhibitor C75. *TP53*-null cell lines are more sensitive to the chemical inhibition of FASN than *TP53*-WT cells. Data are shown as AUC values, obtained from the GDSC screen (*146*). One-tailed Student's t-test p=0.03. CCLE BC cell lines: *TP53*-WT n=3, *TP53*-null n=2.

**C,** Representative images of mCherry-labeled *Trp53*-WT and *Trp53*-null mouse EMT6 BC cells treated with the FASN inhibitor C75 (20µM) or with control-DMSO for 28h. *TP53*-null cells were much more sensitive to the drug treatment.

**D,** Live imaging-based proliferation curves of *Trp53*-WT and *Trp53*-null mouse EMT6 BC cells treated with the FASN inhibitor C75 (20µM) or with control-DMSO for 28h. After 18h of drug exposure, a reduction of fluorescent signal was observed in *Trp53*-null cells, whereas DMSO-control *Trp53*-null cells and C75-treated *Trp53*-WT cells kept proliferating. C75-treated *Trp53*-WT vs *Trp53*-null cells relative to respective SFM controls, one-sided Student's t-test. p=0.01 at 24h, p=0.005 at 28h.

**E,** Comparison of the sensitivity of *Trp53*-WT and *Trp53*-null mouse BC cells in response to the FASN inhibitor C75 (20µM). Viability by MTT shown relative to respective SFM controls, one-sided Student's t-test, p=0.05.

**F,** Live imaging-based comparison of spheroid size (µ$m^2$) between *Trp53*-WT and *Trp53*-null spheroids treated with the FASN inhibitor C75 (20µM) for 48h, relative to DMSO-control conditions. One-sided Student's t-test p=$2*10^{-4}$. *Trp53*-WT n=3, *Trp53*-null n=3.

**Legends to Supplementary Tables:**

**Table S1** GSEA of brain metastases compared to metastases in other organs (p53 signatures, 7 tabs).

Shown are gene sets that are differentially expressed between brain metastases and metastases in other organs. Unbiased GSEA and ssGSEA of 3 datasets (Tab 1-4: GSE14017, Tab 5: GSE14018 and Tab 6-7: GSE245414) using the MSigDB 'Oncogenic' gene sets. Highlighted in red are p53 signatures that indicate lower p53 activity in BM. Gene sets with $p<0.05$ are listed. Q-values for ssGSEA results were computed using the Benjamini–Hochberg method.

**Table S2** GSEA of primary breast tumors that later metastasized to the brain vs those that metastasized to other organs (p53 signatures, 7 tabs).

Tabs 1-5: Unbiased GSEA/ssGSEA comparing primary BC that would later metastasize to the brain vs. primary tumors that would later metastasize to other metastatic sites, shown for 2 datasets (MBP and GSE12276). Tab 6: GSEA of human breast cancer cell lines, comparing cell lines with high brain metastatic potential (top quartile) vs. those with low brain metastatic potential (bottom quartile). Tab 7: GSEA of human carcinoma cell lines, with high brain metastatic potential (top quartile) vs. those with low brain metastatic potential (bottom quartile). Highlighted in red are p53 signatures that indicate decreased p53 activity in tumors with high brain metastatic capacity. Q-values for ssGSEA results were computed using the Benjamini–Hochberg method.

**Table S3** ssGSEA of *TP53*-null vs *TP53*-WT tumors (BM signatures, 3 tabs)

Unbiased GSEA of primary BC with vs. without *TP53* alterations (*TP53* copy number loss or mutation). Shown are the top100 significant results, sorted by effect size, from the analysis of 3 independent data sets (METABRIC, TCGA, MBP). brain metastasis/relapse signatures, indicating increased metastatic potential to the brain in *TP*53-null BC, are highlighted in red. Q-values for ssGSEA results were computed using the Benjamini–Hochberg method.

**Table S4** GSEA of primary breast tumors that later metastasized to the brain vs those that metastasized to other organs (BM signatures, 2 tabs). Validation of gene signatures associated with brain metastasis/relapse by GSEA comparing: (a) primary BC that would metastasize to the brain vs. BC that would metastasize to other sites (Tab 1); (b) human breast cancer cell lines, sorted by brain metastatic potential (top quartile vs bottom quartile). Shown are GSEA signatures sorted by effect size (top50). The gene signatures SMID_BREAST_CANCER_RELAPSE_IN_BRAIN, -_UP and -_DN are highlighted in red. Q-values for ssGSEA results were computed using the Benjamini–Hochberg method.

**Table S5** IHC analysis of 17 BCBM clinical samples
Assessment of P53, P21, SCD1 and GFAP immunostainings on BCBM clinical samples. For details, please see Methods section.

**Table S6** List of the GSEA signatures related to Adipogenesis, FA synthesis, FA uptake and FA activation, as presented in Fig. 4A-D and Fig. 16A-D (8 tabs).

**Table S7** GSEA of BM compared to metastases in other organs (Lipid metabolism signatures, 3 tabs). Differential gene expression analysis between BCBM and metastases in other organs. Unbiased GSEA and ssGSEA of 3 datasets (GSE14017, GSE14018 and GSE245414) using the 'Reactome', 'Kegg' and 'GOMF' gene sets. Highlighted in red are signatures that indicate increased fatty acid biosynthesis and transport in BM. Q-values for ssGSEA results were computed using the Benjamini–Hochberg method.

**Table S8** ssGSEA of BCBM vs. their matched primary tumors (Lipid metabolism signatures, 2 tabs). Differential expression analysis between primary BC and their matching BM focusing on the 'Reactome', 'GOBP' and 'GOMF' gene sets. Highlighted in red are signatures that indicate increased fatty acid synthesis/transport and glutamate transport in BM. Q-values for ssGSEA results were computed using the Benjamini–Hochberg method.

**Table S9** GSEA of BCs that later metastasized to brain vs those that metastasized to other sites (Lipid metabolism signatures, 2 tabs). Differential gene expression analyses of primary BC that would metastasize to the brain vs. those that would metastasize to other organs (GSE12276) using 'Reactome' and 'GO' gene sets (Tab 1). GSEA of human breast cancer cell lines, the top and bottom quartiles of brain metastatic potential (Tab 2). Highlighted in red are FA biosynthesis and transport signatures elevated in BC with brain metastatic capacity. Q-values for ssGSEA results were computed using the Benjamini–Hochberg method.

**Table S10** ssGSEA of *TP53*-null vs *TP53*-WT tumors (Lipid metabolism signatures, 2 tabs). Unbiased gene expression analysis of primary BC with vs. without *TP53* alterations (*TP53* copy number loss or mutation). Shown are the 'Reactome', 'Kegg', 'GOBP' and 'GOMF' gene sets. Results are shown for two independent data sets (METABRIC, MBC). Highlighted in red are elevated FA biosynthesis signatures in BC with *TP53* alterations.

**Table S11** ssGSEA of CAL51 *TP53*-KO vs *TP53*-WT (Lipid metabolism signatures, 1 tab) Unbiased ssGSEA of CAL51 BC cells subjected to *TP53* KO compared to their controls. Highlighted in red are FA biosynthesis and FA transport signatures that are elevated in *TP53* KO. Q-values for ssGSEA results were computed using the Benjamini–Hochberg method.

**Table S12** Metabolic profiling of ACM by LC-MS. Comparison of lipids and polar metabolites in ACM and serum-free base medium in triplicates. Shown as normalized data to internal standards and volume.

**Table S13** Analysis of FA content in *Trp53*-null and *Trp53*-WT EMT6 cells grown in ACM using gas chromatography. Comparison of FA content as mole % in *Trp53*-null and *Trp53*-WT cells that were cultured for 20h in ACM or SFM.

**Table S14** Comparison of gene essentiality (RNAi sensitivity) between *TP53*-null and *TP53*-WT BC cell lines. The top 500 differential RNAi sensitivities between *TP53*-null and *TP53*-WT were subjected to GSEA. Shown are the top 25 most significantly enriched pathways (MsigDB 'Hallmark' and C2 'Canonical Pathways' gene sets).

**Table S15** Reagents List: detailed information on reagents including CRISPR targeting sequences, primer sequences, antibodies and drugs.
